# Effects of microRNA-mediated negative feedback on gene expression noise

**DOI:** 10.1101/2022.11.09.515756

**Authors:** Raunak Adhikary, Arnab Roy, Mohit Kumar Jolly, Dipjyoti Das

## Abstract

Micro-RNAs (miRNAs) are small non-coding RNAs that regulate gene expression post-transcriptionally in eukaryotes by binding with target mRNAs and preventing translation. miRNA-mediated feedback motifs are ubiquitous in various genetic networks which control cellular decision-making. A key question is how such a feedback mechanism may affect gene expression noise. To answer this, we have developed a mathematical model to study the effects of a miRNA-dependent negative feedback loop on mean expression and noise in target mRNAs. Combining analytics and simulations, we show the existence of an expression threshold demarcating repressed and expressed regimes in agreement with earlier studies. The steady-state mRNA distributions are bimodal near the threshold, where copy numbers of mRNAs and miRNAs exhibit enhanced anticorrelated fluctuations. Moreover, variation of negative-feedback strength shifts the threshold locations and modulates the noise profiles. Notably, the miRNA-mRNA binding affinity and feedback strength collectively shape the bimodality. We also compare our model with a direct auto-repression motif, where a gene produces its own repressor. Auto-repression fails to produce bimodal mRNA distributions as found in miRNA-based indirect repression, suggesting the crucial role of miRNAs in creating phenotypic diversity. Together, we demonstrate how miRNA-dependent negative feedback modifies the expression threshold and leads to a broader parameter regime of bimodality compared to the no-feedback case.

## 1 Introduction

In order to survive, cells actively regulate the processes of transcription (synthesis of mRNAs from genes) and translation (synthesis of proteins from mRNAs) in response to environmental cues [91]. However, since biological molecules are present in low numbers in a cell and due to the heterogeneity of environmental factors, large cell-to-cell fluctuations can exist in mRNA and protein copy numbers. Thus, transcription and translation are inherently stochastic processes leading to the heterogeneity of gene products in genetically identical cell populations – this phenomenon is known as ‘gene expression noise’ [40, 43, 67, 70]. Understanding how gene expression noise is regulated has emerged as an important research area in molecular biology [20, 59, 66, 70, 73, 86].

An effective molecular mechanism for gene regulation is via the binding and unbinding of diverse transcription factors (TFs) at specific sites close to a gene’s promoter. Such control on the *promoter architecture* through TF-binding can regulate expression noise, as found in predictive theoretical studies [16, 18, 70–72, 74, 76] and experiments [14, 22, 32, 39, 65, 85]. Phenotypic consequences of expression variability were also investigated, and it was found that gene expression noise can control cell fate switching and have survival benefits in varying environments [6, 46, 55, 87].

Post-transcriptional regulation by short noncoding RNAs is another common mechanism for controlling gene expression in both prokaryotes and eukaryotes. For instance, non-coding small RNAs (sRNAs) in bacteria bind with target mRNAs to facilitate mRNA degradation [28–30, 45, 50, 57]. Similarly, eukaryotic microRNAs (miRNAs) are about 22 nucleotide-long noncoding RNAs which interact with mRNAs in a sequence-dependent manner. After maturing into RNA-Induced Silencing Complexes (RISC), miRNAs can bind to the 3’UTR of the target mRNAs and then either degrade the transcript or inhibit their translation [9, 91]. Such post-transcriptional control by miRNAs affects various biological processes, including animal development, stabilization of gene expression as a stress response, and inhibition of cancer metastasis. [12, 48, 51, 52, 78].

Previous quantitative studies have suggested that the interplay between mRNAs and miRNAs (or sRNAs) occurs via a molecular titration-like mechanism [50, 58]. However, the mode of interaction between miRNAs and mRNAs is still debated. Some studies advocate an almost catalytic interaction [3, 37], and some suggest stoichiometric interaction [5, 41, 63]. During catalytic interaction, miRNAs recycle back, maintaining a pool [9, 10, 17], while miRNAs and mRNAs could destroy each other through stoichiometric interaction. On the other hand, prokaryotic sRNAs mainly act stoichiometrically on their target mRNAs [31, 90]. Nevertheless, both eukaryotic miRNA and prokaryotic sRNA can produce a threshold-like behavior at the mean expression level, with a sharp demarcation of low and high expression regimes, due to a competitive titration between miRNAs (or sRNAs) and their targets [50, 57, 58, 62]. Though some theoretical studies have focused on miRNA-mediated gene expression noise [7, 17, 62, 75], there are still some differences in opinions. For instance, one study [34] claimed that catalytic interaction reduces gene expression noise compared to stoichiometric interaction involving miRNAs, but another study [62] asserted that both stoichiometric and catalytic interactions produce qualitatively similar noise in certain parameter regimes. Moreover, additional mechanisms involving miRNAs can complicate the noise regulation in genetic networks.

An exciting aspect of miRNA-mediated regulation is that one class of miRNAs can affect many distinct genes, leading to the *competing endogenous RNA* (ceRNA) hypothesis, which suggests regulating the expression of one gene by affecting the transcription of another gene that shares the same pool of miRNAs. Theoretical and experimental works based on the ceRNA hypothesis have shown that multiple target mRNAs influence each other nonlinearly through a common pool of miRNAs, producing high variability in expression [2, 9, 10, 24, 83]. As reported in sRNA-dependent bimodal gene expression [49], miRNA-dependent bimodal expression in ceRNA networks was also predicted theoretically [9, 62] and later shown in experiments [10, 17].

Notably, miRNA-dependent feedback motifs are ubiquitous in regulatory networks [88], which dictate diverse physiological and developmental processes, including cell cycle control [21], cancer cell proliferation, chemo-resistance, angiogenesis [1, 15, 92], and host-HIV interaction [25]. Feedback involving miRNAs establishes a cross-talk between post-transcriptional and transcriptional layers to precisely regulate some transitions of biological states. For example, the network controlling epithelial-mesenchymal transition during cancer progression involves miRNA-mediated feedback loops. [38, 54]. Another well-known example is the differentiation of precursor cells into dopamine neurons in the midbrain, which is governed by a negative feedback loop involving the miRNA, miR-133b, and the TF, *pitx3* [44].

Despite mounting evidence that miRNAs are critical down-regulators of gene expression, the theoretical understanding of miRNA-based feedback loops in regulating gene expression noise is still incomplete. Several theoretical studies have focused on the miRNA-based feedback loops, albeit mainly from a deterministic standpoint focusing on the mean level (reviewed in [47]). Among different feedback motifs, those involving negative feedback are of particular interest because of their capacity to buffer gene expression noise [8, 47, 79]. Using a deterministic mean-field approach, Zhou et al. have studied a miRNA-based Single Negative Feedback Loop (SNFL), where a TF promotes the expression of a miRNA that, in turn, inhibits the TF expression [93]. Another study focused on a similar SNFL to investigate both the mean and noise of expression, but the interactions between the miRNA-producing gene and TFs were modeled in a coarse-grained way [89].

In this paper, we developed a detailed model of the miRNA-mediated single-negative-feedback loop and investigated the properties of mean and variance of expression in the steady-state. We found that the steady-state mean mRNA shows a threshold-linear behavior, i.e., the mean is zero below a threshold transcription rate, and the mean increases almost linearly above the threshold. This observation agrees with earlier studies without any feedback [58, 62]. Such a threshold stems from a competitive titration between miRNAs and mRNAs, and the threshold point corresponds to the situation when the numbers of mRNAs and miRNAs are comparable. Our mean-field calculation also provided an analytical expression for the threshold, which shows that the threshold point crucially depends on miRNA’s catalytic interaction and feedback strength. Notably, the noise in mRNA copy numbers also peaked in the threshold’s vicinity, suggesting the genetic motif’s high sensitivity at this region. We also found bimodal mRNA distributions near the threshold, unlike a similar theoretical study on miRNA-mediated negative feedback [89]. Such bimodal distributions correspond to large anticorrelated fluctuation in miRNA and mRNA numbers, suggesting stochastic switching of mRNAs between two states: miRNA-bound (repressing the target mRNAs) and miRNA-unbound (expressing the mRNAs). We summarized the emergence of bimodality in a 2D phase diagram, signifying that the bimodality is mainly controlled by the miRNA-mRNA binding strength and the negative feedback strength.

Finally, we compared the outputs of a miRNA-mediated negative feedback loop with a negative auto-regulatory loop since auto-regulatory motifs are essential components in many genetic networks often seen in various biological contexts [23, 77, 84]. An auto-regulatory motif, where a protein expressed from a gene acts as a TF for the same gene, can be either positive or negative, depending on whether the TF enhances or suppresses its own expression. A positive auto-regulation produces bi-stable expression, and subsequent bimodal protein distribution, enhancing the expression noise [23, 35, 36, 42, 64, 82]; whereas negative auto-regulation can suppress noise [19, 61, 80, 81]. However, some recent theoretical studies created a difference in opinion using different assumptions, and showed that negative auto-regulation can also enhance noise in certain situations [53, 56].

When we compared our model of miRNA-based negative feedback with an auto-repression, at first glance, both circuits exhibited qualitatively similar behavior of the mean and noise as functions of respective feedback strengths. However, we found that negative feedback involving miRNAs produces bimodal mRNA distribution by amplifying noise, but auto-repression leads to only bell-shaped mRNA distributions. Our study thus highlights the importance of miRNA-based negative feedback in producing phenotypic diversity.

## 2 Model

We adapted a published model [9] of miRNA-mRNA interaction (without any feedback) to describe the miRNA-mediated negative feedback loop (see Fig. 1). In the model, the mRNA molecules are synthesized from an mRNA-coding gene at a constant rate *k*_*r*_, and then protein molecules are produced from the mRNAs at a rate *k*_*p*_. The mRNAs and proteins degrade with rates *g*_*r*_ and *g*_*p*_, respectively. The translation of mRNAs into proteins is inhibited post-transcriptionally by the miRNA molecules produced from a miRNA-coding gene at a basal rate 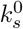. The miRNAs degrade at a rate *g*_*s*_. The proteins can activate the miRNA-coding gene and increase the miRNA synthesis rate, establishing a negative feedback on gene expression. Thus, the miRNA-coding gene can be in two distinct states based on the miRNA synthesis rate: miRNAs are produced either at a basal rate 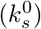 or at an enhanced rate 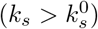 (see Fig. 1). These states are termed ‘OFF’ (basal) and ‘ON’ (enhanced), respectively. The protein binds at a rate *k*_*act*_ to the miRNA-coding gene making it switch from OFF (basal) to ON (enhanced) state. Thus, *k*_*act*_ is termed as the activation rate of the miRNA-gene. On the other hand, the switch from the ON to OFF state occurs when the bound protein dissociates from the miRNA-coding gene (rate, *k*_*deact*_). Note that the non-dimensional ratio *β* = *k*_*act*_*/k*_*deact*_ can be considered as the negative feedback strength. Since the post-transcriptional regulation of mRNAs takes place via direct association and dissociation between mRNAs and miRNAs, we consider the binding (rate *k*_+_) and unbinding (rate *k*_−_) processes between an mRNA and a miRNA, thereby forming an mRNA-miRNA complex. The mRNA-miRNA complex may disintegrate in two ways: (i) either the complex fully degrades at a rate *αγ*, simultaneously destroying both the mRNA and miRNA molecules or (ii) only the bound mRNA is degraded (with a rate (1 − *α*)*γ*) and the miRNA is recycled back. Here, *γ* denote a complex disintegration rate, and *α* (0 ≤ *α* ≤ 1) is the ‘catalyticity parameter’. Note that *α* determines the fraction of miRNAs recycled after dissociating from their target mRNAs. All the processes described here can be expressed in terms of a set of chemical reactions (see Eq. S0.1-S0.11 in SI).

**Fig 1.**
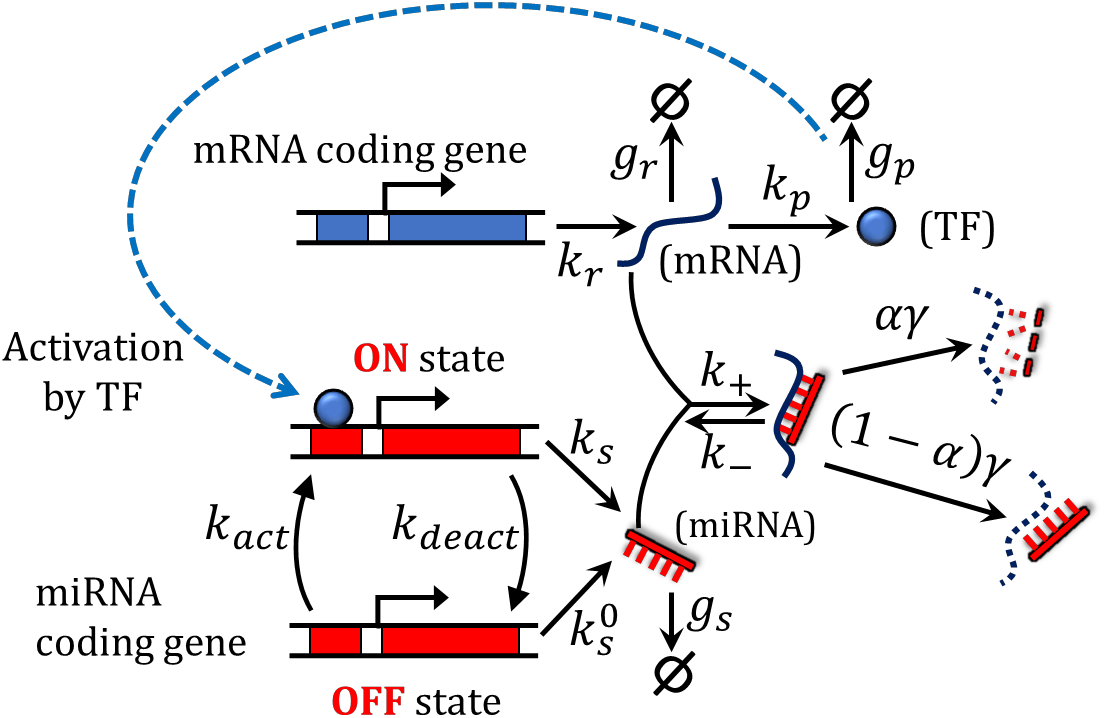
Schematic diagram of miRNA-mediated negative feedback showing various kinetic processes with corresponding rates (see Model and Table 1). The miRNAs bind with the target mRNAs to form miRNA-mRNA complexes. The complexes degrade with a rate *αγ*, or the miRNAs act catalytically, degrading only the mRNAs (rate (1 − *α*)*γ*). The parameter *α* (0 ≤ *α* ≤ 1; called catalyticity parameter) determines the fraction of miRNAs that are degraded when they are bound to mRNAs. The mRNAs produce a transcription factor (TF) that activates a miRNA coding gene and enhances its transcription rate more than the basal rate (in general,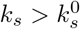). Thus, the miRNA-coding gene toggles between ON (activated) and OFF (basal) states. Here, *β* = *k*_*act*_*/k*_*deact*_ can be considered as the ‘feedback strength’, while *g* = *k*_+_*γ/*(*k*_−_ + *γ*) is an ‘effective association rate’ between miRNAs and mRNAs.

Based on the model, we described the time-evolution of four stochastic integer-valued variables, namely the instantaneous numbers of mRNAs, proteins, miRNAs, and mRNA-miRNA complexes (denoted by *r*(*t*), *p*(*t*), *s*(*t*), and *c*(*t*), respectively). Using the framework of Stochastic Processes, we write down below the Master equation governing the joint probability distribution, *P*_*r,p,s,c*_(*t*), defined as the probability of observing *r, p, s*, and *c* number of mRNAs, proteins, miRNAs and complexes, respectively, at a time *t*.

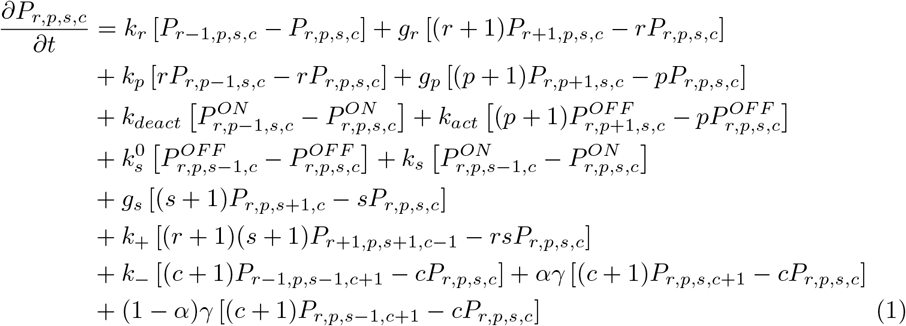

**Table 1.** Description of model parameters and their values taken from literature [9, 62]

| Symbol | Description | Value |
| --- | --- | --- |
| $k_r$ | mRNA synthesis rate | varied ($0.0s^{-1}$ - $1.0s^{-1}$) |
| $g_r$ | mRNA degradation rate | $0.0004s^{-1}$ |
| $k_p$ | protein synthesis rate | $0.4s^{-1}$ |
| $g_p$ | protein degradation rate | $0.0005s^{-1}$ |
| $k_s$ | miRNA synthesis rate varied in the activated (ON) state | varied ($0.0s^{-1}$ - $1.0s^{-1}$) |
| $k_s^0$ | miRNA synthesis rate in the basal (OFF) state | $0.05s^{-1}$ |
| $g_s$ | miRNA degradation rate | $0.0003s^{-1}$ |
| $k_+$ | mRNA-miRNA association rate | varied ($0.0001s^{-1}$-$1.0s^{-1}$) |
| $k_-$ | mRNA-miRNA dissociation rate | $0.0036s^{-1}$ |
| $\gamma$ | mRNA-miRNA complex degradation rate | $0.0004s^{-1}$ |
| $\alpha$ | Cataliticity parameter | varied (0 - 1) |
| $k_{act}$ | activation rate (OFF to ON) of miRNA coding gene | varied ($0.0s^{-1}$ - $0.01s^{-1}$) |
| $k_{deact}$ | deactivation rate (ON to OFF) of miRNA coding gene | $0.01s^{-1}$ |

The Master equation (Eq. 1) is constructed by accounting for the ‘gain’ and ‘loss’ terms corresponding to all possible changes in the model variables (*r, p, s*, or *c*). Here, *P*_*r,p,s,c*_(*t*) is the joint probability distribution irrespective of the miRNA-gene state (ON or OFF), and hence it can be broken into two 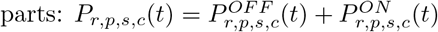 (corresponding to each states — see Eq. S1.1-S1.3 in SI).

We obtained the time-evolution equations for means and variances of our variables (i.e., first and second-order moments) from the Master equation described above(see SI, Eq S2.1-S2.29). However, the moment equations do not close themselves since each equation contains the next higher-order moment. Due to this challenge, we obtained approximate equations for the means and variances (by neglecting higher-order moments) that can be tackled analytically (Eq. S3.1-S3.5 and Eq. S6.1-S6.20). Alternatively, we performed exact stochastic simulations [27] (Gillespie simulations) of our model using biologically relevant kinetic rates (see Table 1) to obtain the moments of mRNA and miRNA distributions in the steady-state. Below we summarize the results.

## 3 Results

### 3.1 Feedback strength tunes the expression threshold exhibited by target mRNAs and enhances the noise around the threshold

Using the stochastic framework outlined before, we first investigated how the negative feedback affects the mean of mRNA copy numbers (ė*r*⟩) in the steady-state. Fig 2A shows the mean mRNA obtained from the simulations as a function of the mRNA transcription rate (*k*_*r*_) for different feedback strengths. Both in the absence and presence of the feedback, we found a threshold-like behavior of the mean — the miRNAs repress the target mRNAs making the mRNA mean zero up to a threshold transcription rate, and the mean increases almost linearly above the threshold. This behavior was already reported in previous studies of miRNA-based repression in the abscence of feedback [9, 58, 62]. However, when the feedback strength (*β* = *k*_*act*_*/k*_*deact*_) is varied (by changing the miRNA-gene activation rate, *k*_*act*_), the location of the threshold changes (Fig 2A). The mean mRNA sharply transits from low to high levels across the thresholds in the limits of no-feedback and very high feedback (see the curves for *β* = 0 and *β* = 1 in Fig 2A), while the sharpness of transitions around thresholds decreases for intermediate feedback strengths.

**Fig 2.**
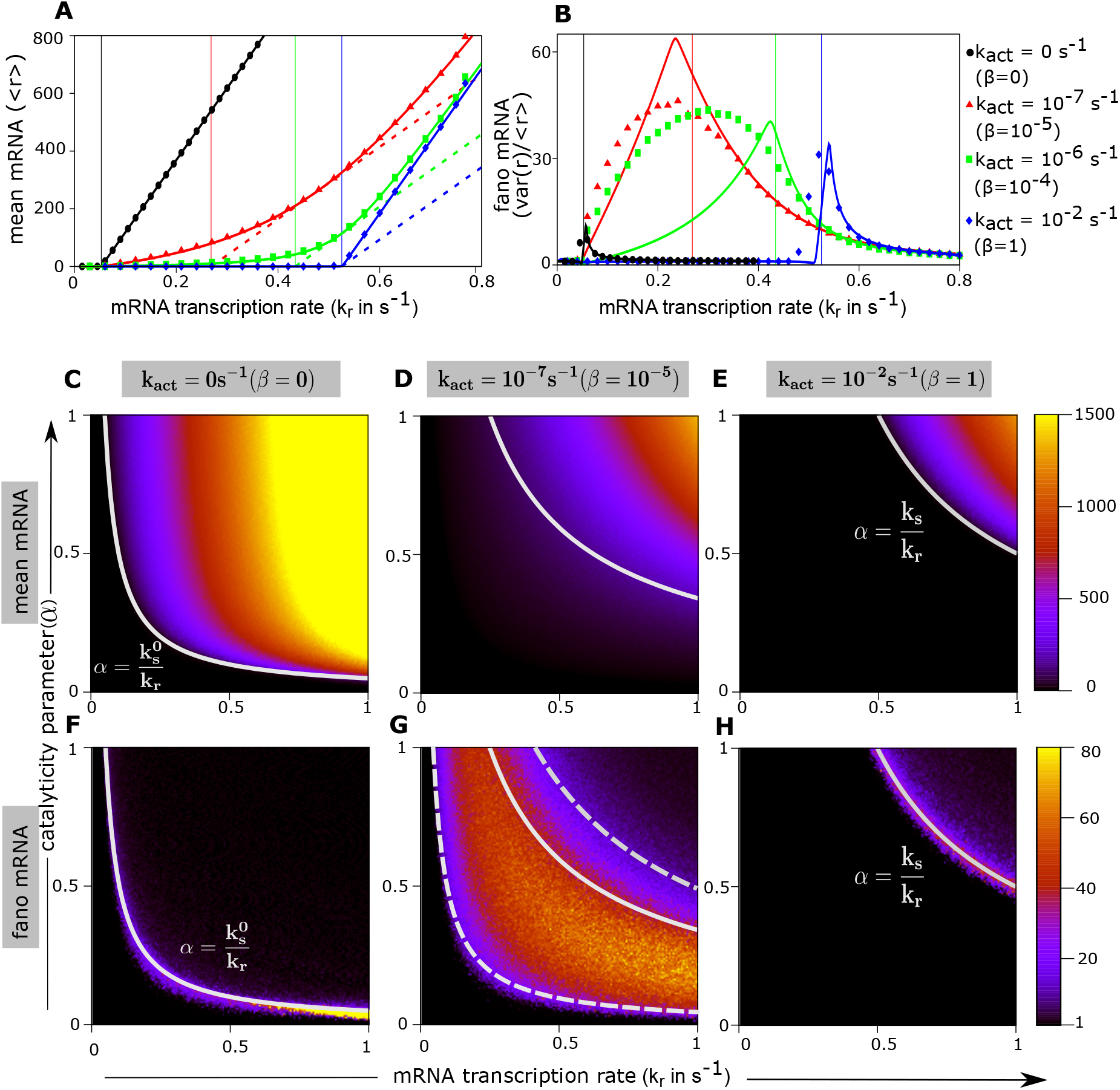
Varying negative feedback strength modifies the expression threshold and enhances mRNA noise around the threshold. (A-B) The steady-state mean mRNA and its Fano factor are shown against mRNA transcription rate for various feedback strengths (*k*_*act*_ was varied, while *k*_*deact*_ was kept fixed). In panels A and B, data points are from simulations, while solid curves represent analytical solutions of the approximate moment equations (see Eq. 2 in main text and Eq. S2.1-S2.9, S6.1-S6.20 in SI). Dashed slanted lines in panel A are the tangents to the mean mRNA curves (solid curves) at a point where the change in slopes are maximum — these tangents intersect the *k*_*r*_ axis at the threshold points. The vertical straight lines in A and B indicate respective threshold values (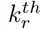 from Eq. 3, 5). In panel B, the Fano factors peak near the thresholds. (C-H) Heat plots show mRNA means (C-E) and Fano factors (F-H) for different *k*_*act*_ in the plane of catalyticity parameter (*α*) and mRNA transcription rate (*k*_*r*_). Color bars represent magnitudes of means and Fano factors. Analytical formula of 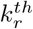 (Eq. 3, 5) yields respective separation-boundaries between low and high expression regimes (white curves in C-H). For the cases of no-feedback (F) and high feedback (H), the high noise regime coincides with the threshold boundaries. However, for low feedback strength (G), the high noise regime is broadened and lies between the threshold boundaries corresponding to the limiting cases of no-feedback and very high feedback (white dash curves in G re-plotted from panels F and H). For all panels, *k*_*s*_ = 0.5*s*^−1^, *k*_+_ = 1*s*^−1^; while *α* = 0.95 for panels A-B. Other parameters were from Table 1.

To obtain the thresholds in mRNA transcription rate analytically, we derived the time-evolution equations for the mean variables from the Master equation (Eq 1) under the mean-field approximation. The mean-field equations (MFE) are as below (see SI for details):

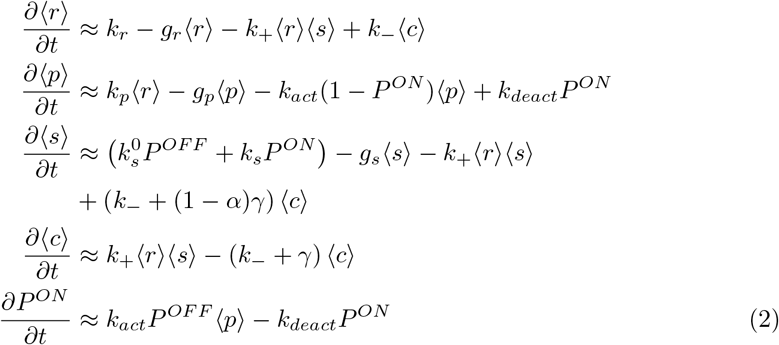

Here, the notation, ‘⟨ · · · ⟩’ denotes the average of respective variables, and *P*^*ON*^ is the probability of the miRNA gene to be in ON state. We also introduce an ‘effective’ mRNA-miRNA association rate defined as *g* = *k*_+_*γ/*(*k*_−_ + *γ*). Despite the mean-field approximation, the solutions of Eq. 2 matched with the exact simulation data (see the solid curves and data points in Fig. 2A).

The effect of negative feedback on the threshold behavior can be understood intuitively in the limit *g* → ∞ (i.e., strong binding of miRNAs to mRNAs). In this limit, solving Eq. 2, we can show:

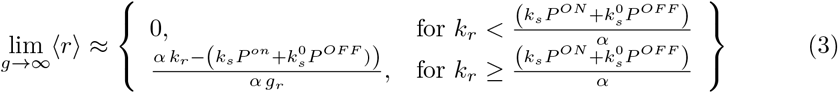

where,

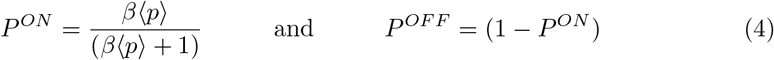

From Eq. 3, the threshold mRNA transcription rate is given by 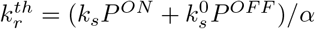 (under mean-field approximation), and this mainly depends on the state probabilities of the miRNA gene (*P*^*ON*^ and *P*^*OFF*^). The state probabilities, in turn, depend on the feedback strength (*β* = *k*_*act*_*/k*_*deact*_) and the mean protein level (⟨*p*⟩ ≈ (*k*_*p*_*/g*_*p*_) ⟨*r*⟩). Thus, when the feedback strength is varied, the threshold position shifts between two extreme limits: (i) In the absence of the feedback (*β* = 0), the threshold is 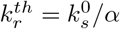 (as reported previously [9, 62], and (ii) in the limit of extremely high feedback (*β* → ∞), the threshold becomes 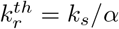.

However, we can derive a general expression of the threshold mRNA transcription rate for any nonzero feedback strength. To this end, we first solved the MFE (Eq. 2) to obtain ⟨*r*⟩ as an explicit function of *k*_*r*_, and then defined the threshold 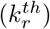 to be the intersection point of the *k*_*r*_-axis and the tangent to the mean mRNA curve drawn at a point where the change in slope is the maximum (see the dashed lines in Fig 2A, and details are in SI-Section S5.1). When the mRNA-miRNA effective association rate is high (*g* → ∞), the general *k*_*r*_ threshold is given by,

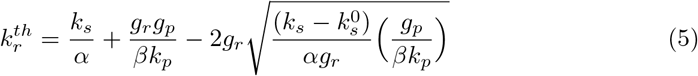

The expression of the threshold, 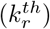, (Eq. 5) fairly locates the transition from low to high mean mRNA level (see the vertical lines in Fig 2A representing respective thresholds).

We next explored how the feedback strength affects the noise in mRNA copy number quantified by the mRNA Fano factor, which is the ratio of variance to the mean (*var*(*r*)*/* ⟨*r*⟩). As reported previously [9, 62], Fano factors showed peaks around the threshold points when plotted against the mRNA transcription rate (Fig. 2B). We found that negative feedback enhances the noise (i.e., peaks in Fano factors are higher) compared to the no-feedback case. Moreover, our expression of the threshold (Eq. 5) can roughly locate the positions of the peaks in Fano factors (Fig. 2B). In addition to our simulation results, we numerically solved the approximate second-order moment equations derived from the Master equation (Eq S2.1-S2.9 and S6.1-S6.20 in SI), though these solutions poorly aligned with the simulation data except for extreme cases of no-feedback and very high feedback (Fig. 2B, solid curves).

To understand how the calatyticity parameter (*α*) affect the threshold behavior, we plotted both the mean and Fano factors of mRNA copy number in the two-dimensional parameter-space of *k*_*r*_ − *α* (see the heatmaps in Fig 2C-H). Note that our derived expression of the threshold (Eq. 3 and 5) represents a curve in the *k*_*r*_ − *α* plane separating the expressed and repressed regimes in the heatmaps of mean mRNA (see solid curves in Fig. 2C-E). In the Fano-factor heat maps, the high noise regions are concentrated around the threshold curves for the limiting cases of no-feedback and very high feedback (Fig. 2F and 2H). For intermediate feedback strengths, the high noise regime broadly spans around the threshold curve (see the high-intensity zone in Fig. 2G). Nevertheless, the high noise regime remains bounded within the threshold curves corresponding to the limiting cases of no-feedback and very high feedback (dashed curves in Fig. 2G).

Since the miRNA transcription rate in the ON state (*k*_*s*_) is another important parameter, we next explored how the variation of *k*_*s*_ modulates the expression threshold and the mRNA noise around the threshold (Fig. 3). Similar to the derivation of a threshold mRNA transcription rate, we also derived an approximate expression of the threshold miRNA transcription rate 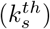, given by (for any nonzero feedback strength):

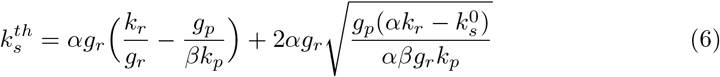

**Fig 3.**
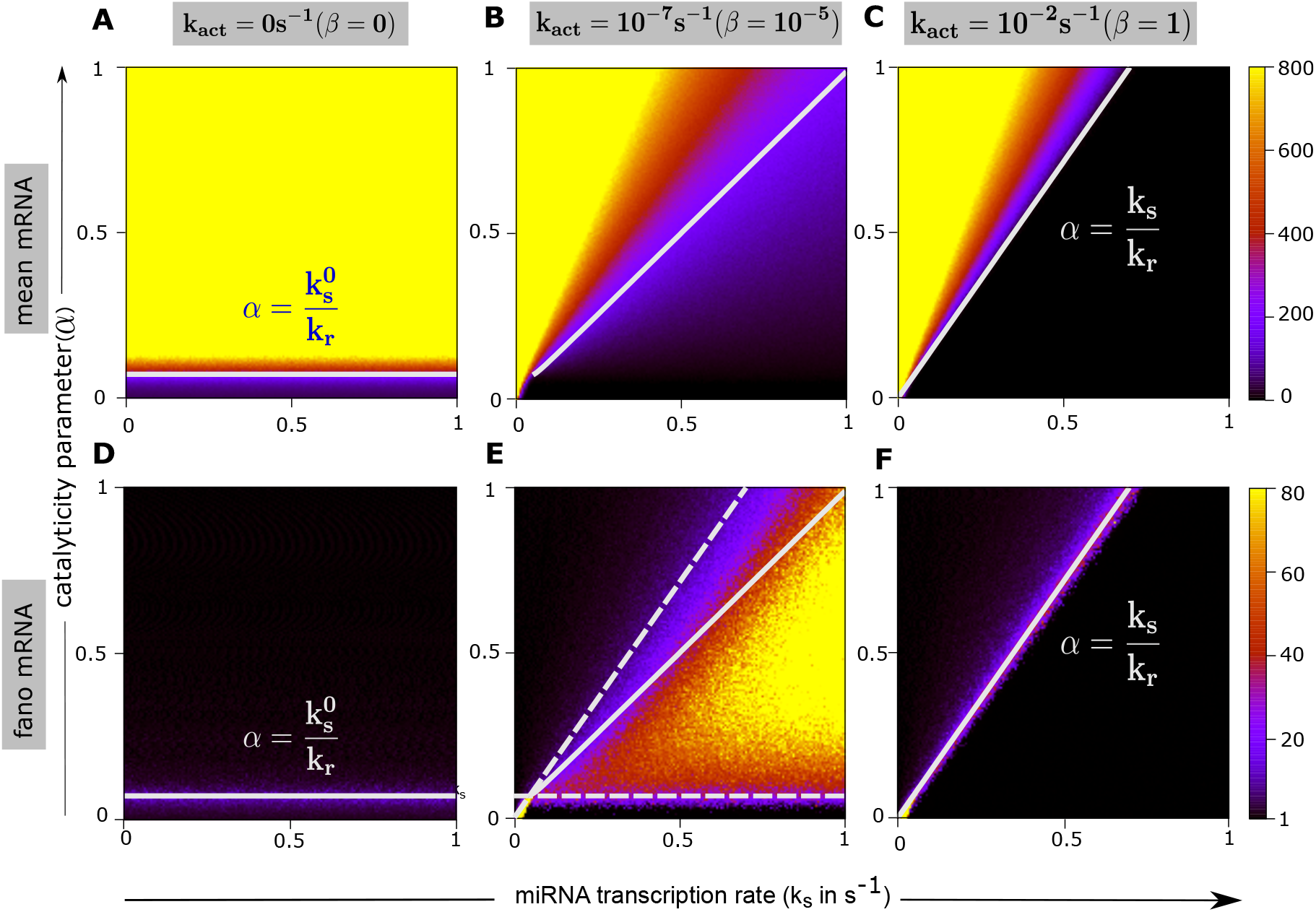
Effects of varying miRNA transcription rate on expression thresholds and Fano factors for different feedback strengths. Heat plots of steady-state means (A-C) and Fano factors (D-F) are shown in the *k*_*s*_ − *α* plane for different *k*_*act*_. Similar to Fig 2, we observed expression thresholds and high noise around these thresholds. The solid white line in each plot locates the threshold curves (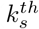 from Eq. 6). In panel E, the high noise regime lies between two limiting threshold boundaries corresponding to no-feedback and high feedback cases (white dash lines in E are re-plotted from panels D and F). Parameters were: *k*_*r*_ = 0.7*s*^−1^, *k*_+_ = 1*s*^−1^, and others were taken from Table 1.

For very high feedback (*β* → ∞), the expression of 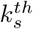 simplifies to 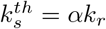 (However, *k*_*s*_ has no role for the no-feedback case as the miRNA production happens always at the basal rate,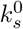). The expression of 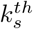 again correctly identified the separation between the expressed and repressed regimes in the *k*_*s*_ − *α* plane (see heat plots of the mean mRNA in Fig. 3A-C). Also, the high noise regimes spanned around the curves of 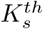 in heat plots of Fano factors (Fig. 3D-F). As before, for intermediate feedback strengths, the high noise regime is broader than the limiting cases, and it is bounded within the corresponding threshold curves of the limiting cases (see dashed curves in Fig. 3E).

We further investigated the role of the association between mRNAs and miRNAs in the transitions of expression levels around the thresholds (Fig S1, S2). We found that a higher effective association rate (g) mainly led to a more sharpened transition (see Fig. S1A). Moreover, lowering the effective association rate decreases the overall mRNA noise (see Fig S1F-H and S2D-F).

Together, we conclude that tuning the negative feedback strength modifies the threshold locations separating the high and low mean expressions. On the other hand, the effective association of miRNAs with the mRNAs largely determines the sharpness of the transition at the thresholds. Moreover, the mRNA noise peaks near the thresholds suggesting that the mRNA distributions could pass through qualitatively distinct regimes across the threshold points.

### 3.2 Bimodal mRNA distributions around the threshold

We next asked how the mRNA distribution changes across the expression threshold in the steady-state when the miRNA transcription rate is varied. We first focused on a regime where the miRNA-mRNA association is strong (i.e., *k*_+_ ≫ (*k*_−_ + *γ*), implying a high *g*) and the feedback strength is high (i.e., high *β* = *k*_*act*_*/k*_*deact*_). In this regime, the means of mRNA and miRNA showed sharp transitions around the threshold 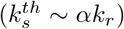, and they exhibited opposing trends with increasing miRNA transcription rate (*k*_*s*_) — see Fig. 4A. The mean mRNA (⟨*r*⟩) is higher than the mean miRNA (⟨*s*⟩) below 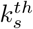, while ⟨*s*⟩ is higher than ⟨*r*⟩ above 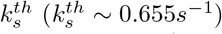. The miRNAs and mRNAs are equal to each other on average at the threshold (Fig. 4A), where both are present in small copy numbers suggesting that their coupled number fluctuations become vital near the threshold. Correspondingly, the correlation between mRNAs and miRNAs (quantified by the Pearson correlation coefficient, *ρ* = (⟨*rs*⟩ − ⟨*r*⟩⟨*s*⟩)*/σ*_*r*_*σ*_*s*_, where *σ*_*r*_ and *σ*_*s*_ are standard deviations) showed a sharp deep at the threshold point (Fig. 4B) signifying high anticorrelation between mRNAs and miRNAs. Thus, due to strongly coupled and enhanced number-fluctuation near the threshold, mRNAs randomly switch between ‘miRNA-bound’ and ‘miRNA-unbound’ states biasing either repression or expression respectively, which in turn could generate substantial expression variability. Accordingly, the steady-state mRNA distribution became bimodal near the threshold (Fig. 4C); however, the distribution was bell-shaped (approaching a Poisson-like shape) above the threshold, and it was like an exponential below the threshold (in the repressed regime). This shape transition of mRNA distributions could be further elucidated by simulating the time traces of the free miRNA and mRNA molecules across the threshold (see Fig. 4 D-F). As anticipated, both miRNAs and mRNAs showed high anticorrelated fluctuations near the threshold, toggling stochastically between almost zero and non-zero numbers (see Fig. 4E). But, only one species (either mRNA or miRNA) dominated over the other away from the threshold (Fig. 4D and 4F).

**Fig 4.**
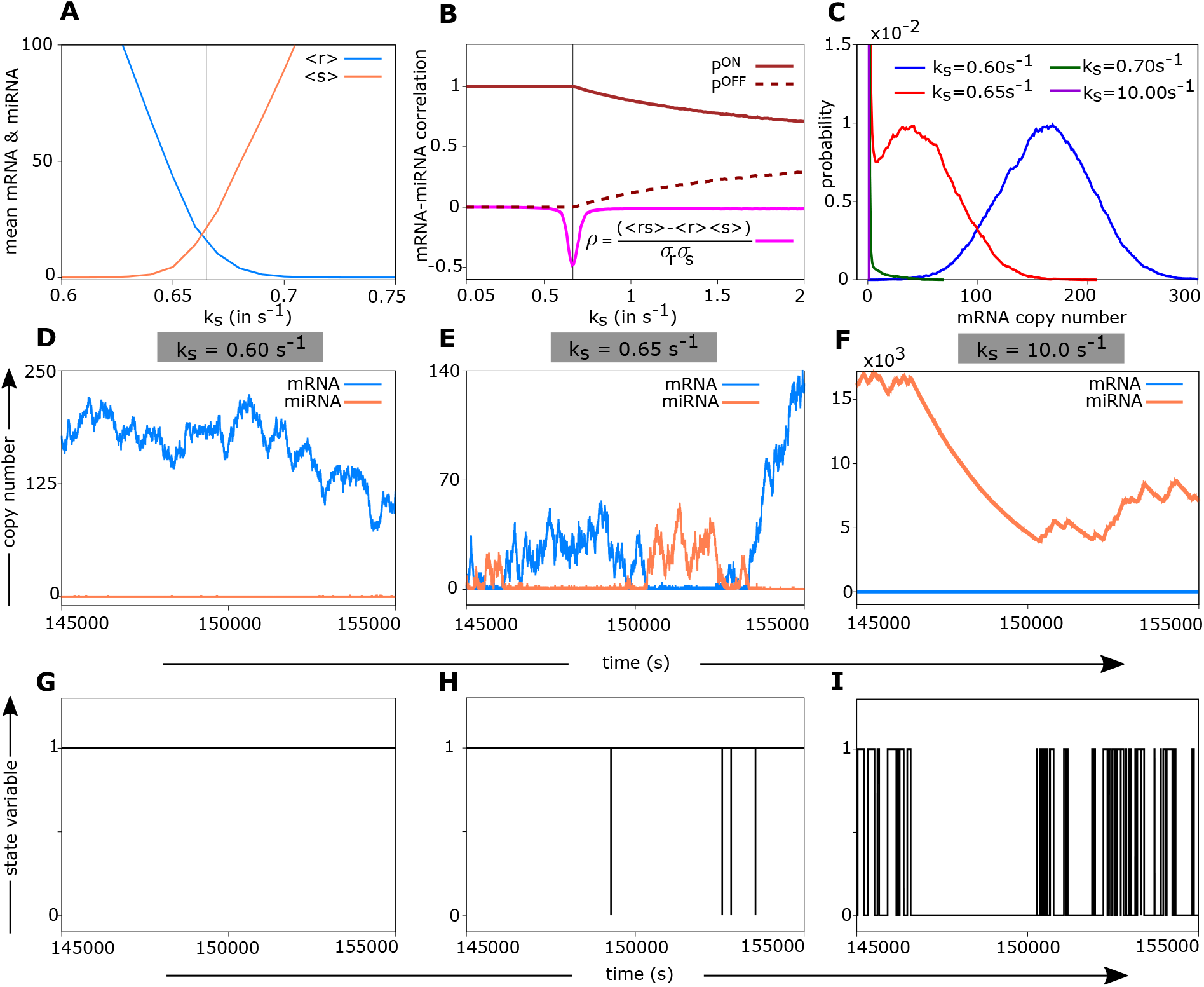
Enhanced and anticorrelated copy-number fluctuation of mRNAs and miRNAs link with bimodal mRNA distributions near the threshold. (A) Threshold behaviors of mean mRNA and mean miRNA are shown as functions of miRNA transcription rate (*k*_*s*_). (B) The mRNA-miRNA Pearson correlation (*ρ*) are plotted against *k*_*s*_. The probability of the miRNA-coding gene to be in the activated state (*P*^*ON*^) or basal state (*P*^*OFF*^) are also shown. The black vertical lines in panels A and B indicate the threshold 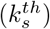, where mRNA-miRNA correlation showed a sharp deep in panel B. (C) Shape changes in steady-state mRNA distributions spanning the threshold region. Note the bimodal distribution near the threshold (red curve). (D-F) Temporal evolution of mRNA copy number (*r*(*t*)) and miRNA copy number (*s*(*t*)) at different *k*_*s*_ values across the threshold. Around the threshold, miRNAs and mRNAs are present in small numbers and show large anticorrelated fluctuations (E). Away from the threshold, either mRNA or miRNA dominates over the other (D and F). (G-I) Time traces of the state of the miRNA-coding gene at different *k*_*s*_ values spanning the threshold,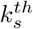. Here, ‘1’ and ‘0’ denote activated (ON) and basal (OFF) states of the miRNA-coding gene, respectively. In panels D-I, the same time windows were chosen to make correspondence. We used high feedback strength and high miRNA-mRNA association strength (*k*_*act*_*/k*_*deact*_ = 1 and *k*_+_*/k*_−_ ≈ 278), while other parameters were from Table 1.

Since the miRNA-coding gene switches between ON (activated) and OFF (basal) states due to negative feedback, we also checked how the miRNA gene states behave across the threshold where bimodal expression was observed. Note that free mRNAs dominate over the miRNAs below the threshold (Fig 4A), and hence the proteins translated from free mRNAs could push the miRNA gene mostly into the ON state. This expectation was consistent with the observed behavior of state probabilities (*P*_*on*_ and *P*_*off*_) across the threshold (Fig. 4B). We further monitored the time traces of miRNA gene states (denoted by 1 for ON and 0 for OFF), shown in Fig. 4G-I. Below the threshold, the miRNA gene was in ON state (Fig. 4G), and it mostly was in OFF state above the threshold (Fig. 4I).

Intuitively, the miRNA gene is expected to be in the OFF (basal) state when the feedback is negligible (*β* ∼ 0), while it could be mostly in the ON state for very high feedback (*β* → ∞) (see Eq. 4). Between these two extreme limits, the state switching of the miRNA gene could be too frequent, leading to expression variability. Therefore, we anticipate that two different processes could collectively modulate the emergence of bimodality: (i) the switching between miRNA-bound and miRNA-unbound states of free mRNAs near the threshold, i.e., the complex formation and dissociation dynamics (governed by the effective association rate, *g*), and (ii) the switching between ON (activated) and OFF (basal) states of the miRNA gene, i.e., the protein-mediated activation and deactivation of the miRNA gene (governed by the feedback strength, *β*).

### 3.3 Tuning miRNA-mRNA association and feedback strength modulate the emergence of bimodality

We asked how the effective association rate (*g* = *k*_+_*γ/*(*k*_−_ + *γ*)) and the feedback strength (*β* = *k*_*act*_*/k*_*deact*_) affect the emergence of bimodal mRNA distributions in the steady-state. To this aim, we systematically varied the mRNA-miRNA binding/unbinding rate (*k*_+_ or *k*_−_) and the miRNA gene activation rate (*k*_*act*_) to monitor how the mRNA distributions change across the expression threshold. Specifically, for a wide range of *g* (0.0001*s*^−1^ − 0.1*s*^−1^) and *β* (0 − 1), we first identified the threshold mRNA transcription rate 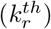, and then investigated the mRNA distributions across the threshold (see Fig. 5). For a strong miRNA-mRNA association (*g* ∼ 0.01*s*^−1^ − 0.1*s*^−1^), bimodal mRNA distributions appeared near the threshold irrespective of the absence and presence of feedback (i.e., for both *β* = 0 and *β* ≠ 0) (see Fig 5A-F). Thus, for a strong miRNA-mRNA association, the switching between miRNA-bound and unbound states of the mRNAs primarily leads to bimodal distributions since bimodality was already present in the no-feedback case (Fig. 5A and 5D), and bimodality persisted when feedback was introduced. On the other hand, for a minimal miRNA-mRNA association (*g* = 0.0001*s*^−1^), bimodality disappeared both in the presence and absence of feedback (see Fig. 5M-O) as miRNAs hardly bound with mRNAs in this regime. More interestingly, for intermediate-level of miRNA-mRNA association (*g* ∼ 0.0004*s*^−1^ − .001*s*^−1^), bimodal distributions emerged only when the feedback strength was intermediate (see Fig. 5H and 5K); while bimodality disappeared for the extreme cases of no-feedback (Fig. 5G and 5J) and very high feedback (Fig. 5I and 5L). These observations suggest that the switching between ON (activated) and OFF (basal) states of the miRNA gene could be the determining factor for the bimodality when the miRNA-mRNA association (*g*) was intermediate. Note that the miRNA gene was always in the OFF (basal) state in the no-feedback case. However, for an intermediate feedback strength, the miRNA gene’s average residency time in the OFF and ON states kept changing across the threshold (see Fig. 6A-F, corresponding to Fig. 5H). Below the threshold 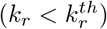, the mean mRNA (and correspondingly translated transcription factors) was low, and hence the miRNA gene mainly was in the OFF state (Fig. 6A and 6D). Conversely, the miRNA gene was largely in the ON state above the threshold as the mean mRNA and translated transcription factors were high (Fig. 6C and 6F). Near the threshold, the miRNA gene spent almost a similar amount of time in ON and OFF states, leading to the mRNA copy number variability and subsequent bimodality. In contrast, such frequent toggling between ON and OFF states could not occur when the feedback strength was very high — here, the miRNA gene mostly stayed in the ON state across the threshold (see Fig. 6G-L, corresponding to Fig. 5I), reducing the heterogeneity in expression. Therefore, the miRNA-gene state switching (due to negative feedback) plays a key role in modulating bimodality for an intermediate miRNA-mRNA association rate.

**Fig 5.**
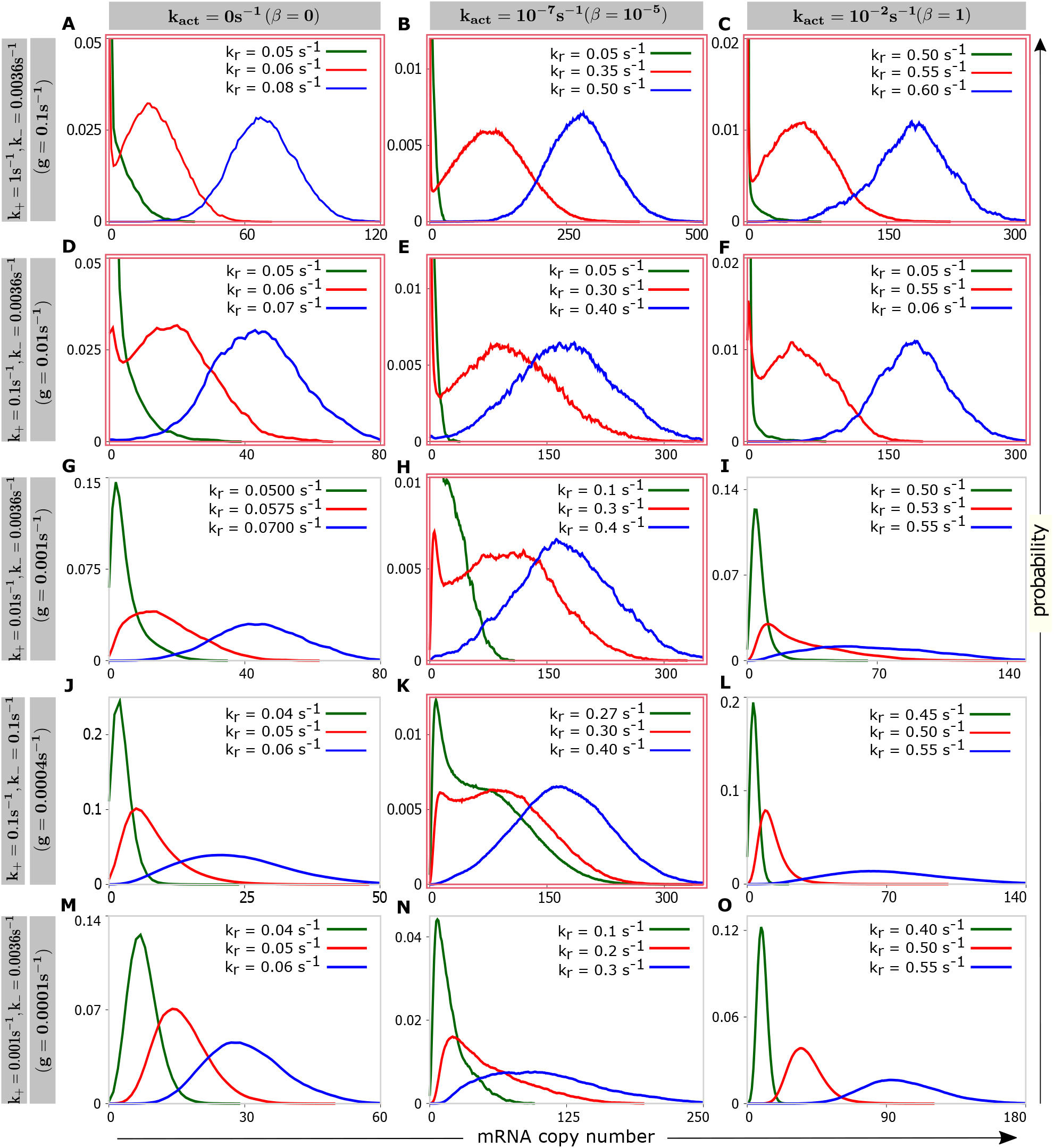
Effects of miRNA-mRNA binding affinity and feedback strength on the emergnece of bimodality. Steady-state mRNA distributions at different mRNA transcription rates (*k*_*r*_) are shown by varying the miRNA-mRNA association/dissociation rate (*k*_+_ or *k*_−_) and the activation rate of the miRNA-coding gene (*k*_*act*_). For strong miRNA-mRNA association, bimodal mRNA distributions appeared both in the absence and presence of feedback (A-F). Conversely, bimodality disappeared for a weak miRNA-mRNA association (M-O). For intermediate association strength, bimodal distributions did not emerge for the limiting cases of no-feedback (G, J) and very high feedback (I, L); bimodality emerged only when the feedback strength was intermediate (H, K). We used *k*_−_ = 0.1*s*^−1^ for panels (J-L) and for rest of the panels *k*_−_ = 0.0036*s*^−1^. *k*_+_ and *k*_*act*_ are varied as shown in figure. *α* = 0.95 and other parameters were from Table 1. Panels containing bimodal distributions are marked with red double-lined border.

**Fig 6.**
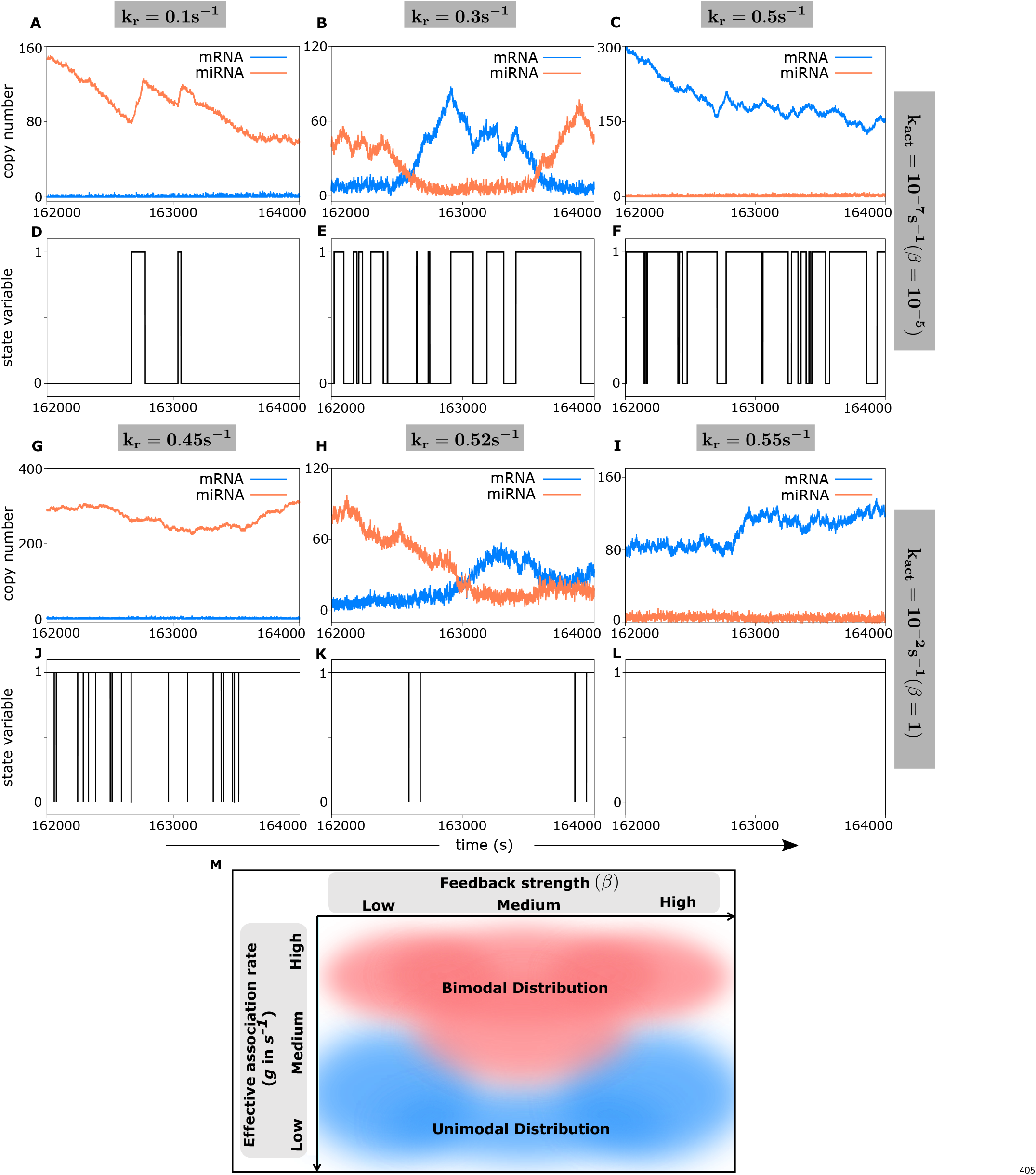
State switching of the miRNA-gene and stochastic time-traces can explain the ‘unimodal-bimodal phase diagram’. Switching of the miRNA gene and time-traces of miRNAs and mRNAs are shown for an intermediate level of miRNA-mRNA effective association (*g* = 0.001*s*^−1^). Panels A-F are for *β* = 10^−5^ corresponding to Fig. 5H; while panels G-L corresponds to Fig. 5I for a very high feedback strength (*β* = 1). For *β* = 10^−5^, we observed anti-corelated fluctuations in mRNA and miRNA copy numbers near the threshold (B) and frequent toggling of the miRNA gene between ‘ON’ and ‘OFF’ states (E), which corresponds to the bimodal mRNA distribution (in Fig 5H). Increasing the feedback strength creates an overall bias towards the ‘ON’ state (J-L). For a high feedback strength, the frequency of state toggling near the threshold is markedly reduced (compare panels E and K). The parameters were *k*_*s*_ = 0.5*s*^−1^, *k*_+_ = 0.01*s*^−1^, *α* = 0.95. Other parameters were as in Table 1. (M) A schematic phase diagram showing the regions of unimodal and bimodal mRNA distributions in the *β* − *g* plane.

We thus conclude that (i) negative feedback strength (*β*, controlling the miRNA gene state switching) and (ii) the effective association rate (*g*, controlling the mRNA-miRNA binding strength) are the main governing factors for bimodal distributions. We can represent the emergence of bimodality as a schematic 2D phase-diagram in the *β* − *g* parameter space (see Fig. 6M). Evidently, miRNA-mediated negative feedback broadens the parameter regime of bimodal mRNA distributions since bimodality could be observed for intermediate miRNA-mRNA association strength as well, not only for a strong miRNA-mRNA association (Fig. 5H, 5K, and 6M).

A question naturally arises, how does the miRNA-mediated repression compare with a simple auto-repression (i.e., a gene producing its own repressors) since both motifs are ubiquitous in various biological contexts?

### 3.4 A comparison of miRNA-mediated negative feedback with an auto-repression

There already exist several theoretical [36, 80, 81] and experimental studies [19, 61] of auto-regulatory negative feedback where a gene expresses its repressor. A general understanding is that an auto-regulatory negative-feedback motif can buffer the gene expression noise [19, 61, 80, 81], but an auto-regulatory positive-feedback motif generally exhibits bistability in expression [23, 26, 35, 42, 64].

Here we revisited a model of an auto-repression motif to compare it with our model of miRNA-mediated negative feedback (see Fig. 7A and 7B). Note that the motif of miRNA-mediated negative feedback represents a scenario of indirect repression, i.e., repression through intermediate steps of miRNA-gene activation (Fig. 7A). In contrast, the simple negative feedback represents direct repression or auto-repression, where the gene produces a protein (repressor) that binds with the same gene and reduces its transcription rate from the basal rate (Fig. 7B). In this auto-repression motif, the binding and unbinding rates of the repressor are *k*_*off*_ and *k*_*on*_, respectively. The repressor-bound gene transcribes mRNAs at a lower rate 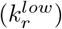, and the transcription rate for the repressor-unbound gene is higher 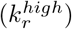.

**Fig 7.**
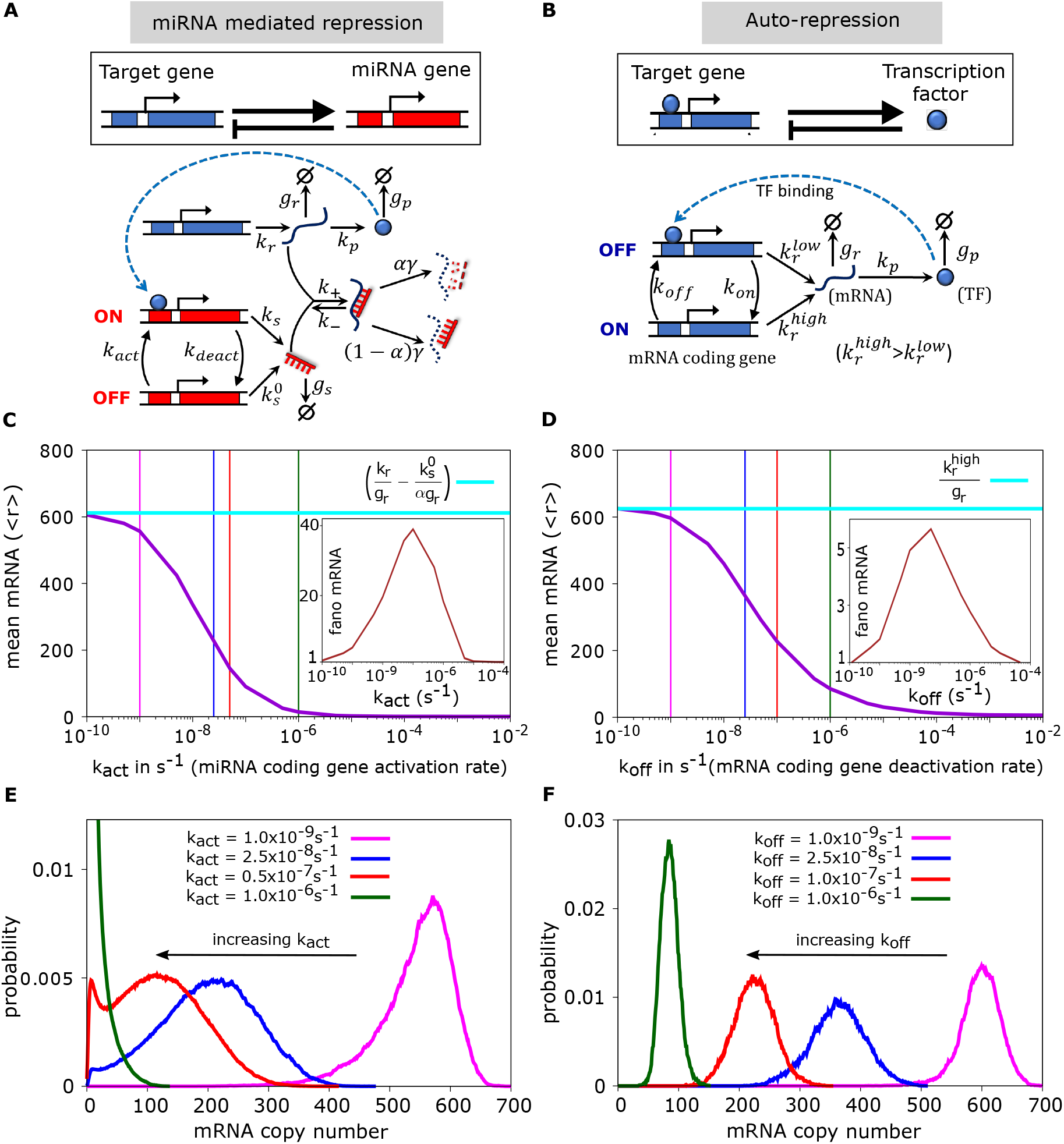
A Comparison of miRNA-mediated negative feedback with auto-repression shows distinctive features in target mRNA distributions. (A) Our model of a miRNA-mediated negative feedback loop (same as in Fig 1). (B) Schematic and detailed model of a negative auto-regulatory motif where a gene produces its own repressor. In A, the miRNA production increases as *k*_*act*_ increases, which in turn suppresses target mRNAs. In B, for higher *k*_*off*_, the mRNA production rate switches from 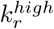 to 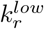. Thus, both *k*_*act*_ and *k*_*off*_ produce a bias towards repression in respective motifs, when *k*_*deact*_ and *k*_*on*_ are fixed. (C, D) Means and Fano factors of mRNA are shown by varying *k*_*act*_ (for miRNA-mediated negative feedback) and *k*_*off*_ (for auto-repression). In both cases, means cross over from high to low expression regimes as the bias towards repression increases, and the Fano factors peak near the cross-over region. The vertical lines in C and D represent the locations where mRNA distributions were sampled and shown in panels E and F. (E, F) Steady-state mRNA distributions for respective motifs. Parameter values of *k*_*p*_, *g*_*p*_, and *g*_*r*_ were the same in both motifs (as in Table 1), while other parameters were chosen in such a way that the mean curves look similar in both cases. Parameters for C and E were *k*_*r*_ = 0.25*s*^−1^, *k*_*s*_ = 0.5*s*^−1^, 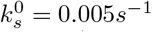, *k*_+_ = *k*_−_ = 0.1*s*^−1^, *α* = 0.95, while others from Table 1. For D and F, we used 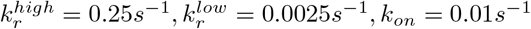, *k*_*on*_ = 0.01*s*^−1^.

In order to compare the outputs of the two motifs on an equal footing, we first set the same protein synthesis rate and the same degradation rates of mRNAs and proteins in both motifs (*k*_*p*_, *g*_*p*_, and *g*_*r*_ in Fig. 7A-B). We then chose other parameters in such a way that in both motifs, the mean mRNA levels are the same in the limit of no-feedback (when *k*_*act*_*/k*_*deact*_ → 0 or *k*_*off*_ */k*_*on*_ → 0 in respective motifs, see Fig. 7A-B). These limiting means are roughly 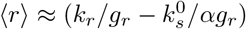 for the miRNA-mediated repression (Eq. 3), and 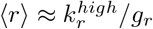 for the auto-repression. We further noted that increasing *k*_*off*_ (i.e., the deactivation rate of the target gene by repressors) biases the auto-repression motif towards a low expression regime (Fig. 7B); while increasing *k*_*act*_ (i.e., the activation rate of the miRNA gene by proteins) push the miRNA-mediated repression motif towards low expression. Thus, increasing *k*_*off*_ or *k*_*act*_ in respective motifs can increase the negative feedback strengths (when other parameters are fixed). We, therefore, varied *k*_*off*_ or *k*_*act*_ in respective motifs and calculated the mean mRNA and its Fano factors, as shown in Fig. 7C-D. As expected, both motifs exhibited similar transitions of mean mRNAs from high to low values when the bias towards repression was increased by varying respective feedback strengths (Fig. 7C-D). The Fano factors also peaked near the cross-over region (see insets, Fig. 7C-D). Thus, both motifs are qualitatively indistinguishable at the mRNA mean and noise level. However, when we monitored the steady-state mRNA distributions across the peak regions of Fano factors, the bimodal distribution was observed in the miRNA-mediated repression (Fig. 7E), but the auto-repression did not produce bimodality (Fig. 7F). In fact, it was known from a recent experiment [19] that direct negative feedback or auto-repression does not produce bimodal distributions. On the contrary, as discussed in the previous section, the bimodality in miRNA-mediated repression stems from the frequent switching of miRNA-gene between ON (activated) and OFF (basal) states near the high fano-factor region; while one of the states dominated away from this region (Fig. S3). Together, we conclude that the indirect negative feedback via miRNAs can potentially generate phenotypic diversity compared to direct negative feedback.

## 4 Discussion

### Summary

Negative feedback loops involving miRNAs are often found in several gene regulatory networks governing diverse cellular processes like cell differentiation and cancer progression [1, 12, 15, 21, 44, 48, 51, 52, 54, 78, 92]. In order to investigate the effects of a miRNA-mediated single-negative-feedback loop (SNFL), we here built a stochastic framework describing the competitive titration between miRNAs and their target mRNAs. Earlier theoretical studies have suggested that there are essentially three factors governing gene expression in miRNA-dependent SNFL [8, 89, 93]: (i) activation and deactivation of the miRNA-coding gene by the TF, (ii) association-dissociation between miRNAs and target mRNAs, and (iii) miRNA’s catalytic mode of action on target mRNAs. However, previous models did not incorporate the above three factors altogether. In contrast, our detailed model incorporates all these factors, each of which significantly regulates miRNA-mediated gene expression (Fig. 1). In our model, a gene produces a specific protein that acts as a transcriptional activator of a miRNA-coding gene and, in turn, the miRNA suppresses the transcriptional factor (TF) itself. The output of the motif largely depends on three key parameters (see the Model section): the negative feedback strength (denoted by *β* = *k*_*act*_*/k*_*deact*_), the effective miRNA-mRNA association rate (*g* = *k*_+_*γ/*(*k*_−_ + *γ*)), and the catalyticity parameter (*α*). A systematic model analysis by varying these parameters revealed some interesting aspects of miRNA-mediated negative feedback.

With increasing mRNA transcription rate (or decreasing miRNA transcription rate), the mean mRNA at the steady state showed a threshold-like behavior transiting from a low to a high level (Fig. 2A, 2C-E, 3A-C). Such behavior was previously observed in experiments, and theoretical studies that did not incorporate feedback [9, 58, 62]. The presence of negative feedback further modifies the threshold points, which we found analytically using mean-field approximation (Eq. 5, 6). We also quantified the gene expression noise at the steady state by the mRNA Fano factor (*var*(*r*)*/* ⟨*r*⟩). This intrinsic noise displayed a peak in the vicinity of the threshold (Fig 2B) similar to the previous study in the absence of feedback [62]. However, feedback enhances the noise near the respective thresholds (Fig. 2B).

Away from the threshold, one species (either miRNA or mRNA) dominates over another, but in the proximity of the threshold, their copy numbers are low and near-equal (Fig. 4A), representing a titration-like binding process to form miRNA-mRNA complexes. The miRNA-mRNA titration introduces a negative correlation between these two species that is enhanced at the expression threshold (Fig. 4B). Either all of the miRNAs or mRNAs are fully bound in miRNA-mRNA complexes away from the threshold. Near the threshold, however, mRNAs stochastically become free or miRNA-bound, leading to large anticorrelated fluctuations of miRNA and mRNA numbers (Fig. 4D-F). This stochastic fluctuation between expressed and repressed states (Fig. 4E) is manifested as a noise-induced bimodality in mRNA distributions (Fig. 4C) in the proximity of the threshold. This emergence of bimodality can be further tuned by the strength of negative feedback and the miRNA-mRNA association rate (Fig. 5 and 6M).

Next, we quantitatively compared our model with an auto-repression (Fig. 7A-B). In miRNA-mediated negative feedback, the suppression of transcription factors occurs indirectly via miRNAs, whereas in auto-repression, TFs directly suppress themselves, acting as repressors of their gene. In both circuits, as expected, the mean mRNA displayed a transition from high to low expression with increasing negative feedback strength, and the mRNA Fano factor increased around the transition region (Fig 7C-D). Comparing the peak heights of the mRNA Fano factor, we could conclude that miRNA-mediated feedback is noisier than auto-repression — this is in line with previous results that auto-repression suppresses heterogeneity of gene expression [19, 61, 80, 81]. Though the target gene expression exhibited similar characteristics at the mean and noise level, the contrast between these two types of negative feedback circuits lies at the distribution level (Fig. 7E-F). The miRNA-mediated negative feedback produced bimodality in mRNA distributions, whereas auto-repression did not, suggesting the crucial roles of miRNAs in generating phenotypic diversity.

### Biophysical relevance

A crucial understanding revealed by our study is that two processes largely control the bimodality of mRNA distributions in a miRNA-based negative feedback loop: (i) switching of target mRNAs between miRNA-bound and miRNA-free states (quantified theoretically by the effective miRNA-target association rate, *g*), and (ii) the TF-dependent switching of miRNA-coding gene between basal and activated states (quantified by the negative feedback strength *β*). Thus, two parameters (*g* and *β*) collectively modulate the emergence of bimodal mRNA distributions (Fig. 5), and we can visualize this in a 2D phase diagram (Fig. 6M), linking model parameters with a qualitative prediction. We further investigated if the bimodality observed at the mRNA level would also be translated at the protein level. We found that the steady-state protein distributions are also bimodal when the protein degradation rate is higher than the protein synthesis rate (Fig. S4). In contrast to our findings, another theoretical study on miRNA-based negative feedback reported bell-shaped or long-tailed unimodal distributions of proteins [89]. This difference may arise due to the coarse-grained modeling in the latter study as compared to our detailed modeling. On the other hand, deterministic modeling of miRNA-based negative feedback [93] missed the rich interplay between negative feedback and miRNA-target titration in modulating gene expression noise, as elucidated here.

### Future directions

Though our analytical mean-field approach revealed the crucial dependence of the expression thresholds on model parameters (Eq. 5, 6), exact expressions of the Fano factors and the time-dependent distributions are challenging to obtain. Solving the highly nonlinear chemical Master equation (Eq. 1) for our model would be an open problem. A recent theoretical paper advocated the Linear Mapping Approximation (LMA), a helpful method by which bimolecular interactions can be substituted by zeroth or first-order reactions, reducing a nonlinear Master equation into a linear one and making the solution easier [13]. Applying LMA in our model would be an interesting theoretical direction to pursue.

Our model focused on direct miRNA-mRNA interaction and neglected some other aspects of miRNA biogenesis. For instance, precursor miRNAs ultimately transform into mature miRNAs via several intermediate steps, and the precursor miRNAs also compete with mature miRNAs over the same target mRNA population [68]. The miRNA maturation could lead to time delays that, combined with negative feedback, may generate interesting features like stochastic oscillations in expression [4, 11]. Moreover, during the miRNA maturation process, a single miRNA locus can generate a series of sequences, typically known as isomiRs, [33, 60], which can bind with the same target mRNAs. Such competitive dynamics of precursor miRNAs or isomiRs with matured mRNAs are still unexplored. Note that such competition between several kinds of non-coding RNA sequences sharing a common target mRNA pool is conceptually different from the ceRNA network, where several target mRNAs share the same miRNAs.

Several experimental techniques, including synthetic design of genetic circuits, fluorescent microscopy, and flow cytometry, have been used in single-cell experiments and bulk expression measurements related to miRNA-dependent gene expression [10, 58, 79]. Such experiments can provide accurate quantitative testing of theoretical predictions as described in this paper. Our results further suggests that cells may chemically control many processes involving miRNAs, such as the catalytic mode of repression, miRNA-mRNA association rate, and the feedback strength to tune the gene expression noise. For instance, experimental studies have shown that the phosphorylation of RISC proteins can control the mature miRNA binding to target mRNAs [69], creating a regulating knob for the effective miRNA-mRNA association rate. Our study revealed the key tuning parameters that may be connected with experiments to know how expression diversity is regulated in a miRNA-based negative feedback loop.

## Materials and methods

We simulated the model using Gillespie algorithm [27]. The codes were written in FORTRAN90, and they are freely available in the following link: https://github.com/PhyBi/miRNA-negative-feedback.

## Supporting information

The Supporting Information contains four Supplemental Figures (Figs. S1-S4) and details of mathematical analysis of our model.

## Acknowledgments

DD thanks DST-SERB, Government of India (project No. SRG/2019/001068) for the financial support. RA thanks Council of Scientific & Industrial Research, India (CSIR) for his fellowship (File No: 09/921(0282)/2019-EMR-I).

## Notes

https://github.com/PhyBi/miRNA-negative-feedback

### Competing Interest Statement

The authors have declared no competing interest.

