## Supplementary Information for "Effects of microRNA-mediated negative feedback on gene expression noise"

November 9, 2022

**1** Department of Biological Sciences, Indian Institute of Science Education And Research Kolkata, Mohanpur, Nadia 741246, West Bengal, India.

**2** Centre for BioSystems Science and Engineering, Indian Institute of Science,Bengaluru 560012, India.

†
‡

#### Supplemental Figures

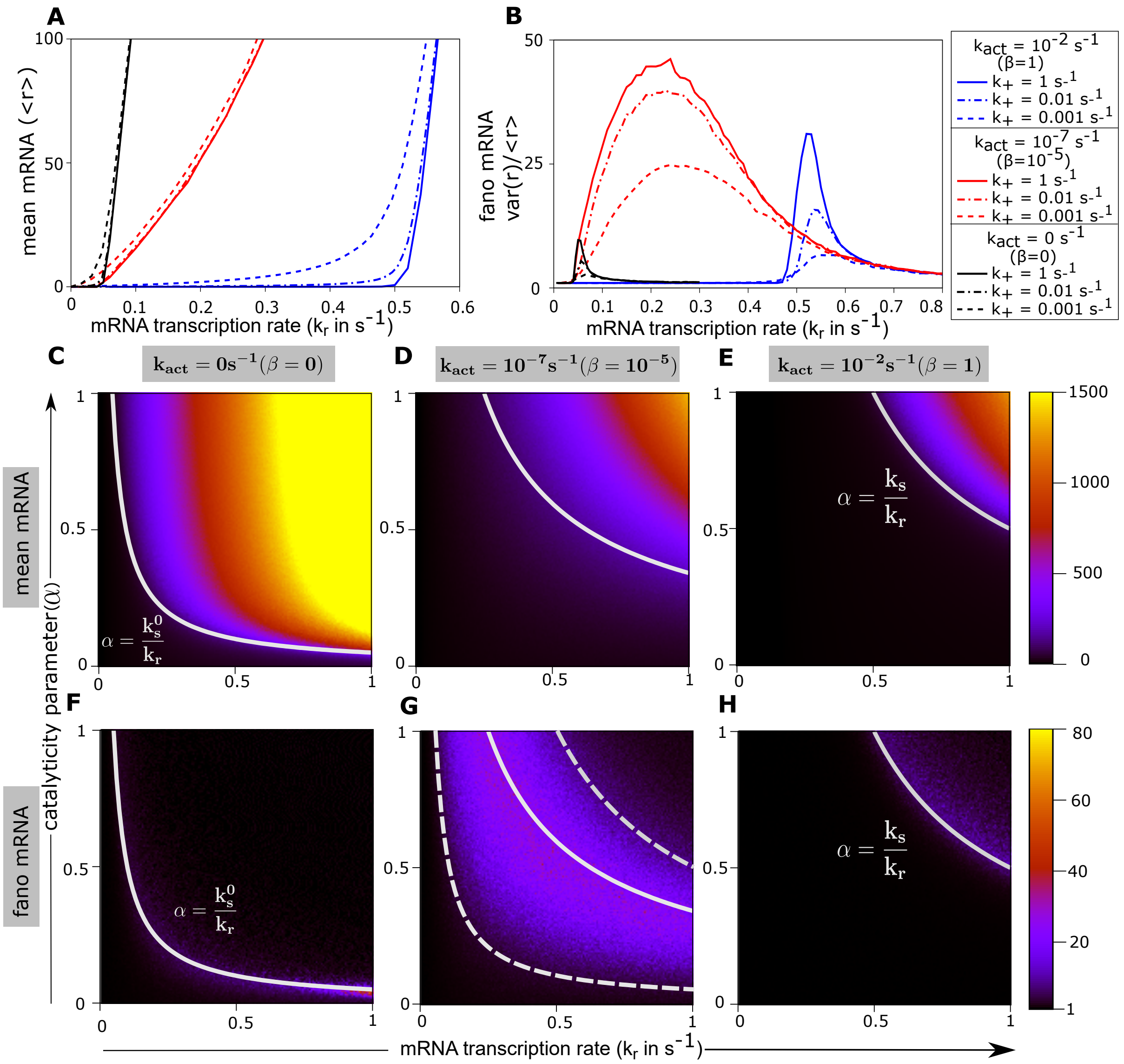


Figure S1: **(Related to Fig. 2) Stronger miRNA-mRNA association produces sharper transition in mean mRNA around thresholds and increases intrinsic noise.** (A-B) Mean and Fano Factors of target mRNA are shown for different combinations of mRNA-miRNA association rates ($k_+$) and miRNA gene activation rate ($k_{act}$). Lowering the effective association rate ($g = (k_+\gamma)/(k_- + \gamma)$) makes the transition shallow from repressed to expressed regime (A), and reduces the mRNA noise near threshold (B). (C-G) Heatplots of mRNA mean and Fano Factor in $k_r - \alpha$ plane for comparable mRNA-miRNA association and dissociation rates (i.e., $k_+ \approx k_-$). The solid and dashed lines in each heatplot denote thresholds ($k_r^{th}$) for respective cases as in Fig 2 (main text). Note that the overall mRNA noise is reduced due to the decrease in effective association rate (compare panels F-H with Fig. 2F-2H.). For all panels, $k_s = 0.5s^{-1}$ and $\alpha = 0.95$ for panels A-B. For the heatplots (C-H), $k_+ = 0.001s^{-1}, k_- = 0.0036s^{-1}$. Other parameters were taken from Table 1.

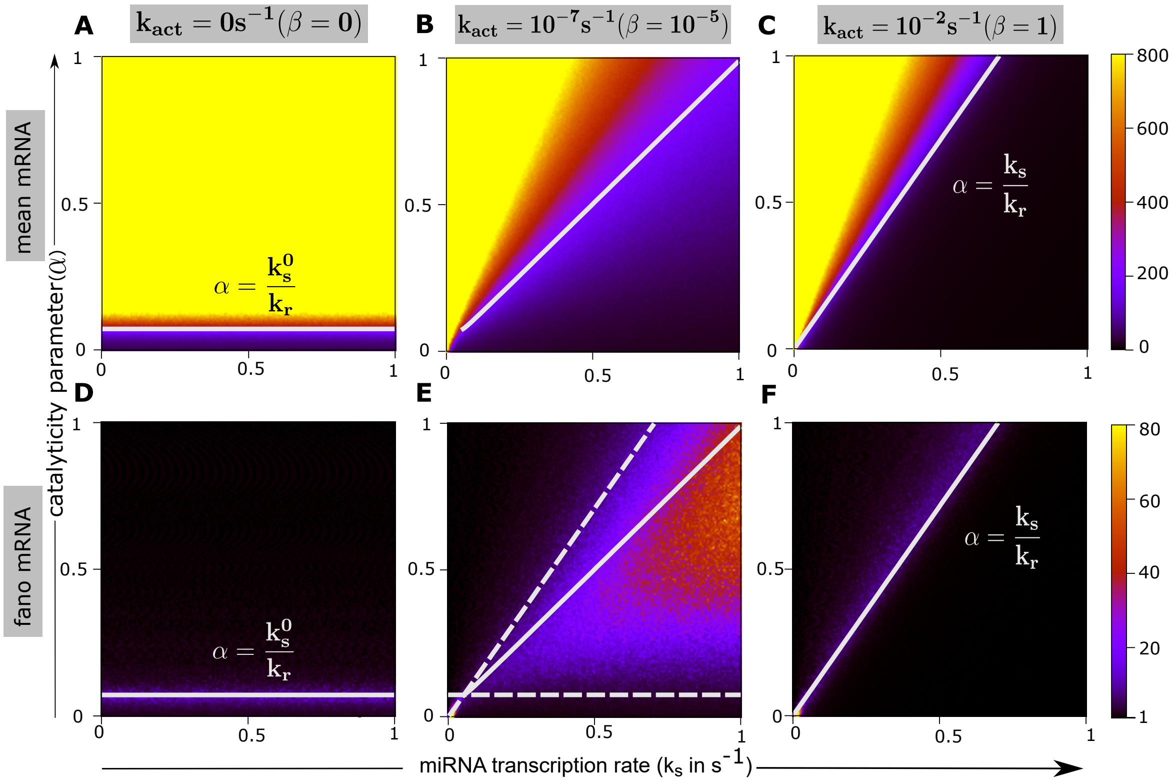


Figure S2: **(Related to Fig. 3) Effect of decreasing the miRNA-mRNA association rate on mRNA mean and noise.** Heatplots represent the steady-state mRNA mean (A-C) and Fano Factors (D-F) in the $k_s - \alpha$ plane for comparable mRNA-miRNA association and dissociation rates ($k_+ \approx k_-$). The solid lines in each heat plot locates the position of threshold curves ($k_s^{th}$) for the respective cases as in Fig. 3 (the dashed lines in panel E are thresholds corresponding to no-feedback and high-feedback cases, re-plotted from panels D and F). Though mRNA expression remains similar to Fig 3, the overall noise is reduced due to the decrease in effective association rate ($g$). We used $k_r = 0.7s^{-1}, k_+ = 0.001s^{-1}, k_- = 0.0036s^{-1}$. Rest of the parameter values were taken from Table 1.

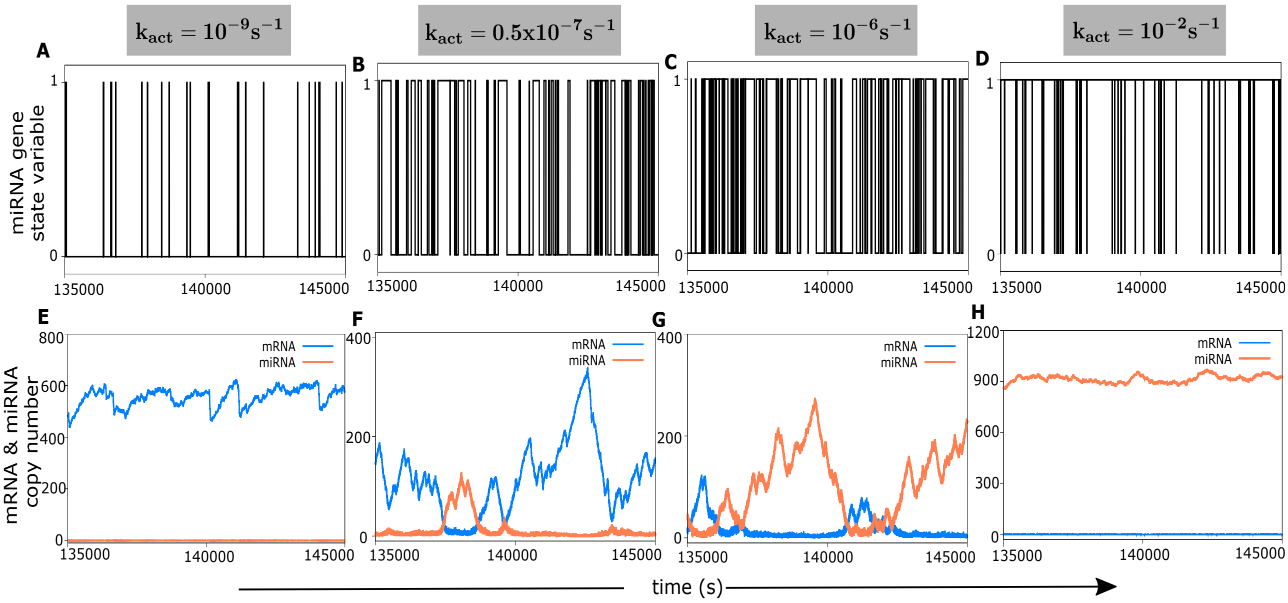


Figure S3: **(Related to Fig. 7) Effect of varying miRNA gene activation rate (i.e., the feedback strength) on the switching behavior of miRNA gene and copy number fluctuations.** For a wide range of activation rate ($k_{act}$), steady state time profiles of miRNA gene state (A-D), and corresponding time evolution of mRNA and miRNA counts (E-H) are shown. The panels correspond to the Fig. 7E in the main text. Note that for a very low or very high activation rate, the states are either biased towards OFF or ON states (A and D) and only one species dominates over the other (E and H). The parameters were $k_r = 0.25s^{-1}, k_s^0 = 0.005s^{-1}, k_s = 0.5s^{-1}, k_+ = k_- = 0.1s^{-1}, \alpha = 0.95s^{-1}$ and rest of the parameters were taken from Table 1.

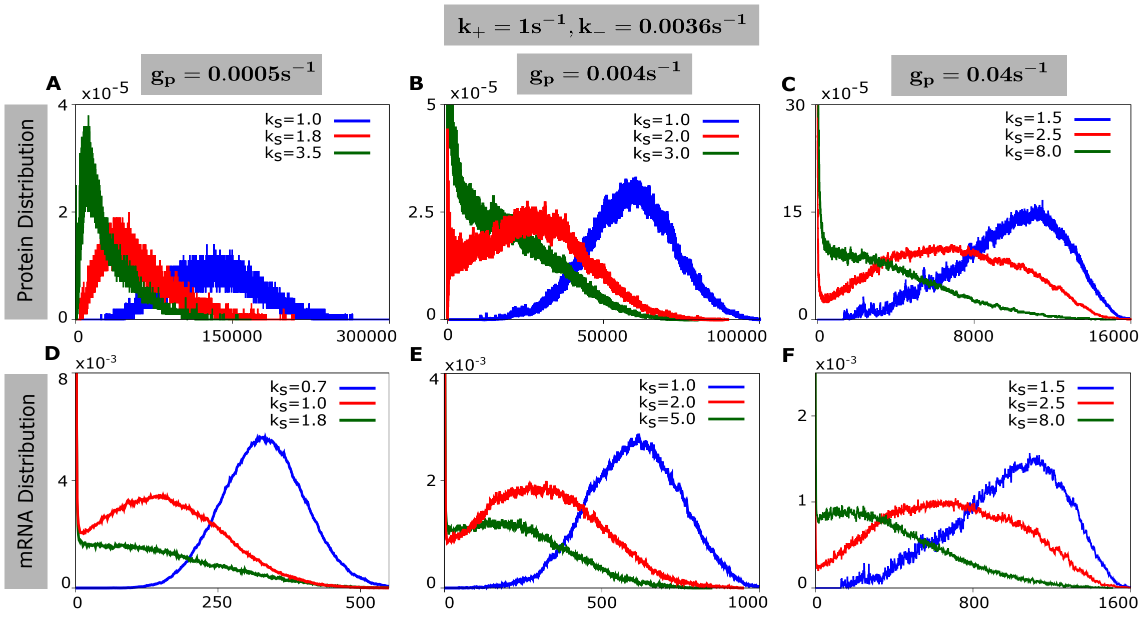


Figure S4: **Effect of varying the protein degradation rate ($g_p$) on the emergence of bimodal protein distributions.** Steady-state protein distributions are shown by varying the miRNA transcription rate ($k_s$) across the thresholds. Note that bimodality was seen when the protein degradation rate is higher compared to the protein synthesis rate ($k_p = 0.4s^{-1}$ for all panels). Corresponding mRNA distributions, however, were bimodal (not shown). The parameters were $k_r = 0.7s^{-1}, k_{act} = 10^{-7}s^{-1}, \alpha = 0.95$. Rest of the parameters were taken from Table 1.

### Reaction scheme of miRNA mediated negative feedback (Fig 1)

The complex scheme of this model can be thought of as a series of simple chemical reactions as follows.

$$\mathrm{G_{mRNA}} \xrightarrow{\mathrm{k}_r} \mathrm{mRNA} \tag{S 0.1}$$

$$\mathrm{mRNA} \xrightarrow{\mathrm{g}_r} \phi \tag{S 0.2}$$

$$\mathrm{mRNA} \xrightarrow{\mathrm{k}_p} \mathrm{Protein} \tag{S 0.3}$$

$$\mathrm{Protein} \xrightarrow{\mathrm{g}_r} \phi \tag{S 0.4}$$

$$\mathrm{Protein} + \mathrm{G_{miRNA}}^{\mathrm{OFF}} \underset{\mathrm{k_{deact}}}{\overset{\mathrm{k_{act}}}{\rightleftharpoons}} \mathrm{G_{miRNA}}^{\mathrm{ON}} \tag{S 0.5}$$

$$\mathrm{G_{miRNA}}^{\mathrm{OFF}} \xrightarrow{\mathrm{k}_s{}^0} \mathrm{miRNA} \tag{S 0.6}$$

$$\mathrm{G_{miRNA}}^{\mathrm{ON}} \xrightarrow{\mathrm{k}_s} \mathrm{miRNA} \tag{S 0.7}$$

$$\mathrm{miRNA} \xrightarrow{\mathrm{g}_s} \phi \tag{S 0.8}$$

$$\mathrm{mRNA{-}miRNA\ Complex} \underset{\mathrm{k}_+}{\overset{\mathrm{k}_-}{\rightleftharpoons}} \mathrm{mRNA} + \mathrm{miRNA} \tag{S 0.9}$$

$$\mathrm{mRNA{-}miRNA\ Complex} \xrightarrow{\alpha\gamma} \phi \tag{S 0.10}$$

$$\mathrm{mRNA{-}miRNA\ Complex} \xrightarrow{\alpha(1-\gamma)} \mathrm{miRNA} \tag{S 0.11}$$

where, $G_{mRNA}$, $G_{miRNA}^{OFF}$ & $G_{miRNA}^{ON}$ denotes mRNA coding gene, miRNA coding gene in OFF and ON states respectively.

### Mathematical Analysis of The Model

#### 1 Stochastic Master Equation

In our model, there are four stochastic variables, which are the copy number of mRNAs, proteins, miRNAs and mRNA-miRNA complexes, denoted by $r(t), p(t), s(t)$ & $c(t)$ respectively. In addition there is a state variable $i(t)$ describing state of the miRNA coding gene at any time $t$. Let $P^{OFF}_{r,p,s,c}(t) := P(r,p,s,c,i=0,t)$ and $P^{ON}_{r,p,s,c}(t) := P(r,p,s,c,i=1,t)$ be the probabilities that miRNA coding gene is in $OFF$ or $ON$ state, and there are $r$ number of mRNAs, $p$ number of proteins, $s$ number of miRNAs and $c$ number of mRNA-miRNA complexes at any time $t$, respectively. Therefore, $P_{r,p,s,c}(t) := P(r,p,s,c,t) = P^{OFF}_{r,p,s,c,}(t) + P^{ON}_{r,p,s,c,}(t)$ be the probability that there are $r$ number of mRNAs, $p$ number of proteins, $s$ number of miRNAs & $c$ number of complexes at any time $t$, respectively.

The master equation corresponding to $ON$ and $OFF$ states are,

$$\begin{aligned}\frac{\partial P^{ON}_{r,p,s,c}}{\partial t} &= k_r\left[P^{ON}_{r-1,p,s,c} - P^{ON}_{r,p,s,c}\right] + g_r\left[(r+1)P^{ON}_{r+1,p,s,c} - rP^{ON}_{r,p,s,c}\right] + k_p\left[rP^{ON}_{r,p-1,s,c} - rP^{ON}_{r,p,s,c}\right]\\ &+ g_p\left[(p+1)P^{ON}_{r,p+1,s,c} - pP^{ON}_{r,p,s,c}\right] + K_{act}(p+1)P^{OFF}_{r,p+1,s,c} - K_{deact}P^{ON}_{r,p,s,c}\\ &+ k_s\left[P^{ON}_{r,p,s-1,c} - P^{ON}_{r,p,s,c}\right] + g_s\left[(s+1)P^{ON}_{r,p,s+1,c} - sP^{ON}_{r,p,s,c}\right]\\ &+ K_+\left[(r+1)(s+1)P^{ON}_{r+1,p,s+1,c-1} - rsP^{ON}_{r,p,s,c}\right] + k_-\left[(c+1)P^{ON}_{r-1,p,s-1,c+1} - cP^{ON}_{r,p,s,c}\right]\\ &+ \alpha\gamma\left[(c+1)P^{ON}_{r,p,s,c+1} - cP^{ON}_{r,p,s,c}\right] + (1-\alpha)\gamma\left[(c+1)P^{ON}_{r,p,s-1,c+1} - cP^{ON}_{r,p,s,c}\right] \qquad \text{(S 1.1)}\end{aligned}$$

and,

$$\begin{aligned}\frac{\partial P^{OFF}_{r,p,s,c}}{\partial t} &= k_r\left[P^{OFF}_{r-1,p,s,c} - P^{OFF}_{r,p,s,c}\right] + g_r\left[(r+1)P^{OFF}_{r+1,p,s,c} - rP^{OFF}_{r,p,s,c}\right] + k_p\left[rP^{OFF}_{r,p-1,s,c} - rP^{OFF}_{r,p,s,c}\right]\\ &+ g_p\left[(p+1)P^{OFF}_{r,p+1,s,c} - pP^{OFF}_{r,p,s,c}\right] + K_{deact}P^{ON}_{r,p-1,s,c} - K_{act}pP^{OFF}_{r,p,s,c}\\ &+ k^0_s\left[P^{OFF}_{r,p,s-1,c} - P^{OFF}_{r,p,s,c}\right] + g_s\left[(s+1)P^{OFF}_{r,p,s+1,c} - sP^{OFF}_{r,p,s,c}\right]\\ &+ K_+\left[(r+1)(s+1)P^{OFF}_{r+1,p,s+1,c-1} - rsP^{OFF}_{r,p,s,c}\right] + k_-\left[(c+1)P^{OFF}_{r-1,p,s-1,c+1} - cP^{OFF}_{r,p,s,c}\right]\\ &+ \alpha\gamma\left[(c+1)P^{OFF}_{r,p,s,c+1} - cP^{OFF}_{r,p,s,c}\right] + (1-\alpha)\gamma\left[(c+1)P^{OFF}_{r,p,s-1,c+1} - cP^{OFF}_{r,p,s,c}\right] \qquad \text{(S 1.2)}\end{aligned}$$

Therefore, using S 1.1 & S 1.2, we get the chemical master equation as follows,

$$\begin{aligned}\frac{\partial P_{r,p,s,c}}{\partial t} &= \left[\frac{\partial P^{ON}_{r,p,s,c}}{\partial t} + \frac{\partial P^{OFF}_{r,p,s,c}}{\partial t}\right]\\ \implies \frac{\partial P_{r,p,s,c}}{\partial t} &= k_r\left[P_{r-1,p,s,c} - P_{r,p,s,c}\right] + g_r\left[(r+1)P_{r+1,p,s,c} - rP_{r,p,s,c}\right] + k_p\left[rP_{r,p-1,s,c} - rP_{r,p,s,c}\right]\\ &+ g_p\left[(p+1)P_{r,p+1,s,c} - pP_{r,p,s,c}\right] + K_{deact}\left[P^{ON}_{r,p-1,s,c} - P^{ON}_{r,p,s,c}\right]\\ &+ K_{act}\left[(p+1)P^{OFF}_{r,p+1,s,c} - pP^{OFF}_{r,p,s,c}\right] + k^0_s\left[P^{OFF}_{r,p,s-1,c} - P^{OFF}_{r,p,s,c}\right] + k_s\left[P^{ON}_{r,p,s-1,c} - P^{ON}_{r,p,s,c}\right]\\ &+ g_s\left[(s+1)P_{r,p,s+1,c} - sP_{r,p,s,c}\right] + K_+\left[(r+1)(s+1)P_{r+1,p,s+1,c-1} - rsP_{r,p,s,c}\right]\\ &+ k_-\left[(c+1)P_{r-1,p,s-1,c+1} - cP_{r,p,s,c}\right] + \alpha\gamma\left[(c+1)P_{r,p,s,c+1} - cP_{r,p,s,c}\right]\\ &+ (1-\alpha)\gamma\left[(c+1)P_{r,p,s-1,c+1} - cP_{r,p,s,c}\right] \qquad \text{(S 1.3)}\end{aligned}$$

#### 2 Equations of first and second order moments

To explore the miRNA-mediated negative feedback model analytically at the mean and variance level we derive the first & second order moments as defined below,

$$\begin{aligned}\langle r\rangle &= \sum_{r,p,s,c=0}^{\infty} rP_{r,p,s,c} = \sum_{r,p,s,c=0}^{\infty} r\left(P^{ON}_{r,p,s,c} + P^{OFF}_{r,p,s,c}\right) = \langle r\rangle^{ON} + \langle r\rangle^{OFF},\\ \langle r^2\rangle &= \sum_{r,p,s,c=0}^{\infty} r^2P_{r,p,s,c} = \sum_{r,p,s,c=0}^{\infty} r^2\left(P^{ON}_{r,p,s,c} + P^{OFF}_{r,p,s,c}\right) = \langle r^2\rangle^{ON} + \langle r^2\rangle^{OFF},\\ \langle rs\rangle &= \sum_{r,p,s,c=0}^{\infty} rsP_{r,p,s,c} = \sum_{r,p,s,c=0}^{\infty} rs\left(P^{ON}_{r,p,s,c} + P^{OFF}_{r,p,s,c}\right) = \langle rs\rangle^{ON} + \langle rs\rangle^{OFF}\text{etc.}\end{aligned}$$

Multiplying equation S 1.1 by $r$ and summing up over all variables, we get,

$$\begin{aligned}
\sum_{r,p,s,c=0}^{\infty} r\frac{\partial P^{ON}_{r,p,s,c}}{\partial t} &= k_r\left[\sum_{r,p,s,c=0}^{\infty} rP^{ON}_{r-1,p,s,c} - \sum_{r,p,s,c=0}^{\infty} rP^{ON}_{r,p,s,c}\right] + g_r\left[\sum_{r,p,s,c=0}^{\infty} r(r+1)P^{ON}_{r+1,p,s,c} - \sum_{r,p,s,c=0}^{\infty} r^2P^{ON}_{r,p,s,c}\right]\\
&+ k_p\left[\sum_{r,p,s,c=0}^{\infty} r^2P^{ON}_{r,p-1,s,c} - \sum_{r,p,s,c=0}^{\infty} r^2P^{ON}_{r,p,s,c}\right] + g_p\left[\sum_{r,p,s,c=0}^{\infty} r(p+1)P^{ON}_{r,p+1,s,c} - \sum_{r,p,s,c=0}^{\infty} rpP^{ON}_{r,p,s,c}\right]\\
&+ K_{act}\sum_{r,p,s,c=0}^{\infty} r(p+1)P^{OFF}_{r,p+1,s,c} - K_{deact}\sum_{r,p,s,c=0}^{\infty} rP^{ON}_{r,p,s,c}\\
&+ k_s\left[\sum_{r,p,s,c=0}^{\infty} rP^{ON}_{r,p,s-1,c} - \sum_{r,p,s,c=0}^{\infty} rP^{ON}_{r,p,s,c}\right] + g_s\left[\sum_{r,p,s,c=0}^{\infty} r(s+1)P^{ON}_{r,p,s+1,c} - \sum_{r,p,s,c=0}^{\infty} rsP^{ON}_{r,p,s,c}\right]\\
&+ K_+\left[\sum_{r,p,s,c=0}^{\infty} r(r+1)(s+1)P^{ON}_{r+1,p,s+1,c-1} - \sum_{r,p,s,c=0}^{\infty} r^2sP^{ON}_{r,p,s,c}\right]\\
&+ k_-\left[\sum_{r,p,s,c=0}^{\infty} r(c+1)P^{ON}_{r-1,p,s-1,c+1} - \sum_{r,p,s,c=0}^{\infty} rcP^{ON}_{r,p,s,c}\right]\\
&+ \alpha\gamma\left[\sum_{r,p,s,c=0}^{\infty} r(c+1)P^{ON}_{r,p,s,c+1} - \sum_{r,p,s,c=0}^{\infty} rcP^{ON}_{r,p,s,c}\right]\\
&+ (1-\alpha)\gamma\left[\sum_{r,p,s,c=0}^{\infty} r(c+1)P^{ON}_{r,p,s-1,c+1} - \sum_{r,p,s,c=0}^{\infty} rcP^{ON}_{r,p,s,c}\right]
\end{aligned}$$

This simplifies to,

- $$\frac{\partial\langle r\rangle^{ON}}{\partial t} = k_rP^{ON} - g_r\langle r\rangle^{ON} + k_{act}\langle rp\rangle^{OFF} - k_{deact}\langle r\rangle^{ON} - k_+\langle rs\rangle^{ON} + k_-\langle c\rangle^{ON} \quad \text{(S 2.1)}$$

where, $P^{ON} := \sum_{r,p,s,c=0}^{\infty} P^{ON}_{r,p,s,c}$ be the probability that miRNA coding gene is in 'ON' state. Using similar technique, we derive the time-evolution equation for all the first order moments as below,

- $$\frac{\partial\langle r\rangle^{OFF}}{\partial t} = k_rP^{OFF} - g_r\langle r\rangle^{OFF} + k_{deact}\langle r\rangle^{ON} - k_{act}\langle rp\rangle^{OFF} - k_+\langle rs\rangle^{OFF} + k_-\langle c\rangle^{OFF} \quad \text{(S 2.2)}$$

where, $P^{OFF} := \sum_{r,p,s,c=0}^{\infty} P^{OFF}_{r,p,s,c}$ be the probability that miRNA coding gene is in 'OFF' state and $P^{ON}+P^{OFF} = 1$.

- $$\frac{\partial\langle p\rangle^{ON}}{\partial t} = k_p\langle r\rangle^{ON} - g_p\langle p\rangle^{ON} + k_{act}\left[\langle p^2\rangle^{OFF} - \langle p\rangle^{OFF}\right] - k_{deact}\langle p\rangle^{ON} \quad \text{(S 2.3)}$$
- $$\frac{\partial\langle p\rangle^{OFF}}{\partial t} = k_p\langle r\rangle^{OFF} - g_p\langle p\rangle^{OFF} + k_{deact}\left[\langle p\rangle^{ON} + P^{ON}\right] - k_{act}\langle p^2\rangle^{OFF} \quad \text{(S 2.4)}$$
- $$\frac{\partial\langle s\rangle^{ON}}{\partial t} = k_{act}\langle ps\rangle^{OFF} - k_{deact}\langle s\rangle^{ON} + k_sP^{ON} - g_s\langle s\rangle^{ON} - k_+\langle rs\rangle^{ON} + \left[k_- + (1-\alpha)\gamma\right]\langle c\rangle^{ON} \quad \text{(S 2.5)}$$
- $$\begin{aligned}\frac{\partial\langle s\rangle^{OFF}}{\partial t} &= k_{deact}\langle s\rangle^{ON} - k_{act}\langle ps\rangle^{OFF} + k_s^0P^{OFF} - g_s\langle s\rangle^{OFF} - k_+\langle rs\rangle^{OFF}\\ &+ \left[k_- + (1-\alpha)\gamma\right]\langle c\rangle^{OFF}\end{aligned} \quad \text{(S 2.6)}$$
- $$\frac{\partial\langle c\rangle^{ON}}{\partial t} = k_{act}\langle pc\rangle^{OFF} - k_{deact}\langle c\rangle^{ON} + k_+\langle rs\rangle^{ON} - (k_- + \gamma)\langle c\rangle^{ON} \quad \text{(S 2.7)}$$
- $$\frac{\partial\langle c\rangle^{OFF}}{\partial t} = k_{deact}\langle c\rangle^{ON} - k_{act}\langle pc\rangle^{OFF} + k_+\langle rs\rangle^{OFF} - (k_- + \gamma)\langle c\rangle^{OFF} \quad \text{(S 2.8)}$$

Now form S 1.1 summing up in $r, p, s$ & $c$ we get,

$$\begin{aligned}
\sum_{r,p,s,c=0}^{\infty} \frac{\partial P^{ON}_{r,p,s,c}}{\partial t} &= k_r \left[ \sum_{r,p,s,c=0}^{\infty} P^{ON}_{r-1,p,s,c} - \sum_{r,p,s,c=0}^{\infty} P^{ON}_{r,p,s,c} \right] + g_r \left[ \sum_{r,p,s,c=0}^{\infty} (r+1) P^{ON}_{r+1,p,s,c} - \sum_{r,p,s,c=0}^{\infty} r P^{ON}_{r,p,s,c} \right] \\
&+ k_p \left[ \sum_{r,p,s,c=0}^{\infty} r P^{ON}_{r,p-1,s,c} - \sum_{r,p,s,c=0}^{\infty} r P^{ON}_{r,p,s,c} \right] + g_p \left[ \sum_{r,p,s,c=0}^{\infty} (p+1) P^{ON}_{r,p+1,s,c} - \sum_{r,p,s,c=0}^{\infty} p P^{ON}_{r,p,s,c} \right] \\
&+ K_{act} \sum_{r,p,s,c=0}^{\infty} (p+1) P^{OFF}_{r,p+1,s,c} - K_{deact} \sum_{r,p,s,c=0}^{\infty} P^{ON}_{r,p,s,c} + k_s \left[ \sum_{r,p,s,c=0}^{\infty} P^{ON}_{r,p,s-1,c} - \sum_{r,p,s,c=0}^{\infty} P^{ON}_{r,p,s,c} \right] \\
&+ g_s \left[ \sum_{r,p,s,c=0}^{\infty} (s+1) P^{ON}_{r,p,s+1,c} - \sum_{r,p,s,c=0}^{\infty} s P^{ON}_{r,p,s,c} \right] \\
&+ K_+ \left[ \sum_{r,p,s,c=0}^{\infty} (r+1)(s+1) P^{ON}_{r+1,p,s+1,c-1} - \sum_{r,p,s,c=0}^{\infty} rs P^{ON}_{r,p,s,c} \right] \\
&+ k_- \left[ \sum_{r,p,s,c=0}^{\infty} (c+1) P^{ON}_{r-1,p,s-1,c+1} - \sum_{r,p,s,c=0}^{\infty} c P^{ON}_{r,p,s,c} \right] \\
&+ \alpha\gamma \left[ \sum_{r,p,s,c=0}^{\infty} (c+1) P^{ON}_{r,p,s,c+1} - \sum_{r,p,s,c=0}^{\infty} c P^{ON}_{r,p,s,c} \right] \\
&+ (1-\alpha)\gamma \left[ \sum_{r,p,s,c=0}^{\infty} (c+1) P^{ON}_{r,p,s-1,c+1} - \sum_{r,p,s,c=0}^{\infty} c P^{ON}_{r,p,s,c} \right] \\
\implies \frac{\partial P^{ON}}{\partial t} &= k_{act} < p >^{OFF} - k_{deact} P^{ON} \qquad \text{(S 2.9)}
\end{aligned}$$

Equation S 2.9 describes the time-evolution of the probability of miRNA coding gene being in the 'ON' state and equations S 2.1 - S 2.8 expresses the time-evolution of first order moments of different variables for both 'ON' and 'OFF' states of miRNA coding gene.

Now from S 1.1 and S 1.2, using similar techniques, we can derive the time-evolution of second order moments as below,

- $$\begin{aligned}\frac{\partial \langle r^2 \rangle^{ON}}{\partial t} &= k_r \left[ 2\langle r \rangle^{ON} + P^{ON} \right] + g_r \left[ \langle r \rangle^{ON} - 2\langle r^2 \rangle^{ON} \right] + k_{act} \langle r^2 p \rangle^{OFF} - k_{deact} \langle r^2 \rangle^{ON} \\ &+ k_+ \left[ \langle rs \rangle^{ON} - 2\langle r^2 s \rangle^{ON} \right] + k_- \left[ \langle c \rangle^{ON} + 2\langle rc \rangle^{ON} \right] \qquad \text{(S 2.10)}\end{aligned}$$
- $$\begin{aligned}\frac{\partial \langle r^2 \rangle^{OFF}}{\partial t} &= k_r \left[ 2\langle r \rangle^{OFF} + P^{OFF} \right] + g_r \left[ \langle r \rangle^{OFF} - 2\langle r^2 \rangle^{OFF} \right] - k_{act} \langle r^2 p \rangle^{OFF} \\ &+ k_{deact} \langle r^2 \rangle^{ON} + k_+ \left[ \langle rs \rangle^{OFF} - 2\langle r^2 s \rangle^{OFF} \right] + k_- \left[ \langle c \rangle^{OFF} + 2\langle rc \rangle^{OFF} \right] \qquad \text{(S 2.11)}\end{aligned}$$
- $$\begin{aligned}\frac{\partial \langle p^2 \rangle^{ON}}{\partial t} &= k_p \left[ \langle r \rangle^{ON} + 2\langle rp \rangle^{ON} \right] + g_p \left[ \langle p \rangle^{ON} - 2\langle p^2 \rangle^{ON} \right] + k_{act} \left[ \langle p^3 \rangle^{OFF} - 2\langle p^2 \rangle^{OFF} + \langle p \rangle^{OFF} \right] \\ &- k_{deact} \langle p^2 \rangle^{ON} \qquad \text{(S 2.12)}\end{aligned}$$
- $$\begin{aligned}\frac{\partial \langle p^2 \rangle^{OFF}}{\partial t} &= k_p \left[ \langle r \rangle^{OFF} + 2\langle rp \rangle^{OFF} \right] + g_p \left[ \langle p \rangle^{OFF} - 2\langle p^2 \rangle^{OFF} \right] + k_{deact} \left[ \langle p^2 \rangle^{ON} + 2\langle p \rangle^{ON} + P^{ON} \right] \\ &- k_{act} \langle p^3 \rangle^{OFF} \qquad \text{(S 2.13)}\end{aligned}$$
- $$\begin{aligned}\frac{\partial \langle s^2 \rangle^{ON}}{\partial t} &= k_{act} \langle ps^2 \rangle^{OFF} - k_{deact} \langle s^2 \rangle^{ON} + k_s \left[ 2\langle s \rangle^{ON} + P^{ON} \right] + g_s \left[ \langle s \rangle^{ON} - 2\langle s^2 \rangle^{ON} \right] \\ &+ k_+ \left[ \langle rs \rangle^{ON} - 2\langle rs^2 \rangle^{ON} \right] + \left[ k_- + (1-\alpha)\gamma \right] \left[ \langle c \rangle^{ON} + 2\langle sc \rangle^{ON} \right] \qquad \text{(S 2.14)}\end{aligned}$$
- $$\begin{aligned}\frac{\partial \langle s^2 \rangle^{OFF}}{\partial t} &= k_{deact} \langle s^2 \rangle^{ON} - k_{act} \langle ps^2 \rangle^{OFF} + k_s^0 \left[ 2\langle s \rangle^{OFF} + P^{OFF} \right] + g_s \left[ \langle s \rangle^{OFF} - 2\langle s^2 \rangle^{OFF} \right] \\ &+ k_+ \left[ \langle rs \rangle^{OFF} - 2\langle rs^2 \rangle^{OFF} \right] + \left[ k_- + (1-\alpha)\gamma \right] \left[ \langle c \rangle^{OFF} + 2\langle sc \rangle^{OFF} \right] \qquad \text{(S 2.15)}\end{aligned}$$
- $$\begin{aligned}\frac{\partial \langle c^2 \rangle^{ON}}{\partial t} &= k_{act} \langle pc^2 \rangle^{OFF} - k_{deact} \langle c^2 \rangle^{ON} + k_+ \left[ \langle rs \rangle^{ON} + 2\langle rsc \rangle^{ON} \right] \\ &+ (k_- + \gamma) \left[ \langle c \rangle^{ON} - 2\langle c^2 \rangle^{ON} \right] \qquad \text{(S 2.16)}\end{aligned}$$
- $$\begin{aligned}\frac{\partial \langle c^2 \rangle^{OFF}}{\partial t} &= k_{deact} \langle c^2 \rangle^{ON} - k_{act} \langle pc^2 \rangle^{OFF} + k_+ \left[ \langle rs \rangle^{OFF} + 2\langle rsc \rangle^{OFF} \right] \\ &+ (k_- + \gamma) \left[ \langle c \rangle^{OFF} - 2\langle c^2 \rangle^{OFF} \right] \qquad \text{(S 2.17)}\end{aligned}$$

Now multiplying $rs$ with S 1.1 and then summing up in $r, p, s$ & $c$ we derive the second order correlations as below,

- $$\frac{\partial \langle rs \rangle^{ON}}{\partial t} = k_r \langle s \rangle^{ON} - g_r \langle rs \rangle^{ON} + k_s \langle r \rangle^{ON} - g_s \langle rs \rangle^{ON} + k_{act} \langle rps \rangle^{OFF} - k_{deact} \langle rs \rangle^{ON} + k_+ \left[ \langle rs \rangle^{ON} - \langle r^2 s \rangle^{ON} - \langle rs^2 \rangle^{ON} \right] + k_- \left[ \langle rc \rangle^{ON} + \langle sc \rangle^{ON} + \langle c \rangle^{ON} \right] + (1-\alpha)\gamma \langle rc \rangle^{ON} \quad \text{(S 2.18)}$$

and from S 1.2 we get,

- $$\frac{\partial \langle rs \rangle^{OFF}}{\partial t} = k_r \langle s \rangle^{OFF} - g_r \langle rs \rangle^{OFF} + k_s^0 \langle r \rangle^{OFF} - g_s \langle rs \rangle^{OFF} + k_{deact} \langle rs \rangle^{ON} - k_{act} \langle rps \rangle^{OFF} + k_+ \left[ \langle rs \rangle^{OFF} - \langle r^2 s \rangle^{OFF} - \langle rs^2 \rangle^{OFF} \right] + k_- \left[ \langle rc \rangle^{OFF} + \langle sc \rangle^{OFF} + \langle c \rangle^{OFF} \right] + (1-\alpha)\gamma \langle rc \rangle^{OFF} \quad \text{(S 2.19)}$$

Similarly,

- $$\frac{\partial \langle rc \rangle^{ON}}{\partial t} = k_r \langle c \rangle^{ON} - g_r \langle rc \rangle^{ON} + k_{act} \langle rpc \rangle^{OFF} - k_{deact} \langle rc \rangle^{ON} + k_+ \left[ \langle r^2 s \rangle^{ON} - \langle rsc \rangle^{ON} - \langle rs \rangle^{ON} \right] + k_- \left[ \langle c^2 \rangle^{ON} - \langle c \rangle^{ON} - \langle rc \rangle^{ON} \right] - \gamma \langle rc \rangle^{ON} \quad \text{(S 2.20)}$$
- $$\frac{\partial \langle rc \rangle^{OFF}}{\partial t} = k_r \langle c \rangle^{OFF} - g_r \langle rc \rangle^{OFF} - k_{act} \langle rpc \rangle^{OFF} + k_{deact} \langle rc \rangle^{ON} + k_+ \left[ \langle r^2 s \rangle^{OFF} - \langle rsc \rangle^{OFF} - \langle rs \rangle^{OFF} \right] + k_- \left[ \langle c^2 \rangle^{OFF} - \langle c \rangle^{OFF} - \langle rc \rangle^{OFF} \right] - \gamma \langle rc \rangle^{OFF} \quad \text{(S 2.21)}$$
- $$\frac{\partial \langle sc \rangle^{ON}}{\partial t} = k_s \langle c \rangle^{ON} - (g_s + \alpha\gamma) \langle sc \rangle^{ON} + k_{act} \langle psc \rangle^{OFF} - k_{deact} \langle sc \rangle^{ON} + k_+ \left[ \langle rs^2 \rangle^{ON} - \langle rsc \rangle^{ON} - \langle rs \rangle^{ON} \right] + [k_- + (1-\alpha)\gamma] \left[ \langle c^2 \rangle^{ON} - \langle c \rangle^{ON} - \langle sc \rangle^{ON} \right] \quad \text{(S 2.22)}$$
- $$\frac{\partial \langle sc \rangle^{OFF}}{\partial t} = k_s^0 \langle c \rangle^{OFF} - (g_s + \alpha\gamma) \langle sc \rangle^{OFF} + k_{deact} \langle sc \rangle^{ON} - k_{act} \langle psc \rangle^{OFF} + k_+ \left[ \langle rs^2 \rangle^{OFF} - \langle rsc \rangle^{OFF} - \langle rs \rangle^{OFF} \right] + [k_- + (1-\alpha)\gamma] \left[ \langle c^2 \rangle^{OFF} - \langle c \rangle^{OFF} - \langle sc \rangle^{OFF} \right] \quad \text{(S 2.23)}$$
- $$\frac{\partial \langle rp \rangle^{ON}}{\partial t} = k_r \langle p \rangle^{ON} - g_r \langle rp \rangle^{ON} + k_p \langle r^2 \rangle^{ON} - g_p \langle rp \rangle^{ON} - k_{deact} \langle rp \rangle^{ON} + k_{act} \left[ \langle rp^2 \rangle^{OFF} - \langle rp \rangle^{OFF} \right] - k_+ \langle rps \rangle^{ON} + k_- \langle pc \rangle^{ON} \quad \text{(S 2.24)}$$
- $$\frac{\partial \langle rp \rangle^{OFF}}{\partial t} = k_r \langle p \rangle^{OFF} - g_r \langle rp \rangle^{OFF} + k_p \langle r^2 \rangle^{OFF} - g_p \langle rp \rangle^{OFF} - k_{act} \langle rp^2 \rangle^{OFF} + k_{deact} \left[ \langle rp \rangle^{ON} + \langle r \rangle^{ON} \right] - k_+ \langle rps \rangle^{OFF} + k_- \langle pc \rangle^{OFF} \quad \text{(S 2.25)}$$
- $$\frac{\partial \langle ps \rangle^{ON}}{\partial t} = k_p \langle rs \rangle^{ON} - g_p \langle ps \rangle^{ON} + k_s \langle p \rangle^{ON} - g_s \langle ps \rangle^{ON} - k_{deact} \langle ps \rangle^{ON} + k_{act} \left[ \langle p^2 s \rangle^{OFF} - \langle ps \rangle^{OFF} \right] - k_+ \langle rps \rangle^{ON} + [k_- + (1-\alpha)\gamma] \langle pc \rangle^{ON} \quad \text{(S 2.26)}$$
- $$\frac{\partial \langle ps \rangle^{OFF}}{\partial t} = k_p \langle rs \rangle^{OFF} - g_p \langle ps \rangle^{OFF} + k_s^0 \langle p \rangle^{OFF} - g_s \langle ps \rangle^{OFF} - k_{act} \langle p^2 s \rangle^{OFF} + k_{deact} \left[ \langle s \rangle^{ON} + \langle ps \rangle^{ON} \right] - k_+ \langle rps \rangle^{OFF} + [k_- + (1-\alpha)\gamma] \langle pc \rangle^{OFF} \quad \text{(S 2.27)}$$
- $$\frac{\partial \langle pc \rangle^{ON}}{\partial t} = k_p \langle rc \rangle^{ON} - [g_p + k_- + \gamma] \langle pc \rangle^{ON} - k_{deact} \langle pc \rangle^{ON} + k_{act} \left[ \langle p^2 c \rangle^{OFF} - \langle pc \rangle^{OFF} \right] + k_+ \langle rps \rangle^{ON} \quad \text{(S 2.28)}$$
- $$\frac{\partial \langle pc \rangle^{OFF}}{\partial t} = k_p \langle rc \rangle^{OFF} - [g_p + k_- + \gamma] \langle pc \rangle^{OFF} + k_{deact} \left[ \langle pc \rangle^{ON} + \langle c \rangle^{ON} \right] - k_{act} \langle p^2 c \rangle^{OFF} + k_+ \langle rps \rangle^{OFF} \quad \text{(S 2.29)}$$

Note that, each moment equation of a stochastic variable contains the next higher order moments, creating an infinite hierarchy. Therefore the equations do not form a closed system. To break the hierarchy we assume two kind of approximations, namely,

➤ mean-field approximation (**MFA**), where $2nd$ order moments are expressed in terms of $1st$ order moments

➤ third order moment approximation (**TOMA**), , where $3rd$ order moments are expressed in terms of lower order moments

This is discussed in the later sections.

### 3 Mean-Field Approximation

First order moment (**FOM**) of variables $r(t), p(t), s(t)$ & $c(t)$ describe the average copy number of mRNAs, proteins, miRNAs & mRNA-miRNA complexes, respectively, at time t. From equations S 2.1 - S 2.9 we have the following,

$$\textbf{FOM 1}: \frac{\partial\langle r\rangle}{\partial t} = \frac{\partial\langle r\rangle^{ON}}{\partial t} + \frac{\partial\langle r\rangle^{OFF}}{\partial t} = k_r - g_r\langle r\rangle - k_+\langle rs\rangle + k_-\langle c\rangle$$

$$\textbf{FOM 2}: \frac{\partial\langle p\rangle}{\partial t} = \frac{\partial\langle p\rangle^{ON}}{\partial t} + \frac{\partial\langle p\rangle^{OFF}}{\partial t} = k_p\langle r\rangle - g_p\langle p\rangle - k_{act}\langle p\rangle^{OFF} + k_{deact}P^{ON}$$

$$\textbf{FOM 3}: \frac{\partial\langle s\rangle}{\partial t} = \frac{\partial\langle s\rangle^{ON}}{\partial t} + \frac{\partial\langle s\rangle^{OFF}}{\partial t} = \left[k_sP^{ON} + k_s^0P^{OFF}\right] - g_s\langle s\rangle - k_+\langle rs\rangle + \left[k_- + (1-\alpha)\gamma\right]\langle c\rangle$$

$$\textbf{FOM 4}: \frac{\partial\langle c\rangle}{\partial t} = \frac{\partial\langle c\rangle^{ON}}{\partial t} + \frac{\partial\langle c\rangle^{OFF}}{\partial t} = k_+\langle rs\rangle - \left[k_- + \gamma\right]\langle c\rangle$$

$$\textbf{FOM 5}: \frac{\partial P^{ON}}{\partial t} = k_{act}\langle p\rangle^{OFF} - k_{deact}P^{ON} \qquad \left[where, P^{ON} + P^{OFF} = 1\right]$$

which describe the time evolution of mean mRNA($\langle r\rangle$), protein($\langle p\rangle$), miRNA($\langle s\rangle$) and mRNA-miRNA complex($\langle c\rangle$) copy number. The equation **FOM 5** is to be considered along with **FOM 1 - 4** in order to solve for $\langle r\rangle, \langle p\rangle, \langle s\rangle$ & $\langle c\rangle$, since $P^{ON}$ is present in **FOM 2** and **FOM 3**

The only second order moment present in **FOM 1 - 5** is $\langle rs\rangle$. Also, $\langle p\rangle^{OFF}$ be another variable that breaks the Closure of the system of equations **FOM 1 - 5**. So we define the mean-field approximations (**MFA**) as below to express $\langle rs\rangle$ and $\langle p\rangle^{OFF}$ in terms of first order moments.

$$\textbf{MFA I.}\ \langle (r - <r>)(s - <s>)\rangle = 0 \implies <rs> = <r><s>$$

$$\textbf{MFA II.}\ \langle (p - <p>)\rangle^{OFF} = 0 \implies <p>^{OFF} = <p>P^{OFF} = <p>\left(1 - P^{ON}\right)$$

we get from **FOM 1 - 5** the following,

$$\frac{\partial\langle r\rangle}{\partial t} \approx k_r - g_r\langle r\rangle - k_+\langle r\rangle\langle s\rangle + k_-\langle c\rangle \tag{S 3.1}$$

$$\frac{\partial\langle p\langle}{\partial t} \approx k_p\langle r\rangle - g_p\langle p\rangle - k_{act}\langle p\rangle\left(1 - P^{ON}\right) + k_{deact}P^{ON} \tag{S 3.2}$$

$$\frac{\partial\langle s\rangle}{\partial t} \approx \left[k_sP^{ON} + k_s^0P^{OFF}\right] - g_s\langle s\rangle - k_+\langle rs\rangle + \left[k_- + (1-\alpha)\gamma\right]\langle c\rangle \tag{S 3.3}$$

$$\frac{\partial\langle c\rangle}{\partial t} \approx k_+\langle r\rangle\langle s\rangle - \left[k_- + \gamma\right]\langle c\rangle \tag{S 3.4}$$

$$\frac{\partial P^{ON}}{\partial t} \approx k_{act}\langle p\rangle\left(1 - P^{ON}\right) - k_{deact}P^{ON} \tag{S 3.5}$$

which we shall refer as Mean-Field Equations (**MFE**).

#### 4 Steady state solution of Mean-Field Approximated Equations

From S 3.2 & S 3.5 at steady state,

$$\langle r\rangle k_p - g_p\langle p\rangle = 0$$
$$\implies \langle p\rangle = \frac{k_p}{g_p}\langle r\rangle \tag{S 4.1}$$

From S 3.1 & S 3.4 at steady state,

$$k_r - g_r\langle r\rangle - \gamma\langle c\rangle = 0$$
$$\implies \langle c\rangle = \frac{1}{\gamma}\Big[k_r - g_r\langle r\rangle\Big] \tag{S 4.2}$$

From S 3.4 using S 4.2 at steady state, we get,

$$k_+\langle r\rangle\langle s\rangle = (k_- + \gamma)\,\langle c\rangle$$
$$\implies \langle s\rangle = \frac{(k_- + \gamma)\,\langle c\rangle}{k_+\langle r\rangle} = \frac{(k_- + \gamma)}{k_+\langle r\rangle}\frac{[k_r - g_r\langle r\rangle]}{\gamma} \tag{S 4.3}$$

From S 3.5 using S 4.1 at steady state, we get,

$$P^{ON} = \frac{k_{act}\langle p\rangle}{(k_{act}\langle p\rangle + k_{deact})}$$
$$\implies P^{ON} = \frac{k_p k_{act}\langle r\rangle}{(k_p k_{act}\langle r\rangle + g_p k_{deact})} \tag{S 4.4}$$

Now in S 3.3 using S 4.4 at steady state we get,

$$\begin{aligned}
&\frac{(k_s - k_s^0)k_p k_{act}\langle r\rangle}{(k_p k_{act}\langle r\rangle + g_p k_{deact})} - g_s\langle s\rangle - \alpha\gamma\langle c\rangle + k_s^0 &&= 0 \\
\implies & k_s k_p k_{act}\langle r\rangle + k_s^0 g_p k_{deact} - \Big[g_s\langle s\rangle + \alpha\big(k_r - g_r\langle r\rangle\big)\Big]\Big[k_p k_{act}\langle r\rangle + g_p k_{deact}\Big] &&= 0 \quad \text{[using S 4.2]} \\
\implies & k_s k_p k_{act}\langle r\rangle + k_s^0 g_p k_{deact} - g_s\frac{(k_- + \gamma)}{k_+\langle r\rangle}\frac{[k_r - g_r\langle r\rangle]}{\gamma}\Big[k_p k_{act}\langle r\rangle + g_p k_{deact}\Big] && \\
& \qquad - \alpha\big(k_r - g_r\langle r\rangle\big)\Big[k_p k_{act}\langle r\rangle + g_p k_{deact}\Big] &&= 0 \quad \text{[using S 4.3]}
\end{aligned}$$

let $g = \frac{k_+\gamma}{(k_- + \gamma)}$, be the 'effective association rate' between mRNAs and miRNAs. From above equation we get,

$$\begin{aligned}
&\alpha g g_r k_p k_{act}\langle r\rangle^3 + \Big[\alpha g\left(g_r g_p k_{deact} - k_r k_p k_{act}\right) + \left(g k_s + g_s g_r\right)k_p k_{act}\Big]\langle r\rangle^2 \\
&\qquad + \Big[\big\{g_s g_r + g\left(k_s^0 - \alpha k_r\right)\big\}g_p k_{deact} - g_s k_r k_p k_{act}\Big]\langle r\rangle - g_s k_r g_p k_{deact} = 0
\end{aligned}$$

In our study we focus on $k_+ \gtrapprox k_-$ (i.e, tight binding between mRNA & mRNA), since these are biologically relevant conditions under which target mRNA gene expression can be regulated by miRNAs. Therefore, the effective association rate $O(g) \gtrapprox O(k_r, k_p, k_+)$

Then steady state mean mRNA copy number $\langle r\rangle$ is governed by the following equation,

$$A\langle r\rangle^3 + B\langle r\rangle^2 + C\langle r\rangle + D = 0 \tag{S 4.5}$$

$A = \alpha g g_r k_p k_{act}$; $B = \Big[\alpha g\left(g_r g_p k_{deact} - k_r k_p k_{act}\right) + \left(g k_s + g_s g_r\right)k_p k_{act}\Big]$;
$C = \Big[\big\{g_s g_r + g\left(k_s^0 - \alpha k_r\right)\big\}g_p k_{deact} - g_s k_r k_p k_{act}\Big]$; $D = -g_s k_r g_p k_{deact}$

To study the effect fo feedback strength ($\beta = k_{act}/k_{deact}$) at the mean level we varied the miRNA gene activation strength ($k_{act}$) keeping the deactivation strength ($k_{deact}$) fixed. So, two cases arise, (i) no feedback ($k_{act} = 0s^{-1}$) and (ii) with feedback ($k_{act} \neq 0s^{-1}$). These two cases are illustrated next.

#### 4.1 Case I. No Feedback

when there is no activation by protein molecules (i.e, $k_{act} = 0s^{-1}$), we have $A = 0$ and S 4.5 becomes,

$$B\langle r\rangle^2 + C\langle r\rangle + D = 0$$

where, $B = \alpha g g_r g_p k_{deact}$; $C = \{g_s g_r + g\left(k_s^0 - \alpha k_r\right)\} g_p k_{deact}$; $D = -g_s k_r g_p k_{deact}$

Solving the above equation we get,

$$\langle r\rangle = -\frac{C}{2B} \pm \sqrt{\left(\frac{C}{2B}\right)^2 - \frac{D}{B}}$$

where,

$$\left.\frac{C}{B} = \frac{g_s g_r + g\left(k_s^0 - \alpha k_r\right)}{\alpha g g_r}; \qquad \frac{D}{B} = -\frac{g_s k_r}{\alpha g g_r} \right\} \implies \left(\frac{C}{2B}\right)^2 - \frac{D}{B} \geq 0$$

Therefore steady state mean mRNA copy number is given by,

$$\langle r\rangle = -\frac{C}{2B} + \sqrt{\left(\frac{C}{2B}\right)^2 - \frac{D}{B}}$$

$$\implies \langle r\rangle = -\frac{g_s g_r + g\left(k_s^0 - \alpha k_r\right)}{2\alpha g g_r} + \sqrt{\left\{\frac{g_s g_r + g\left(k_s^0 - \alpha k_r\right)}{2\alpha g g_r}\right\}^2 + \frac{g_s k_r}{\alpha g g_r}} \tag{S 4.6}$$

#### 4.2 Case II. with Feedback

When $k_{act} \neq 0s^{-1}$, we have $A \neq 0$ and using Cardano's method of solving 3rd order polynomial equation S 4.5 we get,

$$\langle r\rangle = \left[-\frac{L}{2} + \sqrt{\left(\frac{L}{2}\right)^2 + \left(\frac{K}{3}\right)^3}\right]^{\frac{1}{3}} - \left[\frac{L}{2} + \sqrt{\left(\frac{L}{2}\right)^2 + \left(\frac{K}{3}\right)^3}\right]^{\frac{1}{3}} - \frac{B}{3A} \tag{S 4.7}$$

where, $L = \left[\left(\frac{D}{A}\right) - \frac{1}{3}\left(\frac{B}{A}\right)\left(\frac{C}{A}\right) + \frac{2}{27}\left(\frac{B}{A}\right)^2\right]$ & $K = \left[\left(\frac{C}{A}\right) - \frac{1}{3}\left(\frac{B}{A}\right)^2\right]$. Here $A, B, C$ & $D$ are same as in S 4.5.

Therefore, the steady state mean mRNA copy number is given by either S 4.6 or S 4.7.

### 5 Threshold behavior in target mRNA

we have $g = \frac{<c>\gamma}{<r><s>} = \frac{k_+\gamma}{(k_-+\gamma)}$ [from S 4.3]. Then from equation S 3.1 & S 3.4 at steady state we have,

$$k_r - \langle r\rangle g_r - g\langle r\rangle\langle s\rangle = 0$$
$$k_s P^{ON} + k_s^0\left(1 - P^{ON}\right) - \langle s\rangle g_s - \alpha g\langle r\rangle\langle s\rangle = 0$$

After rearranging we get,

$$\langle r\rangle = \frac{1}{2\alpha g g_r}\Bigg[ -\Big[g_r g_s + g\big\{k_s P^{ON} + k_s^0\left(1 - P^{ON}\right)\big\} - \alpha g k_r\Big] \pm \sqrt{\Big[g_r g_s + g\big\{k_s P^{ON} + k_s^0\left(1 - P^{ON}\right)\big\} - \alpha g k_r\Big]^2 + 4\alpha g k_r g_r g_s}\ \Bigg]$$

and $$\langle s\rangle = \frac{k_s P^{ON} + k_s^0\left(1 - p^{ON}\right)}{g_s + \alpha g\langle r\rangle}$$

In the limit $g \to \infty$ i.e, $k_+ >> (k_- + \gamma)$ we have,

$$\langle r\rangle = \frac{k_r}{2g_r} - \frac{k_s P^{ON} + k_s^0\left(1 - P^{ON}\right)}{2\alpha g_r} \pm \left|\frac{k_r}{2g_r} - \frac{k_s P^{ON} + k_s^0\left(1 - P^{ON}\right)}{2\alpha g_r}\right|$$

$$\implies \langle r\rangle = \begin{cases} \frac{k_r}{g_r} - \frac{k_s P^{ON} + k_s^0\left(1-P^{ON}\right)}{\alpha g_r} & when \quad k_r \geq \frac{k_s P^{ON} + k_s^0\left(1-P^{ON}\right)}{\alpha} \\ 0 & when \quad k_r < \frac{k_s P^{ON} + k_s^0\left(1-P^{ON}\right)}{\alpha} \end{cases} \tag{S 5.1}$$

Now according to S 4.4,

$$P^{ON} = \begin{cases} 1 & when \quad k_{act} \to \infty \\ 0 & when \quad k_{act} = 0 \end{cases} \tag{S 5.2}$$

Therefore, from S 5.1 using S 5.2 in the limit $g \to \infty$ & $k_{act} = 0$ we have mRNA copy number as follows,

$$\langle r\rangle = \begin{cases} \frac{\alpha k_r - k_s^0}{\alpha g_r} & when \quad k_r \geq \frac{k_s^0}{\alpha} \\ 0 & when \quad k_r < \frac{k_s^0}{\alpha} \end{cases} \tag{S 5.3}$$

and in the limit $g \to \infty$ & $k_{act} \to \infty$ we have,

$$\langle r\rangle = \begin{cases} \frac{\alpha k_r - k_s}{\alpha g_r} & when \quad k_r \geq \frac{k_s}{\alpha} \\ 0 & when \quad k_r < \frac{k_s}{\alpha} \end{cases} \tag{S 5.4}$$

Hence in our miRNA mediated negative feedback model, when the effective association rate is high (i.e, $g \to \infty$) miRNAs are capable of generating a threshold linear behaviour in the target mRNA expression at mean level. S 5.3 and S 5.4 suggest that, $k_r = \frac{k_s^0}{\alpha}$ and $k_r = \frac{k_s}{\alpha}$ are the mRNA transcription rate threshold values for steady state mean mRNA corresponding to the cases of no activation ($k_{act} = 0s^{-1}$) and string activation strength ($k_{act} \to \infty$).

Now we try to find a general threshold for mean mRNA expression for any nonzero activation strength ($k_{act} \neq 0$) which is illustrated next.

#### 5.1 General threshold for mean mRNA copy number

##### 5.1.1 General $k_r$ threshold

Considering mRNA transcription rate ($k_r$) as a independent variable, equations S 4.6 & S 4.7 suggests that mean mRNA copy number can be thought of a function of $k_r$, i.e, $\langle r\rangle = f(k_r)$, where $f$ is the right hand side of equations S 4.6 & S 4.7 corresponding to the cases of $k_{act} = 0$ & $k_{act} \to \infty$. Now in the $\langle r\rangle$vs $k_r$ plane the mean mRNA curve $f(x)$ has a slope defined for every value of $x \equiv k_r$. Keeping the shapes of the functions S 5.3 & S 5.4 in mind, to define the threshold in a general sense, a specific $k_r$ value is considered, where the change in slope of $f(k_r)$ is maximum, and we call this $k_r$ value as $k_r^{slope}$. Hence we have,

$$\begin{aligned}
&\langle r\rangle = f(k_r) \\
\implies &\frac{df(k_r)}{dk_r} \equiv f'(k_r) = \text{slope of } f(k_r) \text{at any } \ k_r \text{value} \\
\implies &f''(k_r) = \text{change in slope of } f(k_r) \text{ at any } k_r \text{value} \\
\implies &f'''(k_r) = 0, \text{ gives } k_r = k_r^{slope}, \text{where the change in slope is maximun}
\end{aligned}$$

In the $\langle r\rangle$vs $k_r$ plane the tangent to the mean mRNA curve ($\langle r\rangle \equiv f(k_r)$) at $k_r = k_r^{slope}$ will eventually intersect the $k_r$ axis. We define this point of intersection as the threshold and denote it by $k_r^{th}$.Therefore,

$$\begin{aligned}
&< r >= f(k_r^{slope}) + f'(k_r^{slope})\left(k_r - k_r^{slope}\right) \text{ be the tangent} \\
&\qquad\qquad \text{to mean mRNA curve at } k_r = k_r^{slope} \text{ in } < r > \text{ vs } k_r \text{ plane} \\
&\implies k_r^{th} f'(k_r^{slope}) + \left[f(k_r^{slope} - k_r^{slope} f'(k_r^{slope}))\right] = 0 \text{ gives the point of intersection} \\
&\qquad\qquad \text{of the tangent to the mRNA curve and } k_r \text{ axis} \\
&\implies k_r^{th} = k_r^{slope} - \frac{f(k_r^{slope})}{f'(k_r^{slope})} \text{ be the general } k_r \text{ threshold.}
\end{aligned}$$

- **Explicit expression of $k_r^{th}$ for nonzero $k_{act}$:**

Since we are focusing on the limit $g \to \infty$ & form equation S 4.7 it is very difficult to calculate $\lim_{g\to\infty}\langle r\rangle$, so we derive the threshold using S 4.5. In presence of activation by protein molecule, the mean mRNA copy number is given as follows,

$$\begin{aligned}
&\langle r\rangle = -\frac{B}{2A} \pm \sqrt{\left(\frac{B}{2A}\right)^2 - \frac{C}{A}} \\
&\implies \langle r\rangle = f(k_r) = -\frac{1}{2}\left[\frac{(k_s - \alpha k_r)}{\alpha g_r} + \frac{g_p k_{deact}}{k_p k_{act}}\right] \pm \sqrt{\frac{1}{4}\left\{\frac{(k_s - \alpha k_r)}{\alpha g_r} + \frac{g_p k_{deact}}{k_p k_{act}}\right\}^2 - \frac{g_p k_{deact}\,(k_s^0 - \alpha k_r)}{\alpha g_r k_p k_{act}}}
\end{aligned}$$

since,

$$\frac{B}{A} \to \left[\frac{(k_s - \alpha k_r)}{\alpha g_r} + \frac{g_p k_{deact}}{k_p k_{act}}\right]; \qquad \frac{C}{A} \to \frac{g_p k_{deact}\left(k_s^0 - \alpha k_r\right)}{\alpha g_r k_p k_{act}} \qquad \& \qquad \frac{D}{A} \to 0 \qquad as \qquad g \to \infty$$

Now,

$$\begin{aligned}
&\left(\frac{B}{2A}\right)^2 - \frac{C}{A} = 0 \\
&\implies k_r = \left(\frac{k_s}{\alpha} - \frac{g_r g_p k_{deact}}{k_p k_{act}}\right) \pm \sqrt{4\left(\frac{k_s^0 - k_s}{\alpha}\right)\frac{g_r g_p k_{deact}}{k_p k_{act}}}
\end{aligned}$$

so $\left(\frac{B}{2A}\right)^2 - \frac{C}{A} = 0$ only at imaginary $k_r$ values, since $k_s^0 < k_s$. Therefore $\left(\frac{B}{2A}\right)^2 - \frac{C}{A} > 0$ and hence in the limit $g \to \infty$ we have

$$\langle r\rangle = f(k_r) = -\frac{1}{2}\left[\frac{(k_s - \alpha k_r)}{\alpha g_r} + \frac{g_p k_{deact}}{k_p k_{act}}\right] + \sqrt{\frac{1}{4}\left\{\frac{(k_s - \alpha k_r)}{\alpha g_r} + \frac{g_p k_{deact}}{k_p k_{act}}\right\}^2 - \frac{g_p k_{deact}\,(k_s^0 - \alpha k_r)}{\alpha g_r k_p k_{act}}} \qquad \text{(S 5.5)}$$

then,

$$f'(k_r) = \frac{1}{2g_r} + \frac{\frac{1}{4g_r}\left\{\frac{g_p k_{deact}}{k_p k_{act}} - \frac{(k_s-\alpha k_r)}{\alpha g_r}\right\}}{\sqrt{\frac{1}{4}\left\{\frac{(k_s-\alpha k_r)}{\alpha g_r} + \frac{g_p k_{deact}}{k_p k_{act}}\right\}^2 - \frac{g_p k_{deact}(k_s^0-\alpha k_r)}{\alpha g_r k_p k_{act}}}}$$

$$\implies f''(k_r) = \frac{\left(\frac{1}{2g_r}\right)^2 \left(\frac{g_p k_{deact}}{k_p k_{act}}\right)\left(\frac{k_s-k_s^0}{\alpha g_r}\right)}{\left\{\frac{1}{4}\left\{\frac{(k_s-\alpha k_r)}{\alpha g_r} + \frac{g_p k_{deact}}{k_p k_{act}}\right\}^2 - \frac{g_p k_{deact}(k_s^0-\alpha k_r)}{\alpha g_r k_p k_{act}}\right\}^{\frac{3}{2}}}$$

$$\implies f'''(k_r) = -\frac{\frac{3}{2}\left(\frac{1}{2g_r}\right)^3 \left(\frac{g_p k_{deact}}{k_p k_{act}}\right)\left(\frac{k_s-k_s^0}{\alpha g_r}\right)\left[\frac{g_p k_{deact}}{k_p k_{act}} - \frac{(k_s-\alpha k_r)}{\alpha g_r}\right]}{\left[\frac{1}{4}\left\{\frac{(k_s-\alpha k_r)}{\alpha g_r} + \frac{g_p k_{deact}}{k_p k_{act}}\right\}^2 - \frac{g_p k_{deact}(k_s^0-\alpha k_r)}{\alpha g_r k_p k_{act}}\right]^{\frac{5}{2}}}$$

Therefore, $k_r^{slope}$ is given by,

$$f'''(k_r^{slope}) = 0$$
$$\implies k_r^{slope} = \left[\frac{k_s}{\alpha} - \frac{g_r g_p k_{deact}}{k_p k_{act}}\right]$$

Hence,

$$k_r^{th} = \left[\frac{k_s}{\alpha} - \frac{g_r g_p k_{deact}}{k_p k_{act}}\right] - \frac{f(k_r^{slope})}{f'(k_r^{slope})}$$

$$\implies k_r^{th} = \left[\frac{k_s}{\alpha} - \frac{g_r g_p k_{deact}}{k_p k_{act}}\right] - 2g_r\left[\sqrt{\left(\frac{g_p k_{deact}}{k_p k_{act}}\right)^2 - \left(\frac{g_p k_{deact}}{k_p k_{act}}\right)\frac{\left(k_s^0 - k_s + \frac{\alpha g_r g_p k_{deact}}{k_p k_{act}}\right)}{\alpha g_r}} - \left(\frac{g_p k_{deact}}{k_p k_{act}}\right)\right]$$

$$\implies k_r^{th} = \frac{k_s}{\alpha} + \frac{g_r g_p k_{deact}}{k_p k_{act}} - 2g_r\sqrt{\frac{(k_s - k_s^0)}{\alpha g_r}\left(\frac{g_p k_{deact}}{k_p k_{act}}\right)} \tag{S 5.6}$$

It is to be noted that when, $k_{act} \to \infty$ then $k_r^{th} \to \frac{k_s}{\alpha}$

We can also derive explicit expression of $k_r^{th}$ for nonzero $k_{act}$.

- **Consistency check of $k_r^{th}$ for $k_{act} = 0s^{-1}$**

From S 4.6 we have the mean mRNA curve as,

$$< r > = f(k_r) = -\frac{g_s g_r + g\left(k_s^0 - \alpha k_r\right)}{2\alpha g g_r} + \sqrt{\left\{\frac{g_s g_r + g\left(k_s^0 - \alpha k_r\right)}{2\alpha g g_r}\right\}^2 + \frac{g_s k_r}{\alpha g g_r}}$$

$$\implies f'(k_r) = \frac{1}{2g_r} + \frac{1}{2g_r}\frac{\left[\frac{g_s g_r - \left(k_s^0 - \alpha k_r\right)}{2\alpha g g_r}\right]}{\sqrt{\left\{\frac{g_s g_r + (k_s^0 - \alpha k_r)}{2\alpha g g_r}\right\}^2 + \frac{g_s k_r}{\alpha g g_r}}}$$

$$\implies f''(k_r) = \left(\frac{1}{2g_r}\right)^2 \frac{\left(\frac{g_s k_s^0}{\alpha^2 g g_r}\right)}{\left[\left\{\frac{g_s g_r + (k_s^0 - \alpha k_r)}{2\alpha g g_r}\right\}^2 + \frac{g_s k_r}{\alpha g g_r}\right]^{3/2}}$$

$$\implies f'''(k_r) = -\frac{3}{2}\frac{k_s^0 g_s}{(2\alpha g_r)^2}\frac{\frac{1}{2\alpha g g_r^2}\left\{g_s g_r - g(k_s^0 - \alpha k_r)\right\}}{\left[\left\{\frac{g_s g_r + (k_s^0 - \alpha k_r)}{2\alpha g g_r}\right\}^2 + \frac{g_s k_r}{\alpha g g_r}\right]^{5/2}}$$

so, $k_r^{slope}$ is given by,

$$f'''(k_r^{slope}) = 0$$
$$\implies k_r^{slope} = \left(\frac{k_s^0}{\alpha} - \frac{g_s g_r}{\alpha g}\right)$$

and hence the threshold is given by,

$$k_r^{th} = k_r^{slope} - \frac{f(k_r^{slope})}{f'(k_r^{slope})} = \Big(\frac{k_s^0}{\alpha} - \frac{g_s g_r}{\alpha g}\Big) - \frac{f(k_r^{slope})}{f'(k_r^{slope})}$$

where,

$$f(k_r^{slope}) = -\frac{g_s}{\alpha g} + \sqrt{\frac{g_s k_s^0}{\alpha^2 g g_r}}$$
$$\& \quad f'(k_r^{slope}) = \frac{1}{2g_r}$$

Now when $k_+ >> (k_- + \gamma)$ ,i.e, $g \to \infty$ we have,

$$\left.\begin{matrix} f(k_r^{slope}) \to 0 & \& & f'(k_r^{slope}) \to \frac{1}{2g_r} \end{matrix}\right\} \implies \frac{f(k_r^{slope})}{f'(k_r^{slope})} \to 0$$

Therefore,

$$\lim_{g\to\infty} k_r^{th} = \lim_{g\to\infty}\left[\Big(\frac{k_s^0}{\alpha} - \frac{g_s g_r}{\alpha g}\Big) - \frac{f(k_r^{slope})}{f'(k_r^{slope})}\right] = \frac{k_s^0}{\alpha}$$

Hence, we get back the same threshold value $\frac{k_s^0}{\alpha}$ as in the equation S 5.3. This compliments our earlier discussion.

##### 5.1.2 General $k_s$ threshold

From S 4.6 we observe that, $\langle r\rangle$ is independent of $k_s$ when $k_{act} = 0$. So the $k_s$ threshold is only sensible when $k_{act} \neq 0$. We follow the same technique as described in section (5.1.1) and consider $\langle r\rangle = f(k_s)$ where $f(k_s)$ is right hand side of S 4.7. Since we are only interested only in the case of $g \to \infty$, from S 5.5,

$$f'(k_s) = -\frac{1}{2\alpha g_r} + \frac{\frac{1}{2\alpha g_r}\left[\frac{(k_s-\alpha k_r)}{\alpha g_r} + \frac{g_p k_{deact}}{k_p k_{act}}\right]}{2\sqrt{\frac{1}{4}\left[\frac{(k_s-\alpha k_r)}{\alpha g_r} + \frac{g_p k_{deact}}{k_p k_{act}}\right]^2 - \frac{g_p k_{deact}(k_s^0-\alpha k_r)}{\alpha g_r k_p k_{act}}}}$$

$$\implies f''(k_s) = \frac{1}{(2\alpha g_r)^2}\cdot\frac{\frac{g_p k_{deact}(\alpha k_r - k_s^0)}{\alpha g_r k_p k_{act}}}{\left[\frac{1}{4}\left\{\frac{(k_s-\alpha k_r)}{\alpha g_r} + \frac{g_p k_{deact}}{k_p k_{act}}\right\}^2 - \frac{g_p k_{deact}(k_s^0-\alpha k_r)}{\alpha g_r k_p k_{act}}\right]^{3/2}}$$

$$\implies f'''(k_s) = \frac{3 g_p k_{deact}(k_s^0 - \alpha k_r)}{8 k_p k_{act}(\alpha g_r)^4}\cdot\frac{\left[\frac{(k_s-\alpha k_r)}{\alpha g_r} + \frac{g_p k_{deact}}{k_p k_{act}}\right]}{\left[\frac{1}{4}\left\{\frac{(k_s-\alpha k_r)}{\alpha g_r} + \frac{g_p k_{deact}}{k_p k_{act}}\right\}^2 - \frac{g_p k_{deact}(k_s^0-\alpha k_r)}{\alpha g_r k_p k_{act}}\right]^{5/2}}$$

So $k_s^{slope}$ is given by,

$$f'''(k_s^{slope}) = 0$$
$$\implies \left[\frac{(k_s^{slope} - \alpha k_r)}{\alpha g_r} + \frac{g_p k_{deact}}{k_p k_{act}}\right] = 0 \qquad \left[(k_s^0 - \alpha k_r) \neq 0 \quad \text{otherwise} \quad \langle r\rangle = f(k_s) = 0\right]$$
$$\implies k_s^{slope} = \alpha g_r\Big(\frac{k_r}{g_r} - \frac{g_p k_{deact}}{k_p k_{act}}\Big)$$

$k_s = k_s^{slope}$ is the $k_s$ value where the change in slope of $\langle r\rangle = f(k_s)$ is maximum.

Therefore, $k_s$ threshold is given by,

$$k_s^{th} = k_s^{slope} - \frac{f(k_s^{slope})}{f'(k_s^{slope})}$$
$$\implies k_s^{th} = \alpha g_r\Big(\frac{k_r}{g_r} - \frac{g_p k_{deact}}{k_p k_{act}}\Big) + 2\alpha g_r\sqrt{\frac{g_p k_{deact}(\alpha k_r - k_s^0)}{\alpha g_r k_p k_{act}}} \qquad \text{(S 5.7)}$$

### 6 Formation of second order moment equations by approximating third order moments

In order to investigate our miRNA-mediated negative feedback model at the noise level we tried to find the second order moments of our variables of interest (i.e, $r(t), p(t), s(t)$ & $c(t)$). Equations S 2.1 - S 2.29 contains the following third order moments,

| Third order moments | | | |
|---|---|---|---|
| $< r^2 s >^{ON}$ | $< r^2 p >^{OFF}$ | $< r^2 s >^{OFF}$ | $< p^3 >^{OFF}$ |
| $< ps^2 >^{OFF}$ | $< pc^2 >^{OFF}$ | $< rps >^{OFF}$ | $< rs^2 >^{ON}$ |
| $< rsc >^{ON}$ | $< rsc >^{OFF}$ | $< rs^2 >^{OFF}$ | $< rpc >^{OFF}$ |
| $< rp^2 >^{OFF}$ | $< rps >^{ON}$ | $< p^2 s >^{OFF}$ | $< p^2 c >^{OFF}$ |
| $< psc >^{OFF}$ | | | |

Now we approximate these third order moments with second order moments as follows,

**I.** $$\left\langle (r-\langle r\rangle)^2(s-\langle s\rangle)\right\rangle^{ON} = 0$$
$$\begin{aligned}\implies \langle r^2 s\rangle^{ON} &= 2\langle rs\rangle^{ON}\left(\langle r\rangle^{ON}+\langle r\rangle^{OFF}\right)-\left(\langle r\rangle^{ON}+\langle r\rangle^{OFF}\right)^2\langle s\rangle^{ON}\\ &-2\langle r\rangle^{ON}\left(\langle r\rangle^{ON}+\langle r\rangle^{OFF}\right)\left(\langle s\rangle^{ON}+\langle s\rangle^{OFF}\right)+\langle r^2\rangle^{ON}\left(\langle s\rangle^{ON}+\langle s\rangle^{OFF}\right)\\ &+\left(\langle r\rangle^{ON}+\langle r\rangle^{OFF}\right)^2\left(\langle s\rangle^{ON}+\langle s\rangle^{ON}\right)P^{ON}\end{aligned}$$

**II.** $$\left\langle (r-\langle r\rangle)^2(p-\langle p\rangle)\right\rangle^{OFF} = 0$$
$$\begin{aligned}\implies \langle r^2 p\rangle^{OFF} &= 2\langle rp\rangle^{OFF}\left(\langle r\rangle^{ON}+\langle r\rangle^{OFF}\right)+\left(\langle r\rangle^{ON}+\langle r\rangle^{OFF}\right)^2\langle p\rangle^{ON}\\ &-2\langle r\rangle^{OFF}\left(\langle r\rangle^{ON}+\langle r\rangle^{OFF}\right)\left(\langle p\rangle^{ON}+\langle p\rangle^{OFF}\right)\\ &+\langle r^2\rangle^{OFF}\left(\langle p\rangle^{ON}+\langle p\rangle^{OFF}\right)-\left(\langle r\rangle^{ON}+\langle r\rangle^{OFF}\right)^2\left(\langle p\rangle^{ON}+\langle p\rangle^{OFF}\right)P^{ON}\end{aligned}$$

**III.** $$\left\langle (r-\langle r\rangle)^2(s-\langle s\rangle)\right\rangle^{OFF} = 0$$
$$\begin{aligned}\implies \langle r^2 s\rangle^{OFF} &= 2\langle rs\rangle^{OFF}\left(\langle r\rangle^{ON}+\langle r\rangle^{OFF}\right)+\left(\langle r\rangle^{ON}+\langle r\rangle^{OFF}\right)^2\langle s\rangle^{ON}\\ &-2\langle r\rangle^{OFF}\left(\langle r\rangle^{ON}+\langle r\rangle^{OFF}\right)\left(\langle s\rangle^{ON}+\langle s\rangle^{OFF}\right)\\ &+\langle r^2\rangle^{OFF}\left(\langle s\rangle^{ON}+\langle s\rangle^{OFF}\right)-\left(\langle r\rangle^{ON}+\langle r\rangle^{OFF}\right)^2\left(\langle s\rangle^{ON}+\langle s\rangle^{OFF}\right)P^{ON}\end{aligned}$$

**IV.** $$\left\langle (p-\langle p\rangle)^3\right\rangle^{OFF} = 0$$
$$\begin{aligned}\implies \langle p^3\rangle^{OFF} &= \left(\langle p\rangle^{ON}+\langle p\rangle^{OFF}\right)\Big[\left(\langle p\rangle^{ON}+\langle p\rangle^{OFF}\right)^2\left(1-P^{ON}\right)+3\langle p^2\rangle^{OFF}\\ &-3\langle p\rangle^{OFF}\left(\langle p\rangle^{ON}+\langle p\rangle^{OFF}\right)\Big]\end{aligned}$$

**V.** $$\left\langle (p-\langle p\rangle)(s-\langle s\rangle)^2\right\rangle^{OFF} = 0$$
$$\begin{aligned}\implies \langle ps^2\rangle^{OFF} &= 2\langle ps\rangle^{OFF}\left(\langle s\rangle^{ON}+\langle s\rangle^{OFF}\right)+\langle p\rangle^{ON}\left(\langle s\rangle^{ON}+\langle s\rangle^{OFF}\right)^2\\ &-2\langle s\rangle^{OFF}\left(\langle p\rangle^{ON}+\langle p\rangle^{OFF}\right)\left(\langle s\rangle^{ON}+\langle s\rangle^{OFF}\right)\\ &+\left(\langle p\rangle^{ON}+\langle p\rangle^{OFF}\right)\langle s^2\rangle^{OFF}-\left(\langle p\rangle^{ON}+\langle p\rangle^{OFF}\right)\left(\langle s\rangle^{ON}+\langle s\rangle^{OFF}\right)^2P^{ON}\end{aligned}$$

**VI.** $$\left\langle (p-\langle p\rangle)(c-\langle c\rangle)^2\right\rangle^{OFF} = 0$$
$$\begin{aligned}\implies \langle pc^2\rangle^{OFF} &= 2\langle pc\rangle^{OFF}\left(\langle c\rangle^{ON}+\langle c\rangle^{OFF}\right)+\langle p\rangle^{ON}\left(\langle c\rangle^{ON}+\langle c\rangle^{OFF}\right)^2\\ &-2\left(\langle p\rangle^{ON}+\langle p\rangle^{OFF}\right)\left(\langle c\rangle^{ON}+\langle c\rangle^{OFF}\right)\langle c\rangle^{OFF}\\ &+\left(\langle p\rangle^{ON}+\langle p\rangle^{OFF}\right)\langle c^2\rangle^{OFF}-\left(\langle p\rangle^{ON}+\langle p\rangle^{OFF}\right)\left(\langle c\rangle^{ON}+\langle c\rangle^{OFF}\right)^2P^{ON}\end{aligned}$$

**VII.** $$\left\langle (r-\langle r\rangle)(s-\langle s\rangle)(c-\langle c\rangle)\right\rangle^{ON} = 0$$
$$\begin{aligned}\implies \langle rsc\rangle^{ON} &= \langle rs\rangle^{ON}\left(\langle c\rangle^{ON}+\langle c\rangle^{OFF}\right)+\langle rc\rangle^{ON}\left(\langle s\rangle^{ON}+\langle s\rangle^{OFF}\right)\\ &-\langle r\rangle^{ON}\left(\langle s\rangle^{ON}+\langle s\rangle^{OFF}\right)\left(\langle c\rangle^{ON}+\langle c\rangle^{OFF}\right)+\left(\langle r\rangle^{ON}+\langle r\rangle^{OFF}\right)\langle sc\rangle^{ON}\\ &-\left(\langle r\rangle^{ON}+\langle r\rangle^{OFF}\right)\langle s\rangle^{ON}\left(\langle c\rangle^{ON}+\langle c\rangle^{OFF}\right)\\ &-\left(\langle r\rangle^{ON}+\langle r\rangle^{OFF}\right)\left(\langle s\rangle^{ON}+\langle s\rangle^{OFF}\right)\langle c\rangle^{ON}\\ &+P^{ON}\left(\langle r\rangle^{ON}+\langle r\rangle^{OFF}\right)\left(\langle s\rangle^{ON}+\langle s\rangle^{OFF}\right)\left(\langle c\rangle^{ON}+\langle c\rangle^{OFF}\right)\end{aligned}$$

**VIII.** $\langle (r-\langle r\rangle)(s-\langle s\rangle)(c-\langle c\rangle)\rangle^{OFF} = 0$

$$\begin{aligned}
\implies \langle rsc\rangle^{OFF} &= \langle rs\rangle^{OFF}\left(\langle c\rangle^{ON}+\langle c\rangle^{OFF}\right)+\langle rc\rangle^{OFF}\left(\langle s\rangle^{ON}+\langle s\rangle^{OFF}\right)\\
&-\langle r\rangle^{OFF}\left(\langle s\rangle^{ON}+\langle s\rangle^{OFF}\right)\left(\langle c\rangle^{ON}+\langle c\rangle^{OFF}\right)\\
&+\left(\langle r\rangle^{ON}+\langle r\rangle^{OFF}\right)\langle sc\rangle^{OFF}\\
&-\left(\langle r\rangle^{ON}+\langle r\rangle^{OFF}\right)\langle s\rangle^{OFF}\left(\langle c\rangle^{ON}+\langle c\rangle^{OFF}\right)\\
&+\left(\langle r\rangle^{ON}+\langle r\rangle^{OFF}\right)\left(\langle s\rangle^{ON}+\langle s\rangle^{OFF}\right)\langle c\rangle^{ON}\\
&-P^{ON}\left(\langle r\rangle^{ON}+\langle r\rangle^{OFF}\right)\left(\langle s\rangle^{ON}+\langle s\rangle^{OFF}\right)\left(\langle c\rangle^{ON}+\langle c\rangle^{OFF}\right)
\end{aligned}$$

**IX.** $\langle (r-\langle r\rangle)(p-\langle p\rangle)(s-\langle s\rangle)\rangle^{OFF} = 0$

$$\begin{aligned}
\implies \langle rps\rangle^{OFF} &= \langle rp\rangle^{OFF}\left(\langle s\rangle^{ON}+\langle s\rangle^{OFF}\right)+\langle rs\rangle^{OFF}\left(\langle p\rangle^{ON}+\langle p\rangle^{OFF}\right)\\
&-\langle r\rangle^{OFF}\left(\langle p\rangle^{ON}+\langle p\rangle^{OFF}\right)\left(\langle s\rangle^{ON}+\langle s\rangle^{OFF}\right)\\
&+\left(\langle r\rangle^{ON}+\langle r\rangle^{OFF}\right)\langle ps\rangle^{OFF}\\
&-\left(\langle r\rangle^{ON}+\langle r\rangle^{OFF}\right)\langle p\rangle^{OFF}\left(\langle s\rangle^{ON}+\langle s\rangle^{OFF}\right)\\
&+\left(\langle r\rangle^{ON}+\langle r\rangle^{OFF}\right)\left(\langle p\rangle^{ON}+\langle p\rangle^{OFF}\right)\langle s\rangle^{ON}\\
&-P^{ON}\left(\langle r\rangle^{ON}+\langle r\rangle^{OFF}\right)\left(\langle p\rangle^{ON}+\langle p\rangle^{OFF}\right)\left(\langle s\rangle^{ON}+\langle s\rangle^{OFF}\right)
\end{aligned}$$

**X.** $\left\langle (r-\langle r\rangle)(s-\langle s\rangle)^2\right\rangle^{ON} = 0$

$$\begin{aligned}
\implies \langle rs^2\rangle^{ON} &= 2\langle rs\rangle^{ON}\left(\langle s\rangle^{ON}+\langle s\rangle^{OFF}\right)-\langle r\rangle^{ON}\left(\langle s\rangle^{ON}+\langle s\rangle^{OFF}\right)^2\\
&+\left(\langle r\rangle^{ON}+\langle r\rangle^{OFF}\right)\langle s^2\rangle^{ON}\\
&-2\left(\langle r\rangle^{ON}+\langle r\rangle^{OFF}\right)\left(\langle s\rangle^{ON}+\langle s\rangle^{OFF}\right)\langle s\rangle^{ON}\\
&+\left(\langle r\rangle^{ON}+\langle r\rangle^{OFF}\right)\left(\langle s\rangle^{ON}+\langle s\rangle^{OFF}\right)^2P^{ON}
\end{aligned}$$

**XI.** $\left\langle (r-\langle r\rangle)(s-\langle s\rangle)^2\right\rangle^{OFF} = 0$

$$\begin{aligned}
\implies \langle rs^2\rangle^{OFF} &= 2\langle rs\rangle^{OFF}\left(\langle s\rangle^{ON}+\langle s\rangle^{OFF}\right)+\langle r\rangle^{ON}\left(\langle s\rangle^{ON}+\langle s\rangle^{OFF}\right)^2\\
&+\left(\langle r\rangle^{ON}+\langle r\rangle^{OFF}\right)\langle s^2\rangle^{OFF}\\
&-2\left(\langle r\rangle^{ON}+\langle r\rangle^{OFF}\right)\left(\langle s\rangle^{ON}+\langle s\rangle^{OFF}\right)\langle s\rangle^{OFF}\\
&-\left(\langle r\rangle^{ON}+\langle r\rangle^{OFF}\right)\left(\langle s\rangle^{ON}+\langle s\rangle^{OFF}\right)^2P^{ON}
\end{aligned}$$

**XII.** $\langle (r-\langle r\rangle)(p-\langle p\rangle)(c-\langle c\rangle)\rangle^{OFF} = 0$

$$\begin{aligned}
\implies \langle rpc\rangle^{OFF} &= \langle rp\rangle^{OFF}\left(\langle c\rangle^{ON}+\langle c\rangle^{OFF}\right)+\langle rc\rangle^{OFF}\left(\langle p\rangle^{ON}+\langle p\rangle^{OFF}\right)\\
&-\langle r\rangle^{OFF}\left(\langle p\rangle^{ON}+\langle p\rangle^{OFF}\right)\left(\langle c\rangle^{ON}+\langle c\rangle^{OFF}\right)\\
&+\left(\langle r\rangle^{ON}+\langle r\rangle^{OFF}\right)\langle pc\rangle^{OFF}\\
&+\left(\langle r\rangle^{ON}+\langle r\rangle^{OFF}\right)\langle p\rangle^{ON}\left(\langle c\rangle^{ON}+\langle c\rangle^{OFF}\right)\\
&-\left(\langle r\rangle^{ON}+\langle r\rangle^{OFF}\right)\left(\langle p\rangle^{ON}+\langle p\rangle^{OFF}\right)\langle c\rangle^{OFF}\\
&-P^{ON}\left(\langle r\rangle^{ON}+\langle r\rangle^{OFF}\right)\left(\langle p\rangle^{ON}+\langle p\rangle^{OFF}\right)\left(\langle c\rangle^{ON}+\langle c\rangle^{OFF}\right)
\end{aligned}$$

**XIII.** $\left\langle (r-\langle r\rangle)(p-\langle p\rangle)^2\right\rangle^{OFF} = 0$

$$\begin{aligned}
\implies \langle rp^2\rangle^{OFF} &= 2\langle rp\rangle^{OFF}\left(\langle p\rangle^{ON}+\langle p\rangle^{OFF}\right)+\langle r\rangle^{ON}\left(\langle p\rangle^{ON}+\langle p\rangle^{OFF}\right)^2\\
&+\left(\langle r\rangle^{ON}+\langle r\rangle^{OFF}\right)\langle p^2\rangle^{OFF}\\
&-2\left(\langle r\rangle^{ON}+\langle r\rangle^{OFF}\right)\left(\langle p\rangle^{ON}+\langle p\rangle^{OFF}\right)\langle p\rangle^{OFF}\\
&-\left(\langle r\rangle^{ON}+\langle r\rangle^{OFF}\right)\left(\langle p\rangle^{ON}+\langle p\rangle^{OFF}\right)^2P^{ON}
\end{aligned}$$

**XIV.** $\langle (r-\langle r\rangle)(p-\langle p\rangle)(s-\langle s\rangle)\rangle^{ON} = 0$

$$\begin{aligned}
\implies \langle rps\rangle^{ON} &= \langle rp\rangle^{ON}\left(\langle s\rangle^{ON}+\langle s\rangle^{OFF}\right)+\langle rs\rangle^{ON}\left(\langle p\rangle^{ON}+\langle p\rangle^{OFF}\right)\\
&-\langle r\rangle^{ON}\left(\langle p\rangle^{ON}+\langle p\rangle^{OFF}\right)\left(\langle s\rangle^{ON}+\langle s\rangle^{OFF}\right)\\
&+\left(\langle r\rangle^{ON}+\langle r\rangle^{OFF}\right)\langle ps\rangle^{ON}\\
&-\left(\langle r\rangle^{ON}+\langle r\rangle^{OFF}\right)\langle p\rangle^{ON}\left(\langle s\rangle^{ON}+\langle s\rangle^{OFF}\right)\\
&-\left(\langle r\rangle^{ON}+\langle r\rangle^{OFF}\right)\left(\langle p\rangle^{ON}+\langle p\rangle^{OFF}\right)\langle s\rangle^{ON}\\
&+P^{ON}\left(\langle r\rangle^{ON}+\langle r\rangle^{OFF}\right)\left(\langle p\rangle^{ON}+\langle p\rangle^{OFF}\right)\left(\langle s\rangle^{ON}+\langle s\rangle^{OFF}\right)
\end{aligned}$$

$$\textbf{XV.} \quad \left\langle (p-\langle p\rangle)^2(s-\langle s\rangle)\right\rangle^{OFF} = 0$$

$$\implies \langle p^2 s\rangle^{OFF} = 2\langle ps\rangle^{OFF}\left(\langle p\rangle^{ON}+\langle p\rangle^{OFF}\right)+\left(\langle p\rangle^{ON}+\langle p\rangle^{OFF}\right)^2\langle s\rangle^{ON}$$
$$-2\langle p\rangle^{OFF}\left(\langle p\rangle^{ON}+\langle p\rangle^{OFF}\right)\left(\langle s\rangle^{ON}+\langle s\rangle^{OFF}\right)$$
$$+\langle p^2\rangle^{OFF}\left(\langle s\rangle^{ON}+\langle s\rangle^{OFF}\right)$$
$$-\left(\langle p\rangle^{ON}+\langle p\rangle^{OFF}\right)^2\left(\langle s\rangle^{ON}+\langle s\rangle^{OFF}\right)P^{ON}$$

$$\textbf{XVI.} \quad \left\langle (p-\langle p\rangle)^2(c-\langle c\rangle)\right\rangle^{OFF} = 0$$

$$\implies \langle p^2 c\rangle^{OFF} = 2\langle pc\rangle^{OFF}\left(\langle p\rangle^{ON}+\langle p\rangle^{OFF}\right)+\left(\langle p\rangle^{ON}+\langle p\rangle^{OFF}\right)^2\langle c\rangle^{ON}$$
$$-2\langle p\rangle^{OFF}\left(\langle p\rangle^{ON}+\langle p\rangle^{OFF}\right)\left(\langle c\rangle^{ON}+\langle c\rangle^{OFF}\right)$$
$$+\langle p^2\rangle^{OFF}\left(\langle c\rangle^{ON}+\langle c\rangle^{OFF}\right)$$
$$-\left(\langle p\rangle^{ON}+\langle p\rangle^{OFF}\right)^2\left(\langle c\rangle^{ON}+\langle c\rangle^{OFF}\right)P^{ON}$$

$$\textbf{XVII.} \quad \langle (p-\langle p\rangle)(s-\langle s\rangle)(c-\langle c\rangle)\rangle^{OFF} = 0$$

$$\implies \langle psc\rangle^{OFF} = \langle ps\rangle^{OFF}\left(\langle c\rangle^{ON}+\langle c\rangle^{OFF}\right)+\langle pc\rangle^{OFF}\left(\langle s\rangle^{ON}+\langle s\rangle^{OFF}\right)$$
$$-\langle p\rangle^{OFF}\left(\langle s\rangle^{ON}+\langle s\rangle^{OFF}\right)\left(\langle c\rangle^{ON}+\langle c\rangle^{OFF}\right)$$
$$+\left(\langle p\rangle^{ON}+\langle p\rangle^{OFF}\right)\langle sc\rangle^{OFF}$$
$$-\left(\langle p\rangle^{ON}+\langle p\rangle^{OFF}\right)\langle s\rangle^{OFF}\left(\langle c\rangle^{ON}+\langle c\rangle^{OFF}\right)$$
$$+\left(\langle p\rangle^{ON}+\langle p\rangle^{OFF}\right)\left(\langle s\rangle^{ON}+\langle s\rangle^{OFF}\right)\langle c\rangle^{ON}$$
$$-P^{ON}\left(\langle p\rangle^{ON}+\langle p\rangle^{OFF}\right)\left(\langle s\rangle^{ON}+\langle s\rangle^{OFF}\right)\left(\langle c\rangle^{ON}+\langle c\rangle^{OFF}\right)$$

Using these approximations we get from equations S 2.10 - S 2.29,

$$\textbf{i)} \quad \frac{\partial\langle r^2\rangle^{ON}}{\partial t} = k_r\left[2\langle r\rangle^{ON}+P^{ON}\right]+g_r\left[\langle r\rangle^{ON}-2\langle r^2\rangle^{ON}\right]-k_{deact}\langle r^2\rangle^{ON}$$
$$+k_{act}\Bigg[2\langle rp\rangle^{OFF}\left(\langle r\rangle^{ON}+\langle r\rangle^{OFF}\right)+\left(\langle r\rangle^{ON}+\langle r\rangle^{OFF}\right)^2\langle p\rangle^{ON}$$
$$-2\langle r\rangle^{OFF}\left(\langle r\rangle^{ON}+\langle r\rangle^{OFF}\right)\left(\langle p\rangle^{ON}+\langle p\rangle^{OFF}\right)+\langle r^2\rangle^{OFF}\left(\langle p\rangle^{ON}+\langle p\rangle^{OFF}\right)$$
$$-\left(\langle r\rangle^{ON}+\langle r\rangle^{OFF}\right)^2\left(\langle p\rangle^{ON}+\langle p\rangle^{OFF}\right)P^{ON}\Bigg]+k_-\left[\langle c\rangle^{ON}+2\langle rc\rangle^{ON}\right]$$
$$-k_+\Bigg[\langle rs\rangle^{ON}-2\Bigg\{2\langle rs\rangle^{ON}\left(\langle r\rangle^{ON}+\langle r\rangle^{OFF}\right)-\left(\langle r\rangle^{ON}+\langle r\rangle^{OFF}\right)^2\langle s\rangle^{ON}$$
$$-2\langle r\rangle^{ON}\left(\langle r\rangle^{ON}+\langle r\rangle^{OFF}\right)\left(\langle s\rangle^{ON}+\langle s\rangle^{OFF}\right)$$
$$+\langle r^2\rangle^{ON}\left(\langle s\rangle^{ON}+\langle s\rangle^{OFF}\right)+\left(\langle r\rangle^{ON}+\langle r\rangle^{OFF}\right)^2\left(\langle s\rangle^{ON}+\langle s\rangle^{OFF}\right)P^{ON}\Bigg\}\Bigg] \qquad \text{(S 6.1)}$$

$$\textbf{ii)} \quad \frac{\partial\langle r^2\rangle^{OFF}}{\partial t} = k_r\left[2\langle r\rangle^{OFF}+P^{OFF}\right]+g_r\left[\langle r\rangle^{OFF}-2\langle r^2\rangle^{OFF}\right]+k_{deact}\langle r^2\rangle^{ON}$$
$$-k_{act}\Bigg[2\langle rp\rangle^{OFF}\left(\langle r\rangle^{ON}+\langle r\rangle^{OFF}\right)+\left(\langle r\rangle^{ON}+\langle r\rangle^{OFF}\right)^2\langle p\rangle^{ON}$$
$$-2\langle r\rangle^{OFF}\left(\langle r\rangle^{ON}+\langle r\rangle^{OFF}\right)\left(\langle p\rangle^{ON}+\langle p\rangle^{OFF}\right)+\langle r^2\rangle^{OFF}\left(\langle p\rangle^{ON}+\langle p\rangle^{OFF}\right)$$
$$-\left(\langle r\rangle^{ON}+\langle r\rangle^{OFF}\right)^2\left(\langle p\rangle^{ON}+\langle p\rangle^{OFF}\right)P^{ON}\Bigg]+k_-\left[2\langle rc\rangle^{OFF}+\langle c\rangle^{OFF}\right]$$
$$+k_+\Bigg[\langle rs\rangle^{OFF}-2\Bigg\{2\langle rs\rangle^{OFF}\left(\langle r\rangle^{ON}+\langle r\rangle^{OFF}\right)+\left(\langle r\rangle^{ON}+\langle r\rangle^{OFF}\right)^2\langle s\rangle^{ON}$$
$$-2\langle r\rangle^{OFF}\left(\langle r\rangle^{ON}+\langle r\rangle^{OFF}\right)\left(\langle s\rangle^{ON}+\langle s\rangle^{OFF}\right)+\langle r^2\rangle^{OFF}\left(\langle s\rangle^{ON}+\langle s\rangle^{OFF}\right)$$
$$-\left(\langle r\rangle^{ON}+\langle r\rangle^{OFF}\right)^2\left(\langle s\rangle^{ON}+\langle s\rangle^{OFF}\right)P^{ON}\Bigg\}\Bigg] \qquad \text{(S 6.2)}$$

**iii)** $$\frac{\partial \langle p^2\rangle^{ON}}{\partial t} = k_p\left[\langle r\rangle^{ON} + 2\langle rp\rangle^{ON}\right] + g_p\left[\langle p\rangle^{ON} - 2\langle p^2\rangle^{ON}\right] - k_{deact}\langle p^2\rangle^{ON}$$
$$+ k_{act}\Bigg[\left(\langle p\rangle^{ON} + \langle p\rangle^{OFF}\right)\Big\{\left(\langle p\rangle^{ON} + \langle p\rangle^{OFF}\right)^2\left(1 - P^{ON}\right) + 3\langle p^2\rangle^{OFF}$$
$$- 3\langle p\rangle^{OFF}\left(\langle p\rangle^{ON} + \langle p\rangle^{OFF}\right)\Big\} - 2\langle p^2\rangle^{OFF} + \langle p\rangle^{OFF}\Bigg] \quad \text{(S 6.3)}$$

**iv)** $$\frac{\partial \langle p^2\rangle^{OFF}}{\partial t} = k_p\left[\langle r\rangle^{OFF} + 2\langle rp\rangle^{OFF}\right] + g_p\left[\langle p\rangle^{OFF} - 2\langle p^2\rangle^{OFF}\right] + k_{deact}\left[\langle p^2\rangle^{ON} + 2\langle p\rangle^{ON} + P^{ON}\right]$$
$$- k_{act}\Bigg[\left(\langle p\rangle^{ON} + \langle p\rangle^{OFF}\right)\Big\{\left(\langle p\rangle^{ON} + \langle p\rangle^{OFF}\right)^2\left(1 - P^{ON}\right)$$
$$+ 3\langle p^2\rangle^{OFF} - 3\langle p\rangle^{OFF}\left(\langle p\rangle^{ON} + \langle p\rangle^{OFF}\right)\Big\}\Bigg] \quad \text{(S 6.4)}$$

**v)** $$\frac{\partial \langle s^2\rangle^{ON}}{\partial t} = k_{act}\Bigg[2\langle ps\rangle^{OFF}\left(\langle s\rangle^{ON} + \langle s\rangle^{OFF}\right) + \langle p\rangle^{ON}\left(\langle s\rangle^{ON} + \langle s\rangle^{OFF}\right)^2$$
$$- 2\langle s\rangle^{OFF}\left(\langle p\rangle^{ON} + \langle p\rangle^{OFF}\right)\left(\langle s\rangle^{ON} + \langle s\rangle^{OFF}\right) + \left(\langle p\rangle^{ON} + \langle p\rangle^{OFF}\right)\langle s^2\rangle^{OFF}$$
$$- \left(\langle p\rangle^{ON} + \langle p\rangle^{OFF}\right)\left(\langle s\rangle^{ON} + \langle s\rangle^{OFF}\right)^2 P^{ON}\Bigg] - k_{deact}\langle s^2\rangle^{ON}$$
$$+ k_s\left[2\langle s\rangle^{ON} + P^{ON}\right] + g_s\left[\langle s\rangle^{ON} - 2\langle s^2\rangle^{ON}\right]$$
$$+ k_+\Bigg[\langle rs\rangle^{ON} - 2\Big\{2\langle rs\rangle^{ON}\left(\langle s\rangle^{ON} + \langle s\rangle^{OFF}\right) - \langle r\rangle^{ON}\left(\langle s\rangle^{ON} + \langle s\rangle^{OFF}\right)^2$$
$$+ \left(\langle r\rangle^{ON} + \langle r\rangle^{OFF}\right)\langle s^2\rangle^{ON} - 2\left(\langle r\rangle^{ON} + \langle r\rangle^{OFF}\right)\left(\langle s\rangle^{ON} + \langle s\rangle^{OFF}\right)\langle s\rangle^{ON}$$
$$+ \left(\langle r\rangle^{ON} + \langle r\rangle^{OFF}\right)\left(\langle s\rangle^{ON} + \langle s\rangle^{OFF}\right)^2 P^{ON}\Big\}\Bigg]$$
$$+ \left[k_- + (1-\alpha)\gamma\right]\left[\langle c\rangle^{ON} + 2\langle sc\rangle^{ON}\right] \quad \text{(S 6.5)}$$

**vi)** $$\frac{\partial \langle s^2\rangle^{OFF}}{\partial t} = k_s^0\left[2\langle s\rangle^{OFF} + P^{OFF}\right] + g_s\left[\langle s\rangle^{OFF} - 2\langle s^2\rangle^{OFF}\right] + k_{deact}\langle s^2\rangle^{ON}$$
$$- k_{act}\Bigg[2\langle ps\rangle^{OFF}\left(\langle s\rangle^{ON} + \langle s\rangle^{OFF}\right) + \langle p\rangle^{ON}\left(\langle s\rangle^{ON} + \langle s\rangle^{OFF}\right)^2$$
$$- 2\langle s\rangle^{OFF}\left(\langle p\rangle^{ON} + \langle p\rangle^{OFF}\right)\left(\langle s\rangle^{ON} + \langle s\rangle^{OFF}\right) + \left(\langle p\rangle^{ON} + \langle p\rangle^{OFF}\right)\langle s^2\rangle^{OFF}$$
$$- \left(\langle p\rangle^{ON} + \langle p\rangle^{OFF}\right)\left(\langle s\rangle^{ON} + \langle s\rangle^{OFF}\right)^2 P^{ON}\Bigg] + \left[k_- + (1-\alpha)\gamma\right]\left[\langle c\rangle^{OFF} + 2\langle sc\rangle^{OFF}\right]$$
$$+ k_+\Bigg[\langle rs\rangle^{OFF} - 2\Big\{2\langle rs\rangle^{OFF}\left(\langle s\rangle^{ON} + \langle s\rangle^{OFF}\right) + \langle r\rangle^{ON}\left(\langle s\rangle^{ON} + \langle s\rangle^{OFF}\right)^2$$
$$+ \left(\langle r\rangle^{ON} + \langle r\rangle^{OFF}\right)\langle s^2\rangle^{OFF} - 2\left(\langle r\rangle^{ON} + \langle r\rangle^{OFF}\right)\left(\langle s\rangle^{ON} + \langle s\rangle^{OFF}\right)\langle s\rangle^{OFF}$$
$$- \left(\langle r\rangle^{ON} + \langle r\rangle^{OFF}\right)\left(\langle s\rangle^{ON} + \langle s\rangle^{OFF}\right)^2 P^{ON}\Big\}\Bigg] \quad \text{(S 6.6)}$$

**vii)** $$\frac{\partial \langle c^2\rangle^{ON}}{\partial t} = k_{act}\Bigg[2\langle pc\rangle^{OFF}\left(\langle c\rangle^{ON} + \langle c\rangle^{OFF}\right) + \langle p\rangle^{ON}\left(\langle c\rangle^{ON} + \langle c\rangle^{OFF}\right)^2$$
$$- 2\left(\langle p\rangle^{ON} + \langle p\rangle^{OFF}\right)\left(\langle c\rangle^{ON} + \langle c\rangle^{OFF}\right)\langle c\rangle^{OFF} + \left(\langle p\rangle^{ON} + \langle p\rangle^{OFF}\right)\langle c^2\rangle^{OFF}$$
$$- \left(\langle p\rangle^{ON} + \langle p\rangle^{OFF}\right)\left(\langle c\rangle^{ON} + \langle c\rangle^{OFF}\right)^2 P^{ON}\Bigg] - k_{deact}\langle c^2\rangle^{ON} + (k_- + \gamma)\left[\langle c\rangle^{ON} - 2\langle c^2\rangle^{ON}\right]$$
$$+ k_+\Bigg[\langle rs\rangle^{ON} + 2\Big\{\langle rs\rangle^{ON}\left(\langle c\rangle^{ON} + \langle c\rangle^{OFF}\right) + \langle rc\rangle^{ON}\left(\langle s\rangle^{ON} + \langle s\rangle^{OFF}\right)$$
$$- \langle r\rangle^{ON}\left(\langle s\rangle^{ON} + \langle s\rangle^{OFF}\right)\left(\langle c\rangle^{ON} + \langle c\rangle^{OFF}\right) + \left(\langle r\rangle^{ON} + \langle r\rangle^{OFF}\right)\langle sc\rangle^{ON}$$
$$- \left(\langle r\rangle^{ON} + \langle r\rangle^{OFF}\right)\langle s\rangle^{ON}\left(\langle c\rangle^{ON} + \langle c\rangle^{OFF}\right) - \left(\langle r\rangle^{ON} + \langle r\rangle^{OFF}\right)\left(\langle s\rangle^{ON} + \langle s\rangle^{OFF}\right)\langle c\rangle^{ON}$$
$$+ P^{ON}\left(\langle r\rangle^{ON} + \langle r\rangle^{OFF}\right)\left(\langle s\rangle^{ON} + \langle s\rangle^{OFF}\right)\left(\langle c\rangle^{ON} + \langle c\rangle^{OFF}\right)\Big\}\Bigg] \quad \text{(S 6.7)}$$

**viii)**

$$
\begin{aligned}
\frac{\partial \langle c^2\rangle^{OFF}}{\partial t} &= k_{deact}\langle c^2\rangle^{ON} + (k_- + \gamma)\left[\langle c\rangle^{OFF} - 2\langle c^2\rangle^{OFF}\right] \\
&- k_{act}\bigg[2\langle pc\rangle^{OFF}\left(\langle c\rangle^{ON} + \langle c\rangle^{OFF}\right) + \langle p\rangle^{ON}\left(\langle c\rangle^{ON} + \langle c\rangle^{OFF}\right)^2 \\
&- 2\left(\langle p\rangle^{ON} + \langle p\rangle^{OFF}\right)\left(\langle c\rangle^{ON} + \langle c\rangle^{OFF}\right)\langle c\rangle^{OFF} + \left(\langle p\rangle^{ON} + \langle p\rangle^{OFF}\right)\langle c^2\rangle^{OFF} \\
&- \left(\langle p\rangle^{ON} + \langle p\rangle^{OFF}\right)\left(\langle c\rangle^{ON} + \langle c\rangle^{OFF}\right)^2 P^{ON}\bigg] \\
&+ k_+\bigg[\langle rs\rangle^{OFF} + 2\Big\{\langle rs\rangle^{OFF}\left(\langle c\rangle^{ON} + \langle c\rangle^{OFF}\right) + \langle rc\rangle^{OFF}\left(\langle s\rangle^{ON} + \langle s\rangle^{OFF}\right) \\
&- \langle r\rangle^{OFF}\left(\langle s\rangle^{ON} + \langle s\rangle^{OFF}\right)\left(\langle c\rangle^{ON} + \langle c\rangle^{OFF}\right) + \left(\langle r\rangle^{ON} + \langle r\rangle^{OFF}\right)\langle sc\rangle^{OFF} \\
&- \left(\langle r\rangle^{ON} + \langle r\rangle^{OFF}\right)\langle s\rangle^{OFF}\left(\langle c\rangle^{ON} + \langle c\rangle^{OFF}\right) \\
&+ \left(\langle r\rangle^{ON} + \langle r\rangle^{OFF}\right)\left(\langle s\rangle^{ON} + \langle s\rangle^{OFF}\right)\langle c\rangle^{ON} \\
&- P^{ON}\left(\langle r\rangle^{ON} + \langle r\rangle^{OFF}\right)\left(\langle s\rangle^{ON} + \langle s\rangle^{OFF}\right)\left(\langle c\rangle^{ON} + \langle c\rangle^{OFF}\right)\Big\}\bigg] \qquad \text{(S 6.8)}
\end{aligned}
$$

**ix)**

$$
\begin{aligned}
\frac{\partial \langle rs\rangle^{ON}}{\partial t} &= k_r\langle s\rangle^{ON} - (g_r + g_s)\langle rs\rangle^{ON} + k_s\langle r\rangle^{ON} - k_{deact}\langle rs\rangle^{ON} \\
&+ k_{act}\bigg[\langle rp\rangle^{OFF}\left(\langle s\rangle^{ON} + \langle s\rangle^{OFF}\right) + \langle rs\rangle^{OFF}\left(\langle p\rangle^{ON} + \langle p\rangle^{OFF}\right) \\
&- \langle r\rangle^{OFF}\left(\langle p\rangle^{ON} + \langle p\rangle^{OFF}\right)\left(\langle s\rangle^{ON} + \langle s\rangle^{OFF}\right) + \left(\langle r\rangle^{ON} + \langle r\rangle^{OFF}\right)\langle ps\rangle^{OFF} \\
&- \left(\langle r\rangle^{ON} + \langle r\rangle^{OFF}\right)\langle p\rangle^{OFF}\left(\langle s\rangle^{ON} + \langle s\rangle^{OFF}\right) + \left(\langle r\rangle^{ON} + \langle r\rangle^{OFF}\right)\left(\langle p\rangle^{ON} + \langle p\rangle^{OFF}\right)\langle s\rangle^{ON} \\
&- P^{ON}\left(\langle r\rangle^{ON} + \langle r\rangle^{OFF}\right)\left(\langle p\rangle^{ON} + \langle p\rangle^{OFF}\right)\left(\langle s\rangle^{ON} + \langle s\rangle^{OFF}\right)\bigg] \\
&+ k_+\bigg[\langle rs\rangle^{ON} - \Big\{2\langle rs\rangle^{ON}\left(\langle r\rangle^{ON} + \langle r\rangle^{OFF}\right) - \left(\langle r\rangle^{ON} + \langle r\rangle^{OFF}\right)^2\langle s\rangle^{ON} \\
&- 2\langle r\rangle^{ON}\left(\langle r\rangle^{ON} + \langle r\rangle^{OFF}\right)\left(\langle s\rangle^{ON} + \langle s\rangle^{OFF}\right) + \langle r^2\rangle^{ON}\left(\langle s\rangle^{ON} + \langle s\rangle^{OFF}\right) \\
&+ \left(\langle r\rangle^{ON} + \langle r\rangle^{OFF}\right)^2\left(\langle s\rangle^{ON} + \langle s\rangle^{ON}\right)P^{ON}\Big\} \\
&- \Big\{2\langle rs\rangle^{ON}\left(\langle s\rangle^{ON} + \langle s\rangle^{OFF}\right) - \langle r\rangle^{ON}\left(\langle s\rangle^{ON} + \langle s\rangle^{OFF}\right)^2 + \left(\langle r\rangle^{ON} + \langle r\rangle^{OFF}\right)\langle s^2\rangle^{ON} \\
&- 2\left(\langle r\rangle^{ON} + \langle r\rangle^{OFF}\right)\left(\langle s\rangle^{ON} + \langle s\rangle^{OFF}\right)\langle s\rangle^{ON} \\
&+ \left(\langle r\rangle^{ON} + \langle r\rangle^{OFF}\right)\left(\langle s\rangle^{ON} + \langle s\rangle^{OFF}\right)^2 P^{ON}\Big\}\bigg] \\
&+ k_-\left[\langle rc\rangle^{ON} + \langle sc\rangle^{ON} + \langle c\rangle^{ON}\right] + (1-\alpha)\gamma\langle rc\rangle^{ON} \qquad \text{(S 6.9)}
\end{aligned}
$$

**x)**

$$
\begin{aligned}
\frac{\partial \langle rs\rangle^{OFF}}{\partial t} &= k_r\langle s\rangle^{OFF} - (g_r + g_s)\langle rs\rangle^{OFF} + k_s^0\langle r\rangle^{OFF} + k_{deact}\langle rs\rangle^{ON} \\
&- k_{act}\bigg[\langle rp\rangle^{OFF}\left(\langle s\rangle^{ON} + \langle s\rangle^{OFF}\right) + \langle rs\rangle^{OFF}\left(\langle p\rangle^{ON} + \langle p\rangle^{OFF}\right) \\
&- \langle r\rangle^{OFF}\left(\langle p\rangle^{ON} + \langle p\rangle^{OFF}\right)\left(\langle s\rangle^{ON} + \langle s\rangle^{OFF}\right) + \left(\langle r\rangle^{ON} + \langle r\rangle^{OFF}\right)\langle ps\rangle^{OFF} \\
&- \left(\langle r\rangle^{ON} + \langle r\rangle^{OFF}\right)\langle p\rangle^{OFF}\left(\langle s\rangle^{ON} + \langle s\rangle^{OFF}\right) + \left(\langle r\rangle^{ON} + \langle r\rangle^{OFF}\right)\left(\langle p\rangle^{ON} + \langle p\rangle^{OFF}\right)\langle s\rangle^{OFF} \\
&- P^{ON}\left(\langle r\rangle^{ON} + \langle r\rangle^{OFF}\right)\left(\langle p\rangle^{ON} + \langle p\rangle^{OFF}\right)\left(\langle s\rangle^{ON} + \langle s\rangle^{OFF}\right)\bigg] \\
&+ k_+\bigg[\langle rs\rangle^{OFF} - \Big\{2\langle rs\rangle^{OFF}\left(\langle r\rangle^{ON} + \langle r\rangle^{OFF}\right) + \left(\langle r\rangle^{ON} + \langle r\rangle^{OFF}\right)^2\langle s\rangle^{ON} \\
&- 2\langle r\rangle^{OFF}\left(\langle r\rangle^{ON} + \langle r\rangle^{OFF}\right)\left(\langle s\rangle^{ON} + \langle s\rangle^{OFF}\right) + \langle r^2\rangle^{OFF}\left(\langle s\rangle^{ON} + \langle s\rangle^{OFF}\right) \\
&- \left(\langle r\rangle^{ON} + \langle r\rangle^{OFF}\right)^2\left(\langle s\rangle^{ON} + \langle s\rangle^{OFF}\right)P^{ON}\Big\} \\
&- \Big\{2\langle rs\rangle^{OFF}\left(\langle s\rangle^{ON} + \langle s\rangle^{OFF}\right) + \langle r\rangle^{ON}\left(\langle s\rangle^{ON} + \langle s\rangle^{OFF}\right)^2 + \left(\langle r\rangle^{ON} + \langle r\rangle^{OFF}\right)\langle s^2\rangle^{OFF} \\
&- \left(\langle r\rangle^{ON} + \langle r\rangle^{OFF}\right)\left(\langle s\rangle^{ON} + \langle s\rangle^{OFF}\right)^2 P^{ON} \\
&- 2\left(\langle r\rangle^{ON} + \langle r\rangle^{OFF}\right)\left(\langle s\rangle^{ON} + \langle s\rangle^{OFF}\right)\langle s\rangle^{OFF}\Big\}\bigg] \\
&+ k_-\left[\langle rc\rangle^{OFF} + \langle sc\rangle^{OFF} + \langle c\rangle^{OFF}\right] + (1-\alpha)\gamma\langle rc\rangle^{OFF} \qquad \text{(S 6.10)}
\end{aligned}
$$

$$\textbf{xi)}\quad \frac{\partial \langle rc\rangle^{ON}}{\partial t} = k_r\langle c\rangle^{ON} - (g_r + k_{deact} + \gamma)\langle rc\rangle^{ON} + k_-\left[\langle c^2\rangle^{ON} - \langle c\rangle^{ON} - \langle rc\rangle^{ON}\right]$$
$$+ k_{act}\Big[\langle rp\rangle^{OFF}\left(\langle c\rangle^{ON} + \langle c\rangle^{OFF}\right) + \langle rc\rangle^{OFF}\left(\langle p\rangle^{ON} + \langle p\rangle^{OFF}\right) + \left(\langle r\rangle^{ON} + \langle r\rangle^{OFF}\right)\langle pc\rangle^{OFF}$$
$$- \langle r\rangle^{OFF}\left(\langle p\rangle^{ON} + \langle p\rangle^{OFF}\right)\left(\langle c\rangle^{ON} + \langle c\rangle^{OFF}\right) + \left(\langle r\rangle^{ON} + \langle r\rangle^{OFF}\right)\langle p\rangle^{ON}\left(\langle c\rangle^{ON} + \langle c\rangle^{OFF}\right)$$
$$- \left(\langle r\rangle^{ON} + \langle r\rangle^{OFF}\right)\left(\langle p\rangle^{ON} + \langle p\rangle^{OFF}\right)\langle c\rangle^{OFF}$$
$$- P^{ON}\left(\langle r\rangle^{ON} + \langle r\rangle^{OFF}\right)\left(\langle p\rangle^{ON} + \langle p\rangle^{OFF}\right)\left(\langle c\rangle^{ON} + \langle c\rangle^{OFF}\right)\Big]$$
$$+ k_+\Big[\Big\{2\langle rs\rangle^{ON}\left(\langle r\rangle^{ON} + \langle r\rangle^{OFF}\right) - \left(\langle r\rangle^{ON} + \langle r\rangle^{OFF}\right)^2\langle s\rangle^{ON}$$
$$- 2\langle r\rangle^{ON}\left(\langle r\rangle^{ON} + \langle r\rangle^{OFF}\right)\left(\langle s\rangle^{ON} + \langle s\rangle^{OFF}\right) + \langle r^2\rangle^{ON}\left(\langle s\rangle^{ON} + \langle s\rangle^{OFF}\right)$$
$$+ \left(\langle r\rangle^{ON} + \langle r\rangle^{OFF}\right)^2\left(\langle s\rangle^{ON} + \langle s\rangle^{ON}\right)P^{ON}\Big\}$$
$$- \Big\{\langle rs\rangle^{ON}\left(\langle c\rangle^{ON} + \langle c\rangle^{OFF}\right) + \langle rc\rangle^{ON}\left(\langle s\rangle^{ON} + \langle s\rangle^{OFF}\right)$$
$$- \langle r\rangle^{ON}\left(\langle s\rangle^{ON} + \langle s\rangle^{OFF}\right)\left(\langle c\rangle^{ON} + \langle c\rangle^{OFF}\right) + \left(\langle r\rangle^{ON} + \langle r\rangle^{OFF}\right)\langle sc\rangle^{ON}$$
$$- \left(\langle r\rangle^{ON} + \langle r\rangle^{OFF}\right)\langle s\rangle^{ON}\left(\langle c\rangle^{ON} + \langle c\rangle^{OFF}\right)$$
$$- \left(\langle r\rangle^{ON} + \langle r\rangle^{OFF}\right)\left(\langle s\rangle^{ON} + \langle s\rangle^{OFF}\right)\langle c\rangle^{ON}$$
$$+ P^{ON}\left(\langle r\rangle^{ON} + \langle r\rangle^{OFF}\right)\left(\langle s\rangle^{ON} + \langle s\rangle^{OFF}\right)\left(\langle c\rangle^{ON} + \langle c\rangle^{OFF}\right)\Big\} - \langle rs\rangle^{ON}\Big] \tag{S 6.11}$$

$$\textbf{xii)}\quad \frac{\partial \langle rc\rangle^{OFF}}{\partial t} = k_r\langle c\rangle^{OFF} - (g_r + \gamma)\langle rc\rangle^{OFF} + k_{deact}\langle rc\rangle^{ON} + k_-\left[\langle c^2\rangle^{OFF} - \langle c\rangle^{OFF} - \langle rc\rangle^{OFF}\right]$$
$$- k_{act}\Big[\langle rp\rangle^{OFF}\left(\langle c\rangle^{ON} + \langle c\rangle^{OFF}\right) + \langle rc\rangle^{OFF}\left(\langle p\rangle^{ON} + \langle p\rangle^{OFF}\right) + \left(\langle r\rangle^{ON} + \langle r\rangle^{OFF}\right)\langle pc\rangle^{OFF}$$
$$- \langle r\rangle^{OFF}\left(\langle p\rangle^{ON} + \langle p\rangle^{OFF}\right)\left(\langle c\rangle^{ON} + \langle c\rangle^{OFF}\right) + \left(\langle r\rangle^{ON} + \langle r\rangle^{OFF}\right)\langle p\rangle^{ON}\left(\langle c\rangle^{ON} + \langle c\rangle^{OFF}\right)$$
$$- \left(\langle r\rangle^{ON} + \langle r\rangle^{OFF}\right)\left(\langle p\rangle^{ON} + \langle p\rangle^{OFF}\right)\langle c\rangle^{OFF}$$
$$- P^{ON}\left(\langle r\rangle^{ON} + \langle r\rangle^{OFF}\right)\left(\langle p\rangle^{ON} + \langle p\rangle^{OFF}\right)\left(\langle c\rangle^{ON} + \langle c\rangle^{OFF}\right)\Big]$$
$$+ k_+\Big[\Big\{2\langle rs\rangle^{OFF}\left(\langle r\rangle^{ON} + \langle r\rangle^{OFF}\right) + \left(\langle r\rangle^{ON} + \langle r\rangle^{OFF}\right)^2\langle s\rangle^{ON}$$
$$- 2\langle r\rangle^{OFF}\left(\langle r\rangle^{ON} + \langle r\rangle^{OFF}\right)\left(\langle s\rangle^{ON} + \langle s\rangle^{OFF}\right) + \langle r^2\rangle^{OFF}\left(\langle s\rangle^{ON} + \langle s\rangle^{OFF}\right)$$
$$- \left(\langle r\rangle^{ON} + \langle r\rangle^{OFF}\right)^2\left(\langle s\rangle^{ON} + \langle s\rangle^{OFF}\right)P^{ON}\Big\}$$
$$- \Big\{\langle rs\rangle^{OFF}\left(\langle c\rangle^{ON} + \langle c\rangle^{OFF}\right) + \langle rc\rangle^{OFF}\left(\langle s\rangle^{ON} + \langle s\rangle^{OFF}\right)$$
$$- \langle r\rangle^{OFF}\left(\langle s\rangle^{ON} + \langle s\rangle^{OFF}\right)\left(\langle c\rangle^{ON} + \langle c\rangle^{OFF}\right) + \left(\langle r\rangle^{ON} + \langle r\rangle^{OFF}\right)\langle sc\rangle^{OFF}$$
$$- \left(\langle r\rangle^{ON} + \langle r\rangle^{OFF}\right)\langle s\rangle^{OFF}\left(\langle c\rangle^{ON} + \langle c\rangle^{OFF}\right)$$
$$+ \left(\langle r\rangle^{ON} + \langle r\rangle^{OFF}\right)\left(\langle s\rangle^{ON} + \langle s\rangle^{OFF}\right)\langle c\rangle^{ON}$$
$$- P^{ON}\left(\langle r\rangle^{ON} + \langle r\rangle^{OFF}\right)\left(\langle s\rangle^{ON} + \langle s\rangle^{OFF}\right)\left(\langle c\rangle^{ON} + \langle c\rangle^{OFF}\right)\Big\} - \langle rs\rangle^{OFF}\Big] \tag{S 6.12}$$

**xiii)** $$\frac{\partial \langle sc\rangle^{ON}}{\partial t} = k_s\langle c\rangle^{ON} - (g_s+\alpha\gamma)\langle sc\rangle^{ON} - k_{deact}\langle sc\rangle^{ON}$$
$$+ k_{act}\Big[\langle ps\rangle^{OFF}\left(\langle c\rangle^{ON}+\langle c\rangle^{OFF}\right) + \langle pc\rangle^{OFF}\left(\langle s\rangle^{ON}+\langle s\rangle^{OFF}\right)$$
$$- \langle p\rangle^{OFF}\left(\langle s\rangle^{ON}+\langle s\rangle^{OFF}\right)\left(\langle c\rangle^{ON}+\langle c\rangle^{OFF}\right) + \left(\langle p\rangle^{ON}+\langle p\rangle^{OFF}\right)\langle sc\rangle^{OFF}$$
$$- \left(\langle p\rangle^{ON}+\langle p\rangle^{OFF}\right)\langle s\rangle^{OFF}\left(\langle c\rangle^{ON}+\langle c\rangle^{OFF}\right)$$
$$+ \left(\langle p\rangle^{ON}+\langle p\rangle^{OFF}\right)\left(\langle s\rangle^{ON}+\langle s\rangle^{OFF}\right)\langle c\rangle^{ON}$$
$$- P^{ON}\left(\langle p\rangle^{ON}+\langle p\rangle^{OFF}\right)\left(\langle s\rangle^{ON}+\langle s\rangle^{OFF}\right)\left(\langle c\rangle^{ON}+\langle c\rangle^{OFF}\right)\Big]$$
$$+ k_+\Big[\Big\{2\langle rs\rangle^{ON}\left(\langle s\rangle^{ON}+\langle s\rangle^{OFF}\right) - \langle r\rangle^{ON}\left(\langle s\rangle^{ON}+\langle s\rangle^{OFF}\right)^2$$
$$+ \left(\langle r\rangle^{ON}+\langle r\rangle^{OFF}\right)\langle s^2\rangle^{ON} - 2\left(\langle r\rangle^{ON}+\langle r\rangle^{OFF}\right)\left(\langle s\rangle^{ON}+\langle s\rangle^{OFF}\right)\langle s\rangle^{ON}$$
$$+ \left(\langle r\rangle^{ON}+\langle r\rangle^{OFF}\right)\left(\langle s\rangle^{ON}+\langle s\rangle^{OFF}\right)^2 P^{ON}\Big\}$$
$$- \Big\{\langle rs\rangle^{ON}\left(\langle c\rangle^{ON}+\langle c\rangle^{OFF}\right) + \langle rc\rangle^{ON}\left(\langle s\rangle^{ON}+\langle s\rangle^{OFF}\right)$$
$$- \langle r\rangle^{ON}\left(\langle s\rangle^{ON}+\langle s\rangle^{OFF}\right)\left(\langle c\rangle^{ON}+\langle c\rangle^{OFF}\right) + \left(\langle r\rangle^{ON}+\langle r\rangle^{OFF}\right)\langle sc\rangle^{ON}$$
$$- \left(\langle r\rangle^{ON}+\langle r\rangle^{OFF}\right)\langle s\rangle^{ON}\left(\langle c\rangle^{ON}+\langle c\rangle^{OFF}\right)$$
$$- \left(\langle r\rangle^{ON}+\langle r\rangle^{OFF}\right)\left(\langle s\rangle^{ON}+\langle s\rangle^{OFF}\right)\langle c\rangle^{ON}$$
$$+ P^{ON}\left(\langle r\rangle^{ON}+\langle r\rangle^{OFF}\right)\left(\langle s\rangle^{ON}+\langle s\rangle^{OFF}\right)\left(\langle c\rangle^{ON}+\langle c\rangle^{OFF}\right)\Big\} - \langle rs\rangle^{ON}\Big]$$
$$+ \left[k_- + (1-\alpha)\gamma\right]\left[\langle c^2\rangle^{ON} - \langle c\rangle^{ON} - \langle sc\rangle^{ON}\right] \tag{S 6.13}$$

**xiv)** $$\frac{\partial \langle sc\rangle^{OFF}}{\partial t} = k_s^0\langle c\rangle^{OFF} - (g_s+\alpha\gamma)\langle sc\rangle^{OFF} + k_{deact}\langle sc\rangle^{OFF}$$
$$- k_{act}\Big[\langle ps\rangle^{OFF}\left(\langle c\rangle^{ON}+\langle c\rangle^{OFF}\right) + \langle pc\rangle^{OFF}\left(\langle s\rangle^{ON}+\langle s\rangle^{OFF}\right)$$
$$- \langle p\rangle^{OFF}\left(\langle s\rangle^{ON}+\langle s\rangle^{OFF}\right)\left(\langle c\rangle^{ON}+\langle c\rangle^{OFF}\right) + \left(\langle p\rangle^{ON}+\langle p\rangle^{OFF}\right)\langle sc\rangle^{OFF}$$
$$- \left(\langle p\rangle^{ON}+\langle p\rangle^{OFF}\right)\langle s\rangle^{OFF}\left(\langle c\rangle^{ON}+\langle c\rangle^{OFF}\right)$$
$$+ \left(\langle p\rangle^{ON}+\langle p\rangle^{OFF}\right)\left(\langle s\rangle^{ON}+\langle s\rangle^{OFF}\right)\langle c\rangle^{ON}$$
$$- P^{ON}\left(\langle p\rangle^{ON}+\langle p\rangle^{OFF}\right)\left(\langle s\rangle^{ON}+\langle s\rangle^{OFF}\right)\left(\langle c\rangle^{ON}+\langle c\rangle^{OFF}\right)\Big]$$
$$+ k_+\Big[\Big\{2\langle rs\rangle^{OFF}\left(\langle s\rangle^{ON}+\langle s\rangle^{OFF}\right) + \langle r\rangle^{ON}\left(\langle s\rangle^{ON}+\langle s\rangle^{OFF}\right)^2$$
$$+ \left(\langle r\rangle^{ON}+\langle r\rangle^{OFF}\right)\langle s^2\rangle^{OFF} - 2\left(\langle r\rangle^{ON}+\langle r\rangle^{OFF}\right)\left(\langle s\rangle^{ON}+\langle s\rangle^{OFF}\right)\langle s\rangle^{OFF}$$
$$- \left(\langle r\rangle^{ON}+\langle r\rangle^{OFF}\right)\left(\langle s\rangle^{ON}+\langle s\rangle^{OFF}\right)^2 P^{ON}\Big\}$$
$$- \Big\{\langle rs\rangle^{OFF}\left(\langle c\rangle^{ON}+\langle c\rangle^{OFF}\right) + \langle rc\rangle^{OFF}\left(\langle s\rangle^{ON}+\langle s\rangle^{OFF}\right)$$
$$- \langle r\rangle^{OFF}\left(\langle s\rangle^{ON}+\langle s\rangle^{OFF}\right)\left(\langle c\rangle^{ON}+\langle c\rangle^{OFF}\right) + \left(\langle r\rangle^{ON}+\langle r\rangle^{OFF}\right)\langle sc\rangle^{OFF}$$
$$- \left(\langle r\rangle^{ON}+\langle r\rangle^{OFF}\right)\langle s\rangle^{OFF}\left(\langle c\rangle^{ON}+\langle c\rangle^{OFF}\right)$$
$$+ \left(\langle r\rangle^{ON}+\langle r\rangle^{OFF}\right)\left(\langle s\rangle^{ON}+\langle s\rangle^{OFF}\right)\langle c\rangle^{ON}$$
$$- P^{ON}\left(\langle r\rangle^{ON}+\langle r\rangle^{OFF}\right)\left(\langle s\rangle^{ON}+\langle s\rangle^{OFF}\right)\left(\langle c\rangle^{ON}+\langle c\rangle^{OFF}\right)\Big\} - \langle rs\rangle^{OFF}\Big]$$
$$+ \left[k_- + (1-\alpha)\gamma\right]\left[\langle c^2\rangle^{OFF} - \langle c\rangle^{OFF} - \langle sc\rangle^{OFF}\right] \tag{S 6.14}$$

**xv)** $$\frac{\partial \langle rp \rangle^{ON}}{\partial t} = k_r \langle p \rangle^{ON} - (g_r + g_p) \langle rp \rangle^{ON} + k_p \langle r^2 \rangle^{ON} - k_{deact} \langle rp \rangle^{ON} + k_- \langle pc \rangle^{ON}$$
$$+ k_{act} \Big[ \Big\{ 2\langle rp \rangle^{OFF} \left( \langle p \rangle^{ON} + \langle p \rangle^{OFF} \right) + \langle r \rangle^{ON} \left( \langle p \rangle^{ON} + \langle p \rangle^{OFF} \right)^2$$
$$+ \left( \langle r \rangle^{ON} + \langle r \rangle^{OFF} \right) \langle p^2 \rangle^{OFF} - 2 \left( \langle r \rangle^{ON} + \langle r \rangle^{OFF} \right) \left( \langle p \rangle^{ON} + \langle p \rangle^{OFF} \right) \langle p \rangle^{OFF}$$
$$- \left( \langle r \rangle^{ON} + \langle r \rangle^{OFF} \right) \left( \langle p \rangle^{ON} + \langle p \rangle^{OFF} \right)^2 P^{ON} \Big\} - \langle rp \rangle^{OFF} \Big]$$
$$- k_+ \Big[ \langle rp \rangle^{ON} \left( \langle s \rangle^{ON} + \langle s \rangle^{OFF} \right) + \langle rs \rangle^{ON} \left( \langle p \rangle^{ON} + \langle p \rangle^{OFF} \right)$$
$$- \langle r \rangle^{ON} \left( \langle p \rangle^{ON} + \langle p \rangle^{OFF} \right) \left( \langle s \rangle^{ON} + \langle s \rangle^{OFF} \right) + \left( \langle r \rangle^{ON} + \langle r \rangle^{OFF} \right) \langle ps \rangle^{ON}$$
$$- \left( \langle r \rangle^{ON} + \langle r \rangle^{OFF} \right) \langle p \rangle^{ON} \left( \langle s \rangle^{ON} + \langle s \rangle^{OFF} \right)$$
$$- \left( \langle r \rangle^{ON} + \langle r \rangle^{OFF} \right) \left( \langle p \rangle^{ON} + \langle p \rangle^{OFF} \right) \langle s \rangle^{ON}$$
$$+ P^{ON} \left( \langle r \rangle^{ON} + \langle r \rangle^{OFF} \right) \left( \langle p \rangle^{ON} + \langle p \rangle^{OFF} \right) \left( \langle s \rangle^{ON} + \langle s \rangle^{OFF} \right) \Big] \tag{S 6.15}$$

**xvi)** $$\frac{\partial \langle rp \rangle^{OFF}}{\partial t} = k_r \langle p \rangle^{OFF} - (g_r + g_p) \langle rp \rangle^{OFF} + k_p \langle r^2 \rangle^{OFF}$$
$$- k_{act} \Big[ 2\langle rp \rangle^{OFF} \left( \langle p \rangle^{ON} + \langle p \rangle^{OFF} \right) + \langle r \rangle^{ON} \left( \langle p \rangle^{ON} + \langle p \rangle^{OFF} \right)^2$$
$$+ \left( \langle r \rangle^{ON} + \langle r \rangle^{OFF} \right) \langle p^2 \rangle^{OFF} - 2 \left( \langle r \rangle^{ON} + \langle r \rangle^{OFF} \right) \left( \langle p \rangle^{ON} + \langle p \rangle^{OFF} \right) \langle p \rangle^{OFF}$$
$$- \left( \langle r \rangle^{ON} + \langle r \rangle^{OFF} \right) \left( \langle p \rangle^{ON} + \langle p \rangle^{OFF} \right)^2 P^{ON} \Big]$$
$$+ k_{deact} \left[ \langle rp \rangle^{ON} + \langle r \rangle^{ON} \right] + k_- \langle pc \rangle^{OFF}$$
$$- k_+ \Big[ \langle rp \rangle^{OFF} \left( \langle s \rangle^{ON} + \langle s \rangle^{OFF} \right) + \langle rs \rangle^{OFF} \left( \langle p \rangle^{ON} + \langle p \rangle^{OFF} \right)$$
$$- \langle r \rangle^{OFF} \left( \langle p \rangle^{ON} + \langle p \rangle^{OFF} \right) \left( \langle s \rangle^{ON} + \langle s \rangle^{OFF} \right) + \left( \langle r \rangle^{ON} + \langle r \rangle^{OFF} \right) \langle ps \rangle^{OFF}$$
$$- \left( \langle r \rangle^{ON} + \langle r \rangle^{OFF} \right) \langle p \rangle^{OFF} \left( \langle s \rangle^{ON} + \langle s \rangle^{OFF} \right)$$
$$+ \left( \langle r \rangle^{ON} + \langle r \rangle^{OFF} \right) \left( \langle p \rangle^{ON} + \langle p \rangle^{OFF} \right) \langle s \rangle^{ON}$$
$$- P^{ON} \left( \langle r \rangle^{ON} + \langle r \rangle^{OFF} \right) \left( \langle p \rangle^{ON} + \langle p \rangle^{OFF} \right) \left( \langle s \rangle^{ON} + \langle s \rangle^{OFF} \right) \Big] \tag{S 6.16}$$

**xvii)** $$\frac{\partial \langle ps \rangle^{ON}}{\partial t} = k_p \langle rs \rangle^{ON} - (g_p + g_s) \langle ps \rangle^{ON} + k_s \langle p \rangle^{ON} - k_{deact} \langle ps \rangle^{ON}$$
$$+ k_{act} \Big[ \Big\{ 2\langle ps \rangle^{OFF} \left( \langle p \rangle^{ON} + \langle p \rangle^{OFF} \right) + \left( \langle p \rangle^{ON} + \langle p \rangle^{OFF} \right)^2 \langle s \rangle^{ON}$$
$$- 2\langle p \rangle^{OFF} \left( \langle p \rangle^{ON} + \langle p \rangle^{OFF} \right) \left( \langle s \rangle^{ON} + \langle s \rangle^{OFF} \right) + \langle p^2 \rangle^{OFF} \left( \langle s \rangle^{ON} + \langle s \rangle^{OFF} \right)$$
$$- \left( \langle p \rangle^{ON} + \langle p \rangle^{OFF} \right)^2 \left( \langle s \rangle^{ON} + \langle s \rangle^{OFF} \right) P^{ON} \Big\} - \langle ps \rangle^{OFF} \Big]$$
$$- k_+ \Big[ \langle rp \rangle^{ON} \left( \langle s \rangle^{ON} + \langle s \rangle^{OFF} \right) + \langle rs \rangle^{ON} \left( \langle p \rangle^{ON} + \langle p \rangle^{OFF} \right)$$
$$- \langle r \rangle^{ON} \left( \langle p \rangle^{ON} + \langle p \rangle^{OFF} \right) \left( \langle s \rangle^{ON} + \langle s \rangle^{OFF} \right) + \left( \langle r \rangle^{ON} + \langle r \rangle^{OFF} \right) \langle ps \rangle^{ON}$$
$$- \left( \langle r \rangle^{ON} + \langle r \rangle^{OFF} \right) \langle p \rangle^{ON} \left( \langle s \rangle^{ON} + \langle s \rangle^{OFF} \right)$$
$$- \left( \langle r \rangle^{ON} + \langle r \rangle^{OFF} \right) \left( \langle p \rangle^{ON} + \langle p \rangle^{OFF} \right) \langle s \rangle^{ON}$$
$$+ P^{ON} \left( \langle r \rangle^{ON} + \langle r \rangle^{OFF} \right) \left( \langle p \rangle^{ON} + \langle p \rangle^{OFF} \right) \left( \langle s \rangle^{ON} + \langle s \rangle^{OFF} \right) \Big]$$
$$+ \Big[ k_- + (1-\alpha)\gamma \Big] \langle pc \rangle^{ON} \tag{S 6.17}$$

$$
\textbf{xviii)}\quad \frac{\partial \langle ps\rangle^{OFF}}{\partial t} = k_p\langle rs\rangle^{OFF} - (g_p+g_s)\langle ps\rangle^{OFF} + k_s^0\langle p\rangle^{OFF}
$$
$$
+ k_{deact}\left[\langle s\rangle^{ON} + \langle ps\rangle^{ON}\right] + \left[k_- + (1-\alpha)\gamma\right]\langle pc\rangle^{OFF}
$$
$$
- k_{act}\Big[2\langle ps\rangle^{OFF}\left(\langle p\rangle^{ON}+\langle p\rangle^{OFF}\right) + \left(\langle p\rangle^{ON}+\langle p\rangle^{OFF}\right)^2\langle s\rangle^{ON}
$$
$$
- 2\langle p\rangle^{OFF}\left(\langle p\rangle^{ON}+\langle p\rangle^{OFF}\right)\left(\langle s\rangle^{ON}+\langle s\rangle^{OFF}\right) + \langle p^2\rangle^{OFF}\left(\langle s\rangle^{ON}+\langle s\rangle^{OFF}\right)
$$
$$
- \left(\langle p\rangle^{ON}+\langle p\rangle^{OFF}\right)^2\left(\langle s\rangle^{ON}+\langle s\rangle^{OFF}\right)P^{ON}\Big]
$$
$$
- k_+\Big[\langle rp\rangle^{OFF}\left(\langle s\rangle^{ON}+\langle s\rangle^{OFF}\right) + \langle rs\rangle^{OFF}\left(\langle p\rangle^{ON}+\langle p\rangle^{OFF}\right)
$$
$$
- \langle r\rangle^{OFF}\left(\langle p\rangle^{ON}+\langle p\rangle^{OFF}\right)\left(\langle s\rangle^{ON}+\langle s\rangle^{OFF}\right) + \left(\langle r\rangle^{ON}+\langle r\rangle^{OFF}\right)\langle ps\rangle^{OFF}
$$
$$
- \left(\langle r\rangle^{ON}+\langle r\rangle^{OFF}\right)\langle p\rangle^{OFF}\left(\langle s\rangle^{ON}+\langle s\rangle^{OFF}\right)
$$
$$
+ \left(\langle r\rangle^{ON}+\langle r\rangle^{OFF}\right)\left(\langle p\rangle^{ON}+\langle p\rangle^{OFF}\right)\langle s\rangle^{ON}
$$
$$
- P^{ON}\left(\langle r\rangle^{ON}+\langle r\rangle^{OFF}\right)\left(\langle p\rangle^{ON}+\langle p\rangle^{OFF}\right)\left(\langle s\rangle^{ON}+\langle s\rangle^{OFF}\right)\Big] \tag{S 6.18}
$$

$$
\textbf{xix)}\quad \frac{\partial \langle pc\rangle^{ON}}{\partial t} = k_p\langle rc\rangle^{ON} - \left[g_p + k_- + \gamma\right]\langle pc\rangle^{ON} - k_{deact}\langle pc\rangle^{ON}
$$
$$
+ k_{act}\Big[\Big\{2\langle pc\rangle^{OFF}\left(\langle p\rangle^{ON}+\langle p\rangle^{OFF}\right) + \left(\langle p\rangle^{ON}+\langle p\rangle^{OFF}\right)^2\langle c\rangle^{ON}
$$
$$
- 2\langle p\rangle^{OFF}\left(\langle p\rangle^{ON}+\langle p\rangle^{OFF}\right)\left(\langle c\rangle^{ON}+\langle c\rangle^{OFF}\right) + \langle p^2\rangle^{OFF}\left(\langle c\rangle^{ON}+\langle c\rangle^{OFF}\right)
$$
$$
- \left(\langle p\rangle^{ON}+\langle p\rangle^{OFF}\right)^2\left(\langle c\rangle^{ON}+\langle c\rangle^{OFF}\right)P^{ON}\Big\} - \langle pc\rangle^{OFF}\Big]
$$
$$
+ k_+\Big[\langle rp\rangle^{ON}\left(\langle s\rangle^{ON}+\langle s\rangle^{OFF}\right) + \langle rs\rangle^{ON}\left(\langle p\rangle^{ON}+\langle p\rangle^{OFF}\right)
$$
$$
- \langle r\rangle^{ON}\left(\langle p\rangle^{ON}+\langle p\rangle^{OFF}\right)\left(\langle s\rangle^{ON}+\langle s\rangle^{OFF}\right)
$$
$$
+ \left(\langle r\rangle^{ON}+\langle r\rangle^{OFF}\right)\langle ps\rangle^{ON} - \left(\langle r\rangle^{ON}+\langle r\rangle^{OFF}\right)\langle p\rangle^{ON}\left(\langle s\rangle^{ON}+\langle s\rangle^{OFF}\right)
$$
$$
- \left(\langle r\rangle^{ON}+\langle r\rangle^{OFF}\right)\left(\langle p\rangle^{ON}+\langle p\rangle^{OFF}\right)\langle s\rangle^{ON}
$$
$$
+ P^{ON}\left(\langle r\rangle^{ON}+\langle r\rangle^{OFF}\right)\left(\langle p\rangle^{ON}+\langle p\rangle^{OFF}\right)\left(\langle s\rangle^{ON}+\langle s\rangle^{OFF}\right)\Big] \tag{S 6.19}
$$

$$
\textbf{xx)}\quad \frac{\partial \langle pc\rangle^{OFF}}{\partial t} = k_p\langle rc\rangle^{OFF} - \left[g_p + k_- + \gamma\right]\langle pc\rangle^{OFF} + k_{deact}\left[\langle pc\rangle^{ON} + \langle c\rangle^{ON}\right]
$$
$$
- k_{act}\Big[2\langle pc\rangle^{OFF}\left(\langle p\rangle^{ON}+\langle p\rangle^{OFF}\right) + \left(\langle p\rangle^{ON}+\langle p\rangle^{OFF}\right)^2\langle c\rangle^{ON}
$$
$$
- 2\langle p\rangle^{OFF}\left(\langle p\rangle^{ON}+\langle p\rangle^{OFF}\right)\left(\langle c\rangle^{ON}+\langle c\rangle^{OFF}\right)
$$
$$
+ \langle p^2\rangle^{OFF}\left(\langle c\rangle^{ON}+\langle c\rangle^{OFF}\right)
$$
$$
- \left(\langle p\rangle^{ON}+\langle p\rangle^{OFF}\right)^2\left(\langle c\rangle^{ON}+\langle c\rangle^{OFF}\right)P^{ON}\Big]
$$
$$
+ k_+\Big[\langle rp\rangle^{OFF}\left(\langle s\rangle^{ON}+\langle s\rangle^{OFF}\right) + \langle rs\rangle^{OFF}\left(\langle p\rangle^{ON}+\langle p\rangle^{OFF}\right)
$$
$$
- \langle r\rangle^{OFF}\left(\langle p\rangle^{ON}+\langle p\rangle^{OFF}\right)\left(\langle s\rangle^{ON}+\langle s\rangle^{OFF}\right)
$$
$$
+ \left(\langle r\rangle^{ON}+\langle r\rangle^{OFF}\right)\langle ps\rangle^{OFF}
$$
$$
- \left(\langle r\rangle^{ON}+\langle r\rangle^{OFF}\right)\langle p\rangle^{OFF}\left(\langle s\rangle^{ON}+\langle s\rangle^{OFF}\right)
$$
$$
+ \left(\langle r\rangle^{ON}+\langle r\rangle^{OFF}\right)\left(\langle p\rangle^{ON}+\langle p\rangle^{OFF}\right)\langle s\rangle^{ON}
$$
$$
- P^{ON}\left(\langle r\rangle^{ON}+\langle r\rangle^{OFF}\right)\left(\langle p\rangle^{ON}+\langle p\rangle^{OFF}\right)\left(\langle s\rangle^{ON}+\langle s\rangle^{OFF}\right)\Big] \tag{S 6.20}
$$

Therefore equations (S 2.1)-(S 2.9) along with equations (S 6.1)-(S 6.20) form a closed system of differential equations, which is called the second order equations (**SOE**), since all the second order moments are linear combination of the variables of this system of equations (for example, $< r^2 >=< r^2 >^{ON} + < r^2 >^{OFF}$ etc.)
